# eDNA metabarcoding outperforms traditional fisheries sampling and reveals fine-scale heterogeneity in a temperate freshwater lake

**DOI:** 10.1101/2020.10.20.347203

**Authors:** Rebecca R Gehri, Wesley A. Larson, Kristen Gruenthal, Nicholas Sard, Yue Shi

## Abstract

Understanding biodiversity in aquatic systems is critical to ecological research and conservation efforts, but accurately measuring species richness using traditional methods can be challenging. Environmental DNA (eDNA) metabarcoding, which uses high-throughput sequencing and universal primers to amplify DNA from multiple species present in an environmental sample, has shown great promise for augmenting results from traditional sampling to characterize fish communities in aquatic systems. Few studies, however, have compared exhaustive traditional sampling with eDNA metabarcoding of corresponding water samples at a small spatial scale. We intensively sampled Boardman Lake (137 ha) in Michigan, USA from May to June in 2019 using gill and fyke nets and paired each net set with lake water samples collected in triplicate. We analyzed water samples using eDNA metabarcoding with 12S and 16S fish-specific primers and compared estimates of fish diversity among methods. In total, we set 60 nets and analyzed 180 1 L lake water samples. We captured a total of 12 fish species in our traditional gear and detected 40 taxa in the eDNA water samples, which included all the species observed in nets. The 12S and 16S assays detected a comparable number of taxa, but taxonomic resolution varied between the two genes. In our traditional gear, there was a clear difference in the species selectivity between the two net types, and there were several species commonly detected in the eDNA samples that were not captured in nets. Finally, we detected spatial heterogeneity in fish community composition across relatively small scales in Boardman Lake with eDNA metabarcoding, but not with traditional sampling. Our results demonstrated that eDNA metabarcoding was substantially more efficient than traditional gear for estimating community composition, highlighting the utility of eDNA metabarcoding for assessing species diversity and informing management and conservation.

## Introduction

Assessing biodiversity is critical to understanding ecosystem function and informing conservation (Gotelli & Colwell, 2011; Iknayan et al., 2014), but estimating species richness can be challenging, especially in aquatic environments (Gu & Swihart, 2004). An accurate understanding of fish community composition and species diversity is nevertheless essential to making informed fisheries management and conservation decisions (Helfman, 2007). For decades, fisheries managers have implemented different standardized tools to sample fish communities (Bonar et al., 2009; Zale et al., 2013), often utilizing multiple gear types to account for known biases with individual methods (Ruetz et al., 2007; Schneider, 2000). For example, electrofishing, gill netting, trawling, and fyke netting all target different groups and sizes of fishes in different habitats and depths, and a combination of these techniques are often used for assessments (Bonar et al., 2009; Zale et al., 2013). However, even if multiple sampling methods are used, some gear selectivity will persist, and sampling may not accurately represent true community composition (Schneider, 2000; Zale et al., 2013). Additional challenges also exist with traditional sampling including misidentification of species in the field, high cost, significant infrastructure and labor requirements, potential destruction of habitats and organisms, and failure to detect rare or elusive species (Deiner et al., 2017; Gotelli & Colwell, 2011; Iknayan et al., 2014; Thomsen & Willerslev, 2015). Often, these rare species are of particular interest to managers (i.e. endangered or invasive), and therefore false absences could lead to erroneous interpretations and inappropriate (or lack of) management action (Gu & Swihart, 2004; Thompson, 2013).

In recent years, environmental DNA (eDNA) metabarcoding has emerged as a useful tool for characterizing aquatic communities (Deiner et al., 2017; Thomsen & Willerslev, 2015). eDNA describes genetic material obtained from an environmental sample such as soil, sediment, snow, air, or water. As organisms interact with their surrounding environment, they shed DNA through excreted cells, sloughed-off tissue, gametes, and waste (Taberlet, 2012). This DNA can persist in the environment and be sampled to detect organisms without needing to physically handle specimens (Deiner et al., 2017; Rees et al., 2014). Metabarcoding utilizes universal primers that target an entire group of taxa in an eDNA sample, from which species barcodes are PCR amplified and sequenced on a high-throughput platform (Deiner et al., 2017; Porter & Hajibabaei, 2018).

Several studies have demonstrated that results from eDNA metabarcoding are comparable to traditional survey methods and/or long-term survey data to quantify fish community composition (e.g. Balasingham et al., 2018; Cilleros et al., 2018; Nakagawa et al., 2018; Pont et al., 2018). For example, Hänfling et al. (2016) collected water samples along established gill net sampling sites in a lake in the United Kingdom and detected 14 of the 16 fish species historically recorded there. Previous studies have also shown that that eDNA metabarcoding can detect more taxa than traditional survey methods in some instances (e.g. Afzali et al., 2020; Civade et al., 2016; Olds et al., 2016; Yamamoto et al., 2017; Zou et al., 2020). Additionally, Sard et al. (2019) demonstrated that eDNA metabarcoding could characterize 95% of a fish community with less sampling effort than traditional gear, and that eDNA detected aquatic invasive species that were not observed in some nets.

The field of eDNA metabarcoding has progressed substantially over the last decade, as researchers have developed reliable workflows for DNA extraction (Djurhuus et al., 2017; Lear et al., 2018), amplicon sequencing (Menning et al., 2018; Miya et al., 2015), and sequence analysis (Bolyen et al., 2019; Callahan et al., 2016a; Porter & Hajibabaei, 2018; Schloss et al., 2009). However, eDNA metabarcoding is still an evolving field and many improvements to sampling and study design could be warranted (Dickie, 2018; Ruppert et al., 2019). Some specific areas of eDNA metabarcoding studies that could be refined include sampling effort, amplicons/genes and databases used for taxonomic assignment, and the use of controls. For example, researchers may sample over a relatively large geographic area but collect only one, or very few, water sample replicates at a site or even for an entire lake. Additionally, some studies only amplify one gene region, which may lead to uncertainties in taxonomic assignment, or omit entire groups of taxa altogether (Shaw et al., 2016; Stat et al., 2017). Other studies may lack proper use of negative controls in the field and/or in the laboratory. Finally, some studies compare eDNA to only one traditional gear type, which could lead to misleading results due to the problem of selectivity mentioned above.

Our objectives were to intensively sample a small temperate lake over several weeks using two different traditional fish sampling methods and pair this sampling with eDNA metabarcoding of lake water samples using two mitochondrial DNA genes. We then compared estimates of fish community composition between traditional and eDNA sampling methods. Next, we assessed how the different net types sample fish communities differently (e.g. species selectivity among gears) and compared the taxa detected in lake water samples using the two metabarcoding genes (e.g. is a species detected with one gene but not the other?). Finally, due to the inherent patchiness of fish distributions, we determined if we could detect any temporal or spatial heterogeneity in fish community composition in the lake with either traditional gear or eDNA metabarcoding.

## Methods

### Study location and field collection

The Boardman River watershed encompasses 740 km^2^ in northwest lower Michigan (USA). In its lower reach, the river flows through Boardman Lake, a 137-hectare natural drowned river mouth lake, before meandering through Traverse City and then emptying into Grand Traverse Bay, an inlet of northeastern Lake Michigan (Figure 1). Boardman Lake and the upper Boardman River watershed are isolated from Lake Michigan by the Union Street Dam, an impassable earthen dam constructed downstream of Boardman Lake in 1867 roughly 1.5 km upstream of the river mouth (Kalish et al., 2018). Boardman Lake was selected for this study because the Great Lakes Fishery Commission (GLFC) was authorized to begin replacing the Union Street Dam with a fish passage structure to facilitate upstream movement of native fish species from Lake Michigan while blocking aquatic invasive species such as sea lamprey *Petromyzon marinus* (GLFC, 2018). Boardman Lake is not a well-studied system, however, and the GLFC is implementing several pre- and post-construction assessments in the lower Boardman River, including this metabarcoding research, to assess how fish assemblages and distributions may change in the future as a result of fish passage (http://www.glfc.org/fishpass.php).

**Figure 1:**
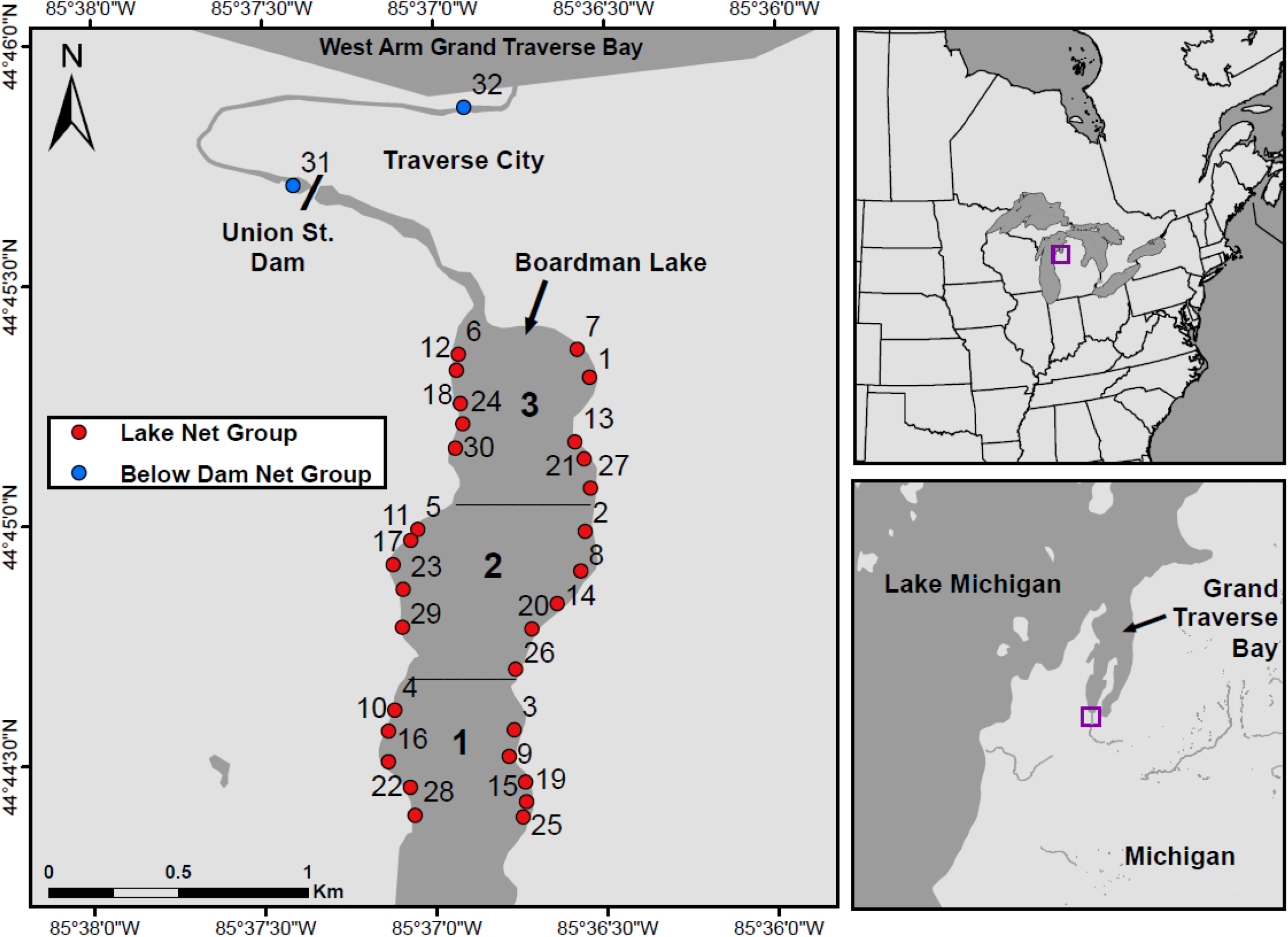
Sampling area with locations of net groups. We divided Boardman Lake into 3 sampling zones, each of which contained 10 net groups. Each net group consisted of one fyke net and one gill net and their corresponding lake water samples (3 per net). Water samples were also collected at 2 locations below the Union Street Dam on the lower Boardman River, but no corresponding nets were set because that river section is narrow and flowing with boat/kayak traffic

We intensively sampled Boardman Lake in Traverse City, MI from May to June in 2019. The lake was split into 3 sampling zones, with Zone 1 being the upstream and southern end of the lake, Zone 2 in the middle, and Zone 3 being the downstream and northern end (Figure 1). We attempted to make these zones roughly equal in surface area, but the southernmost end of the lake was extremely shallow and inaccessible to boats due to recent sediment deposition from dam removals upstream, so we omitted that portion of the lake from sampling and shifted zones northward. In each zone, we sampled 10 locations with five locations on the east side of the lake and five locations on the west side (Figure 1). We attempted to equally space sampling locations within each zone, but due to private property and the presence of docks, we occasionally had to shift sample sites. At each sample site, we set a mini fyke net against the shore and an experimental gill net 5 - 10 meters out from the end of the fyke net, perpendicular to the shoreline. Each pairing of the two net types was later combined into a single “net group” for analysis, with a total of 60 individual nets set and 30 net groups in the lake. Fyke nets had rectangular frames that were 92 cm wide x 60 cm high, with two frames per net. Circular hoops were 60 cm in diameter with three per net, and the mesh size for the entire net was 9.5 mm. Net leads were 7.6 m in length and 92 cm high. Fyke nets were set so the rectangular frames were just beneath the surface of the water (0.7 – 1 m depth). Experimental gill nets were 38 m x 1.8 m with five 7.6 m graded meshes of 3.8 cm stretch, 5.1 cm stretch, 6.4 cm stretch, 7.6 cm stretch and 10 cm stretch. Gill nets were set with the narrow mesh towards the shore (shallow) and the larger mesh away from the shore (deep). Depth of gill nets varied among sites due to the variable nature of the lake’s bathymetry. Depth of the shallow end of the net ranged from 2 – 4 m, and the deep end ranged from 3 - 12 m, depending on location. Fyke nets tend to sample shallow littoral species, and our mini fyke nets target smaller fish due to their relatively small mesh size. Gill nets tend to sample larger pelagic fish, and in general both nets rarely catch benthic or sedentary species (Zale et al., 2013).

At each net set location, we measured water temperature, recorded GPS coordinates, and assigned a unique waypoint number. From the boat and before setting the net, we collected three 1 L surface water samples into 1 L Nalgene wide-mouth HDPE bottles (ThermoFisher Scientific, Waltham, MA). Bottles were previously decontaminated by soaking in a 10% bleach solution for at least 10 minutes. Water samples were collected prior to setting nets to avoid any contamination from the net itself. Each collection event (per fyke or gill net) consisted of three replicates of lake water and one negative control (“field negative”), which was a 1 L Nalgene bottle filled with 1 L distilled water, brought into the field, and handled like all other samples. Gloves were always worn when collecting samples and were changed at each net set location. All bottles of water were stored on ice in a cooler that was decontaminated daily with 10% bleach. After collecting the three lake water samples (including one negative control), we set the corresponding gill or fyke net. We set one net group per zone each sampling day (three net groups, six total net sets) and left nets to fish overnight. The next day, we removed fish from nets, identified species, and recorded total length in mm for each fish. We performed 10 of these sampling days over a one-month period. With 60 individual net sets and three lake water samples and one negative control for each, we collected a total of 180 water samples and 60 negative controls from Boardman Lake.

In addition to the water samples corresponding to net sets in Boardman Lake, we also collected water samples below the Union Street Dam at two locations on the lower Boardman River. Since this river section is directly connected to Lake Michigan, we wanted to see if we could detect any species there that we did not observe in Boardman Lake. Water was collected from each location on three different dates from May to June 2019. As described above, each sampling event consisted of three replicates of river water plus one negative field control. We collected and filtered these water samples in the same manner as the Boardman Lake samples. With three sampling events at two sites consisting of three water samples plus one negative control per site, we had a total of 24 water samples from the lower Boardman River. For later analyses, each downstream sampling location was designated as a unique “net group”.

At the end of each sample collection day, we filtered 1 L of water from each bottle through 47 mm 0.45 μm nitrocellulose Nalgene analytical test filter funnels (Thermo Fisher) using a Gemini 2060 Dry Vacuum Pump (Welch, Prospect, IL). All filtering supplies and instruments were sterilized by soaking in a 10% bleach solution for at least 10 minutes. Between each sample, we changed gloves and sterilized the work benches. Negative controls were filtered in the same manner as lake water samples. Filters were preserved in 95% ethanol in individual 50 ml tubes and stored at room temperature. Including negative field controls and water samples from Boardman Lake and Boardman River below Union Street Dam, we processed a total of 264 water bottles (see Table S1 for water sample metadata).

### Sequencing library preparation

Filters were cut in half; one half was used for extraction, and the other half was left in ethanol in its respective vial in the event we needed to re-extract filters due to potential problems downstream. Ethanol was allowed to evaporate from the half-filters for 24 hours before extraction. We followed the extraction methods outlined in Sard et al. (2019) and Laramie et al. (2015), using a combination of the Qiagen DNeasy Blood & Tissue Kit and the Purification of Total DNA from Animal Tissues Spin-Column Protocol (ThermoFisher). We performed two elutions of 100 μl each using Buffer AE warmed to 70℃. Eluted DNA was treated with Zymo OneStep PCR Inhibitor Removal columns (Zymo Research). Extractions were performed in a UV-sterilized laminar flow hood and all extraction instruments and bench spaces were sterilized with a 10% bleach solution or UV light. All pipetting was performed using sterile barrier filter tips.

Extracted DNA was then transferred from the elution tubes into 96-well plates, so that each plate contained 94 eDNA samples, a “lab negative” control (distilled sterile H_2_O), and a positive control. Our positive control was DNA extracted from a caudal fin clip of *Pygocentrus natterei*, a tropical fish not found in our study system, which we included to visualize any potential cross contamination between wells. We then amplified extracted DNA at two mitochondrial genes, 12S and 16S, using primers also used by Sard et al. (2019) in their study on similar freshwater systems in Michigan. For 12S, the universal primers were originally developed by Riaz et al. (2011): Forward: 5’-ACTGGGATTAGATACCCC-3’, Reverse: 5’- TAGAACAGGCTCCTCTAG-3’. For 16S, we used universal primers originally developed by Deagle et al. (2009) and modified by Sard et al. (2019) to better match diversity in Michigan lakes: Forward: 5’-CGAGAAGACCCTNTGRAGCT-3’, Reverse: 5’CCKYGGTCGCCCCAAC- 3’. Each core primer was synthesized with an Illumina tail on the 5’ end complementary to the adapters used during barcoding described below. Forward primers were tailed with the Small RNA Sequencing Primer (5’-CGACAGGTTCAGAGTTCTACAGTCCGACGATC-3’), while reverse primers were tailed with the Multiplexing Read 2 Sequencing Primer (5’- GTGACTGGAGTTCAGACGTGTGCTCTTCCGATCT-3’).

PCR reactions were conducted in 10 μl volumes with 3 μl of DNA and 7 μl of a master mix. The master mix for the 12S gene contained 1X NEB 10X Standard Taq Reaction Buffer, and concentrations of 0.24 mM dNTPs, 2 mg/ml MgCl2, 1 mg/ml BSA, 1.25 U/μl NEB taq, and 0.8 μM forward and reverse primers. The master mix for the 16S gene contained 1X NEB 10X Standard Taq Reaction Buffer, and concentrations of 0.32 mM dNTPs, 2.5 mg/ml MgCl2, 1 mg/ml BSA, 1.25 U/μl NEB taq, and 0.8 μM forward and reverse primers. Thermal cycling was performed as follows: 95 °C hold for 2 min, followed by 35 cycles of 95 °C for 30 s, 57 °C for 30 s for 12S and 45 s for 16S, 72 °C 45 s and a final extension of 72 °C for 5 min.

We then produced the final metabarcoding libraries using latter steps in the genotyping-in-thousands by sequencing (GT-seq) protocol (Campbell et al., 2015) following Bootsma et al. (2020). Briefly, each individual and plate was barcoded in a second indexing PCR, and barcoded products were normalized using SequalPrep DNA Normalization plates (Invitrogen, Carlsbad, CA). Normalized products were pooled within plates, and the pooled libraries were purified and size-selected using AMPure XP (Beckman Coulter, Brea, CA). Products were visualized on a 2% E-Gel EX agarose gel run on a Power Snap Electrophoresis Device (ThermoFisher) to verify size ranges and quantified using a Qubit 2.0 Fluorometer and dsDNA HS Assay Kit (ThermoFisher). Including positive and negative lab controls, we then sequenced 540 dual-indexed 12S and 16S amplicons on two Illumina MiSeq 2×150 flow cells at the University of Wisconsin Biotechnology Center DNA Sequencing Facility in Madison, WI.

### Data filtering and quality control

We concatenated raw reads from both sequencing runs for each sample and used the program cutadapt (Martin, 2011) to remove adapter contamination from the 5’ and 3’ ends of the 12S and 16S sequences. We then processed cleaned data with the DADA2 package (Callahan et al., 2016a) in R (R Core Development Team). Due to variable sequence lengths, we retained reads between 130 and 145 bp for 12S, and 80 and 145 bp for 16S. For both genes we used the pool = TRUE parameter due to its increased sensitivity to rare sequences (Callahan et al., 2016b). After filtering, denoising, pooling, merging, and removing chimeras in DADA2, we exported sequence tables containing the final amplicon sequence variants (ASVs) and the number of reads per sample.

We used BLASTn (Altschul et al., 1990) to locally BLAST our 12S and 16S sequence tables to reference databases created by Sard et al. (2019), which included species common in Michigan lakes. Using customized R scripts, we parsed aligned ASVs and retained matches with greater than 98% sequence identity and alignment lengths greater than 140 bp for 12S and 80 bp for 16S. These lengths were chosen after exploration of raw data revealed robust 16S alignments down to 80 bp for taxa known to inhabit the Boardman River watershed but few robust alignments below 140 bp for 12S. After filtering, several ASVs still remained with no matches to our reference database, and we used BLASTn to align these sequences to the National Center for Biotechnology Information (NCBI) nucleotide database. We then used the same parameters as above to retain alignments. Sequences that were added through alignment to the NCBI database were combined with the original database to create a fully comprehensive reference. We then reBLASTed our sequences against the updated reference databases using BLASTn locally and ran outputs through our R script using the same cutoffs as above. Any ASVs with no matches after this step were removed, and all species other than fish were removed. ASVs with a single match that met our parameters or that had multiple matches for a single species (unambiguous) were assigned that species. ASVs with a match to two or more species (ambiguous) were either assigned back to common genus, or in some cases, to common family when multiple genera were present. However, some taxonomic groups presented exceptions to these rules due to the lack of within-genera variation within a particular gene and the potential for hybridization between closely related species. We thus collapsed any species in *Oncorhynchus*, *Lepomis*, *Rhinichthys*, and *Cottus* to genus. Finally, because multiple ASVs assigned to the same species, we collapsed all ASVs to unique assignments and summed reads for each sample.

To minimize false-positive detections, we accounted for contamination by subtracting reads in our controls from reads in water samples as follows: For each set of three water sample replicates, we subtracted the maximum number of reads for any single species found in the field negative, the lab negative, or the lab positive for each species other than *P. natterei*, (i.e., whichever control produced the highest per species contaminating read count) from all reads in the corresponding water samples. Species with more than 10 reads after subtracting contaminated reads were considered true hits. After accounting for contamination and removing hits with reads less than 10, we constructed community matrices for 12S and 16S using both read counts and presence/absence. We then grouped Boardman Lake samples by waypoint (three lake water samples) and net group (six lake water samples). For the samples taken below the Union St Dam, each waypoint was also considered a unique net group and consisted of nine samples (three sampling events consisting of three water samples each). A species was considered present at a given waypoint or net group if it was detected in at least one water sample replicate.

We also constructed presence/absence and abundance datasets for the traditional sampling gear to facilitate comparisons to the eDNA dataset. Abundance was the count of individuals from each species in each net, and species were considered present if they were found in the net. We then combined and summed abundances from each set of gill and fyke nets to acquire a total abundance of each species for each net group. To ensure that our comparison of species between methods was accurate, we changed the name of some taxa found in nets to match our eDNA assignments. For example, any *Lepomis macrochirus* detected in the eDNA was assigned to “*Lepomis* sp.” and any species in the genus *Oncorhynchus* were assigned to “*Oncorhynchus* sp.”, therefore, any *L. macrochirus* or *O. mykiss* species caught in nets were assigned back to genus for comparison with the eDNA data.

### Statistical analyses

The first goal of our data analysis was to quantify and visualize basic trends in our data. We constructed boxplots of read counts and catches for each gene and gear and used Student’s 2-sample t-tests for comparisons. We assessed the correlation between number of reads and number of unique taxa detected in each sample for both 12S and 16S using Spearman’s correlation coefficient. We used Pearson’s correlation to assess the relationship between the number of instances a taxon was detecting with 12S and 16S. Additionally, we constructed one heat map to visualize the species in each net group detected with 12S, 16S, or both genes, and a second heatmap to visualize the species detected with gill nets, fyke nets, or both gears. A species was considered “detected” if it appeared in any of the replicates for a given net group. We also constructed separate heatmaps for each gene quantifying the number of eDNA water samples (out of six) where a species was detected by each gene in each net group. Finally, we assessed trends in our negative control samples by constructing a boxplot of total reads in each negative control sample for each gene, a Pearson correlation comparing species-specific detections for 12S compared to 16S in negative controls, and a Spearman correlation comparing the number of eDNA detections for a given species to the number reads of that species found in negative controls summed across both genes. We set α=0.05 for all tests.

To assess how each sampling method (12S eDNA, 16S eDNA, fyke nets, and gill nets) surveyed the fish community in Boardman Lake, we developed species accumulation curves using the “exact” method for the specaccum function in the R package vegan (Oksanen et al., 2019). Curves were constructed for all four methods separately and to compare eDNA (12S and 16S combined) with traditional sampling (fyke and gill nets combined). For combined eDNA and combined traditional gear curves, a species was considered present if it was found in either gear or gene.

We correlated eDNA read counts to catch numbers and biomass of fish sampled in traditional gear to explore the relationship between eDNA read counts and fish abundance. This analysis was isolated to five species that were found most frequently in the traditional gear: *Ambloplites rupestris*, *Catostomus commersonii*, *Esox lucius*, *Sander vitreus*, and *Perca flavescens*. Pearson correlations (read counts in eDNA vs fish counts/biomass in nets) were conducted separately for each gene/gear combination for each of the five most common species. Biomass data for each net was estimated by converting total lengths to weights following species-specific conversion equations in Schneider et al. (2000).

A major objective of this project was to determine whether estimates of fish community composition were influenced by gear, gene, sampling method, spatial location, or other environmental variables. We used presence/absence data and two multivariate approaches, non-metric multidimensional scaling (NMDS) and redundancy analysis (RDA), to address this objective. NMDS was conducted to compare estimates of community composition between 12S and 16S, gill and fyke nets, eDNA (combined 12S/16S) and traditional gear (combined fyke and gill nets). NMDS was also used to compare community composition between samples taken in different zones, different sides of the lake (E vs W), and above and below the Union Street Dam. NMDS analysis by zone and side was conducted for both eDNA data (combined 12S/16S) and traditional data (combined gill and fyke net data), while NMDS comparing composition above and below the dam was only conducted for eDNA as no data for traditional gear were available below the dam. NMDS was conducted with the Bray-Curtis dissimilarity matrix and three ordination axes were generated for each analysis. We explored generating two and four ordination axes and found that observed patterns were similar. We therefore chose to standardize our analyses to retain three ordination axes as this appeared to generally capture a large amount of the variation in the data while minimizing NMDS stress. NMDS analyses were visualized in R, and 95% confidence intervals around each group of datapoints were drawn with the dataEllipse function in the car package (Fox & Weisberg, 2019). We conducted analysis of similarities (ANOSIM; Clarke, 1993) for each NMDS dataset using the R package vegan to assess significant differences in community composition among analysis groups.

We conducted RDA analysis in the R package vegan for eDNA and traditional data to determine the influence of sampling location (zone and side of lake), water temperature, and sampling date on community composition. Our choice to conduct RDA instead of canonical correspondence analysis, a similar multivariate method, was informed by a detrended correspondence analysis, which indicated that variables were linear thus more appropriate for RDA (Legendre & Legendre, 1998). To ensure the robustness of our RDA analyses, we explored whether row normalizing our species presence/absence data or removing rare species (from one to three detections) influenced overall patterns. Additionally, we conducted RDA with all variables and with a single variable from each group of highly correlated variables as suggested by Hair (1995). The significance of each variable (i.e. their influence on variation in community composition) was assessed with ANOVAs. For both multivariate analyses, we used α=0.05.

## Results

### eDNA metabarcoding – 12S vs 16S

For 12S, we began with 12,702,289 raw reads across all samples. After filtering, merging, and chimera removal in the DADA2 pipeline, 7,249,412 reads remained (57.1%). Of the 538 ASVs output from DADA2, 217 remained after applying our filtering parameters, aligning with reference sequences, and removing non-fish organisms. For 16S, we began with 13,202,450 raw reads across all samples. After the DADA2 pipeline, 3,458,683 reads remained (26.2%). Of the 8,855 ASVs output from DADA2, 784 remained after applying our filtering parameters and removing non-fish organisms. There were significantly more reads among samples in 12S (mean=25,407; SD=17,741) than 16S (mean=7,401; SD=10,739) (*p*<0.001, t = 13.522, df = 197) (Figure S1), but the average number of taxa detected by each gene did not significantly differ (*p*=0.76).

After combining filtered ASVs based on common assignments, both genes combined detected 40 unique taxa (Table 1, Figure 2). Both 12S and 16S individually detected 32 unique taxa, although taxonomic assignments were slightly different between the two genes (Table 1, Figures 2, S2, & S3). For example, 12S was able to unambiguously classify species in the genus *Etheostoma*, while 16S could not. 16S detected the *Lampetra* genus and *Petromyzon marinus*, while 12S had zero assignments to any lamprey species. Additionally, due to a lack of sequence variation in salmonids for the 12S gene, several ASVs assigned to more than one species or genus within the family, so we assigned these ASVs to “Salmonidae”. For 16S, however, we did not need to assign back to Salmonidae because assignments were mostly unambiguous to genus. Both genes were often ambiguous for the Cyprinidae family, with a single ASV often assigning to multiple genera, so for both 12S and 16S we assigned these to “Cyprinidae”. 16S appeared to be slightly better at classifying species in this family however, for example detecting *Luxilus cornutus* and *Pimephales notatus*. Overall, 12S detected four unique taxa, while 16S detected seven unique taxa (Table 1). The average number of taxa detected per net group were 14 (SD=3.95) and 13 (SD=3.5), for 12S and 16S, respectively (Table S2). There was a weak positive correlation between the number of species detected in a sample and the number of reads for both 12S and 16S (Figure S4), and the number of detections of common species between genes was positively correlated (r = 0.91; Figure S5). eDNA samples collected below the dam detected similar species to those in Boardman Lake, but some taxa were more common in, or exclusive to, the downstream samples, for example *Alosa sp*. (likely alewife), *Catostomus catostomus* (longnose sucker), *Coregonus artedi* (cisco), *Notemigonus crysoleucas* (golden shiner), *Percina caprodes* (common logperch), and *P. marinus* (sea lamprey) (Figures 2, S2, & S3). It is important to note that although some ASVs from water samples matched to *C. artedi*, it is possible that we could be detecting *C. clupeaformis* (lake whitefish), as this species is also present in Lake Michigan and very little sequence variation exists between these two closely related species.

**Table 1:** Occurrences of all detected taxa using eDNA (12S and 16S genes) and traditional sampling gear (gill nets and fyke nets). Values in “12S” and “16S” columns are the number of net groups out of 32 in which a particular taxon was detected, and values in the “fyke” and “gill” columns are the number of net groups out of 30 in which a species was detected. Darker backgrounds indicate higher values. Asterisks indicate broader taxa (genus or family) in which a narrower assignment occurred for that method, which we used to avoid double-counting redundant assignments (for example, *Etheostoma* sp. for the 12S gene which was able to detect both *Etheostoma exile* and *Etheostoma nigrum*), therefore only taxa without associated asterisks were counted as “unique” for each method.

| Scientific Name | Common Name | 12S | 16S | Fyke | Gill |
| --- | --- | --- | --- | --- | --- |
| Alosa sp. | River herrings | 2 | 1 |  |  |
| Ambloplites rupestris | Rock bass | 28 | 29 | 28 | 15 |
| Ameiurus sp. | Bullheads | 3 | 1 |  |  |
| Catostomidae | Suckers | 1 | * | * | * |
| Catostomus catostomus | Longnose sucker | 2 | 3 |  |  |
| Catostomus commersonii | White sucker | 32 | 32 | 5 | 18 |
| Coregonus artedi | Cisco | 3 |  |  |  |
| Cottus sp. | Sculpins | 28 | 26 |  |  |
| Culaea inconstans | Brook stickleback | 10 | 11 |  |  |
| Cyprinidae | Minnows & carps | 18 | 14 |  |  |
| Cyprinus carpio | Common carp | 10 | 6 |  |  |
| Esox lucius | Northern pike | 31 | 28 | 3 | 22 |
| Esox masquinongy | Muskellunge | 1 |  |  |  |
| Esox sp. | Pikes | 3 | * | * | * |
| Etheostoma exile | Iowa darter | 25 |  |  |  |
| Etheostoma nigrum | Johnny darter | 13 |  |  |  |
| Etheostoma sp. | Darters | * | 30 | 5 |  |
| Lampetra sp. | Lampreys |  | 19 |  |  |
| Lepomis sp. | Sunfishes | 8 | 7 | 2 |  |
| Luxilus cornutus | Common shiner |  | 2 |  |  |
| Micropterus dolomieu | Smallmouth bass | 14 | 20 | 2 | 1 |
| Micropterus salmoides | Largemouth bass | 5 | 4 | 1 |  |
| Neogobius melanostomus | Round goby | 32 | 31 | 7 |  |
| Notemigonus crysoleucas | Golden shiner | 1 | 2 |  |  |
| Oncorhynchus sp. | Pacific salmon & trout | 24 | 24 | 1 | 1 |
| Perca flavescens | Yellow perch | 32 | 31 | 7 | 27 |
| Percina caprodes | Common logperch |  | 1 |  |  |
| Percopsis omiscomaycus | Trout perch | 24 | 22 |  |  |
| Petromyzon marinus | Sea lamprey |  | 2 |  |  |
| Pimephales notatus | Bluntnose minnow |  | 1 |  |  |
| Pomoxis nigromaculatus | Black crappie | 1 | 2 |  |  |
| Rhinichthys sp. | Riffle daces | 13 | 18 |  |  |
| Salmo trutta | Brown trout | 22 | 20 | 1 | 3 |
| Salmonidae | Salmonids | 15 | * | * | * |
| Salvelinus fontinalis | Brook trout |  | 4 |  |  |
| Salvelinus namaycush | Lake trout |  | 1 |  |  |
| Salvelinus sp. | chars and trouts | 4 | * |  |  |
| Sander vitreus | Walleye | 28 | 24 |  | 20 |
| Semotilus atromaculatus | Creek chub | 8 | 3 |  |  |
| Umbra limi | Central mudminnow | 4 | 3 |  |  |
| n unique |  | 4 | 7 | 0 | 0 |

**Figure 2:**
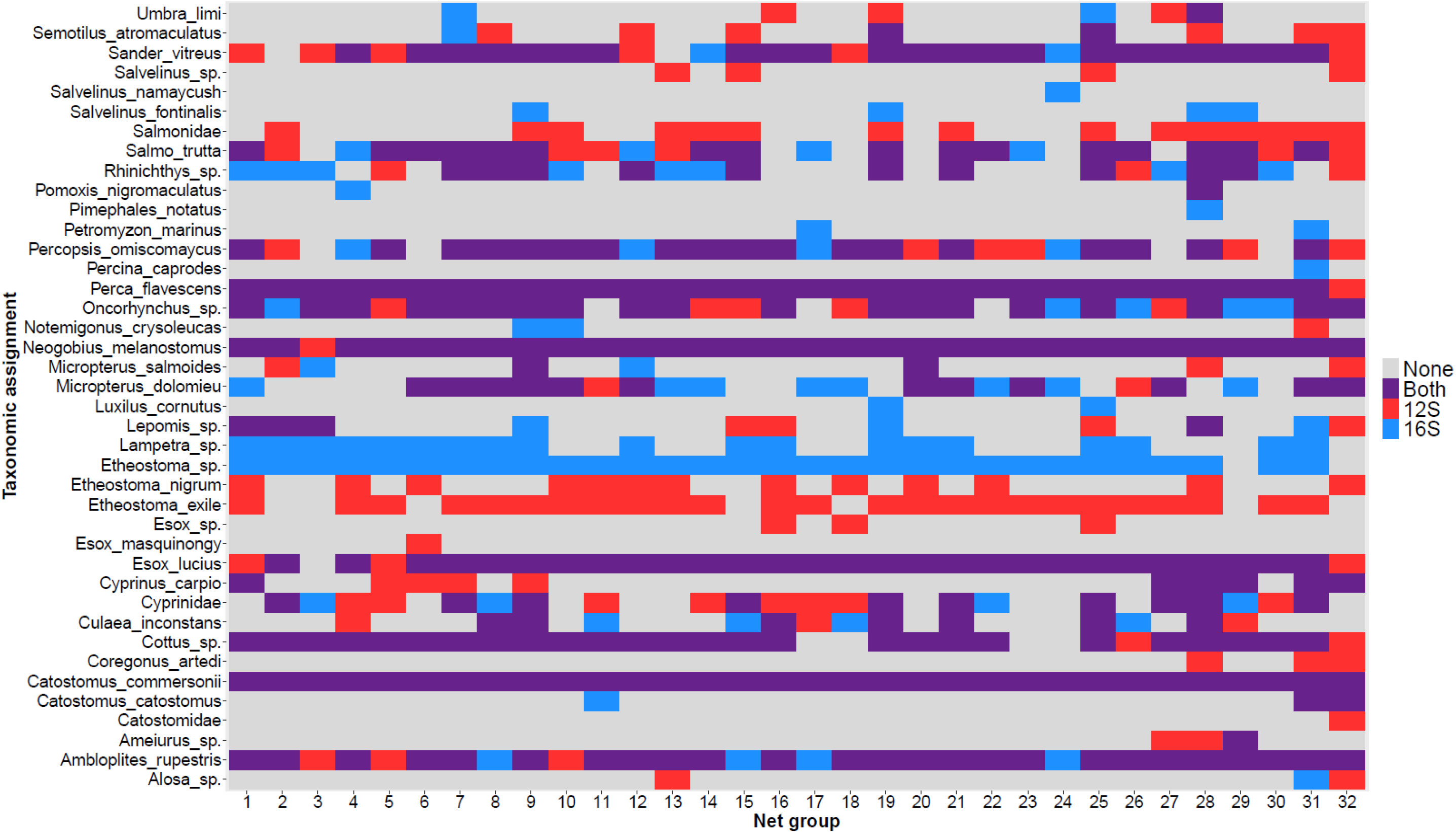
Heatmap indicating presence or absence of all potential taxa by net group for each gene used for eDNA metabarcoding. Blue indicates taxa present in a net group that was detected only with 12S, red indicates a detection only with 16S, purple indicates that taxa were detected with both genes, and grey signifies no detection. Each net group consisted of one fyke net and one gill net and their corresponding lake water samples (3 per net). Note that net groups 31 and 32 are the below-dam river sample sites.

### Contamination in controls

Contamination in field negatives overall was low (i.e. generally less than 10 reads per species at maximum; Figure S6), although three field negatives contained high numbers (>1000) of reads. Two of these were contaminated with *S. vitreus* and one was contaminated with *C. commersonii* (Supplementary Files 1 & 2), two species that were common in the system based on net catches and reads in lake water samples. We removed these outliers for the analyses on field negative controls reported here and speculate on why they may have occurred in the discussion. There was a positive correlation between the number of detections for a species and the number of read counts in negative controls (Figure 3), suggesting that species more common in the system were also more likely to amplify in our field negative controls. Contamination by species was also correlated between 12S and 16S (r = 0.75; Figure S7), suggesting that contamination per species likely occurred before PCR or sequencing.

**Figure 3:**
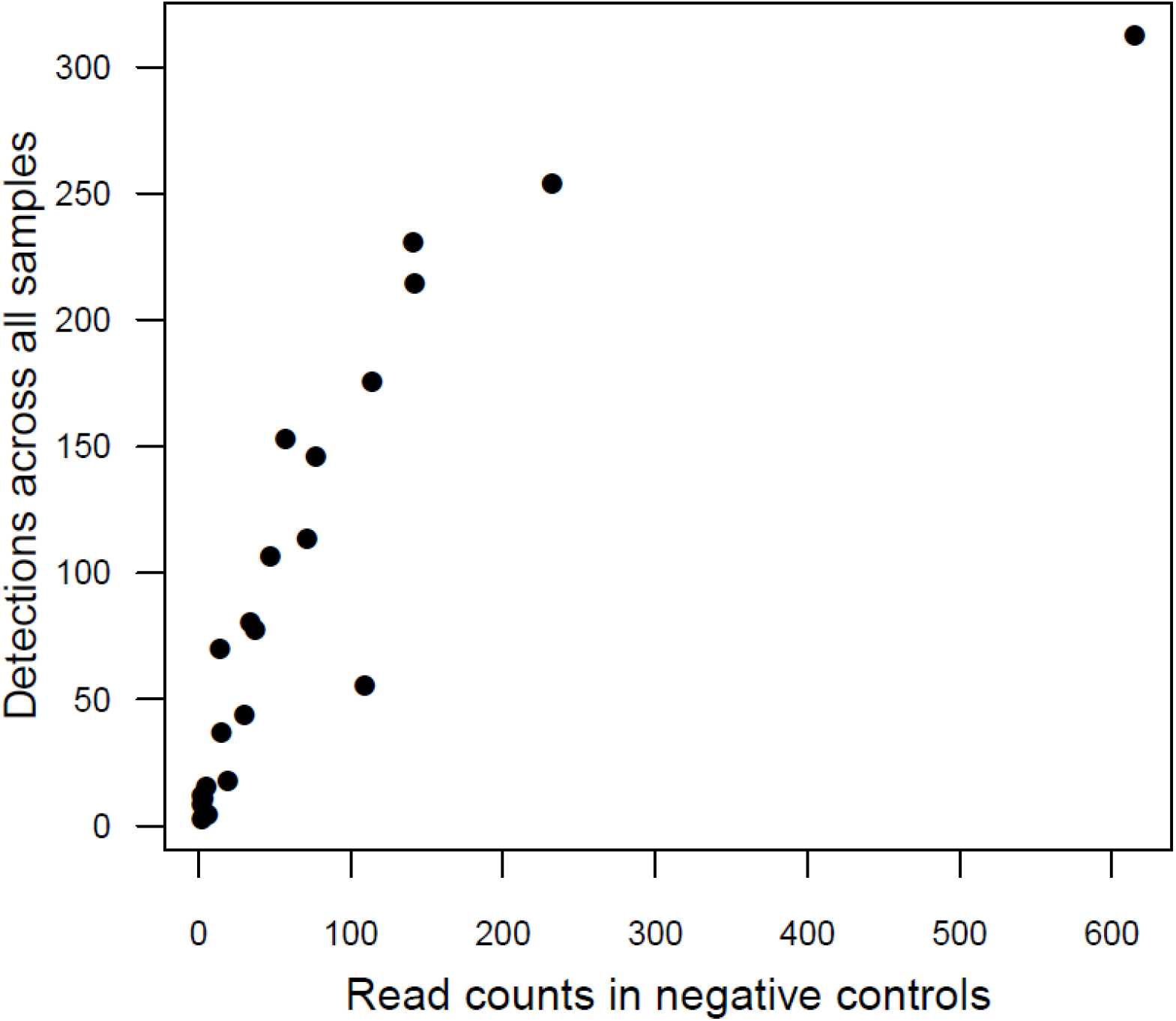
Relationship between the number of detections across all eDNA samples and number of reads in all field negative controls for all detected taxa assessed with Spearman’s correlation coefficient (r_s_=0.935). Each point represents a unique taxon. Three outlier negative controls with high read counts were removed prior to analysis (see results). Data includes both Boardman Lake and below-dam samples.

Our use of the lab positive control was successful, with both 12S and 16S amplifying tens of thousands of reads for *P. natterei* (Supplementary Files 1 & 2). Reads for this species in most other samples was zero, and if any reads existed, they were never greater than two, and were later removed during our filtering step. Contamination of other species in lab negative and lab positive controls was also low overall, with most species never amplifying, although there was a small amount of contamination from some species. For 12S, the highest number of reads in the lab controls was six (from *C. commersonii* in the lab positive control), and for 16S the highest number of reads in the lab controls was 13 (also from *C. commersonii* in the lab positive control) (Supplementary Files 1 & 2).

### Traditional gear - Gill vs fyke nets

Using traditional gear, we caught a total of 12 species. Gill nets caught a total of 350 fish representing 8 unique species, and fyke nets caught a total of 195 fish representing 11 unique species (Table S2). Gill nets caught significantly more fish than fyke nets on average (mean=11.7, SD=7.6 for gill and mean=6.5, SD=4.5 for fyke, *p*=0.004; Figure S8). The average number of unique species caught in a single net were 4 (SD=1.431) and 2 (SD=1.048) for gill nets and fyke nets, respectively (Table S2). Species most common in gill nets were *P. flavescens* (38%), *E. lucius* (18%), *S. vitreus* (15.4%), and *A. rupestris* (14.6%). Species most common in fyke nets were *A. rupestris* (68.2%), followed by *P. flavescens* (9.7%) and *Neogobius melanostomus* (6.7%) (Table S3). Fyke nets caught 4 taxa that were not detected in gill nets (*Etheostoma sp*., *Lepomis sp.*, *Micropterus salmoides*, and *N. melanostomus*). Gill nets caught only one species that was not present in fyke nets (*S. vitreus*; Figure S9). In general, most species were either rare in nets or tended to be susceptible to only a single net type (Figure S9).

### Traditional gear vs eDNA

Overall, eDNA metabarcoding detected significantly more taxa than traditional gear (*p*<0.001). Both 12S and 16S detected all 12 species caught using traditional gears (Table 1). These species were detected more frequently in the eDNA compared to traditional gears, or there were detections with both methods within a given net group (Figure 4). Only two net groups had a detection for a species observed in a net but not detected in the corresponding eDNA samples. Between 12S and 16S, 24 more unique taxa were detected in the eDNA than in the traditional gears (Table 1).

**Figure 4:**
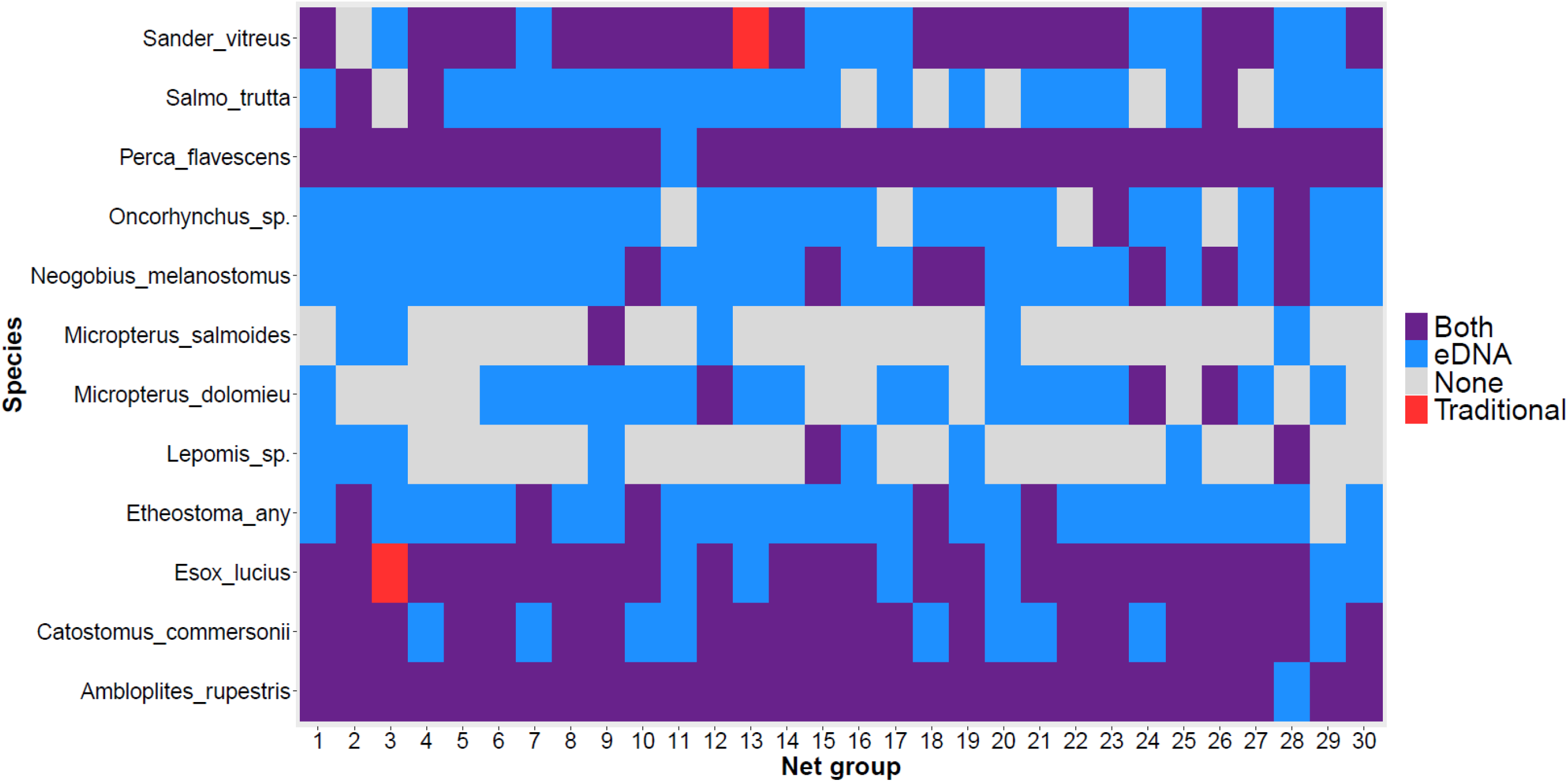
Heatmap indicating presence or absence of all species detected using traditional sampling gears (both fyke and gill nets), and comparison with corresponding detections using eDNA (12S and 16S combined) for each net group in Boardman Lake. Blue indicates that a particular species was detected in a given net group with eDNA only, red indicates it was detected only using traditional sampling, purple indicates it was detected using both methods, and grey signifies no detection with either method.

Among the 12 species detected using traditional gear, there appeared to be three main groups in terms of species abundance and catchability (Figure 5): (1) species present in low numbers in both nets and in eDNA (low abundance in the lake), (2) species present in low numbers in the nets but common in the eDNA (likely high abundance in the lake but with low catchability for our traditional sampling gears), and (3) species common in nets and in the eDNA (species very abundant in the system and also susceptible to our sampling gears).

**Figure 5:**
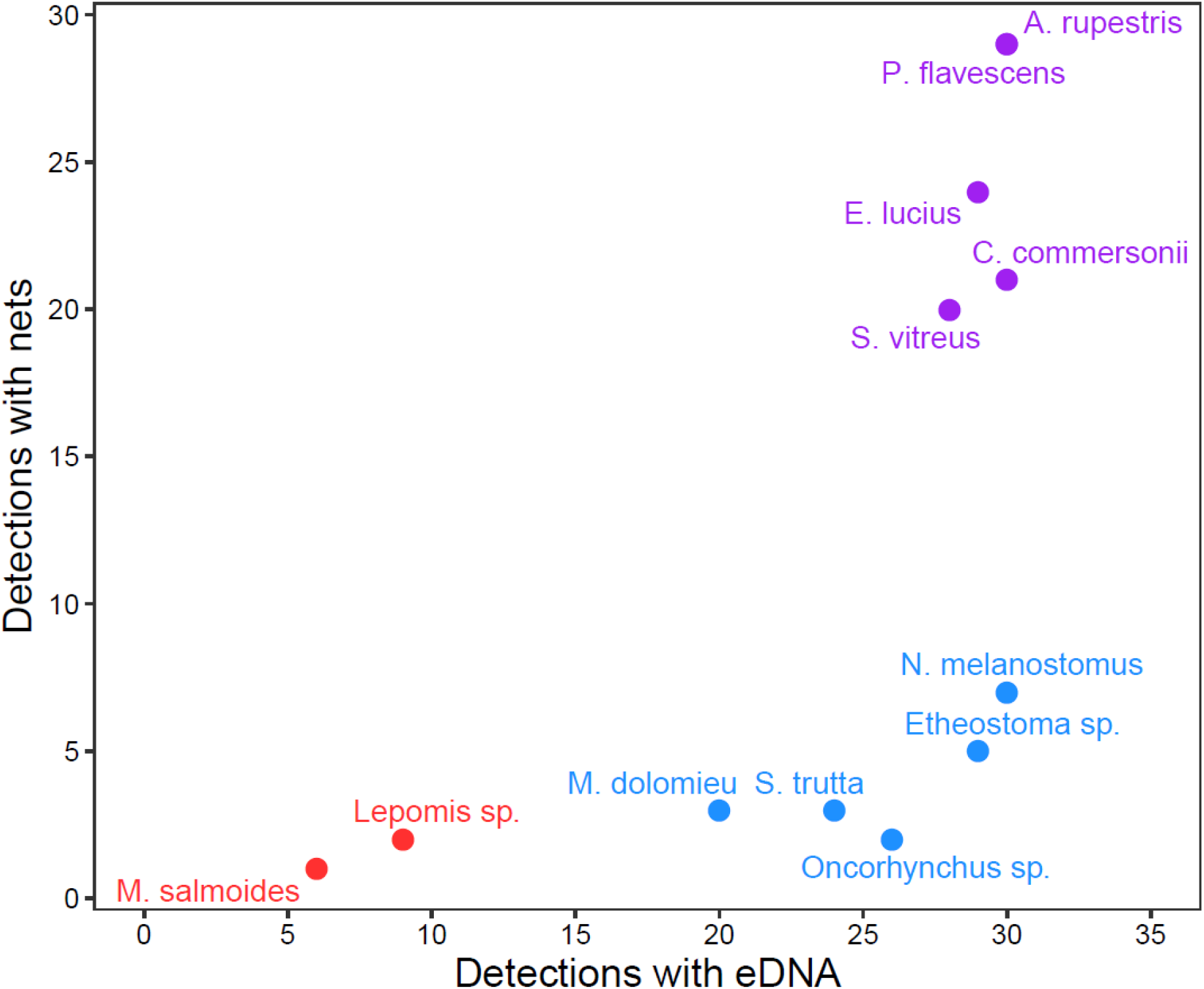
Relationship between the number of detections in eDNA (12S and 16S combined) and detections using traditional gear (fyke and gill nets combined) for species that were detected using both methods in Boardman Lake, with colors representing 3 distinct detection groups. Axes represent the number of net groups in which a taxon was detected. Red represents species that had low presence in both eDNA and traditional gears, suggesting low abundance in the system. Blue indicates species that were commonly detected using eDNA but not traditional gear, suggesting they are abundant in the lake but not susceptible to net sampling. Purple represents species with high detection rates using both methods, suggesting they are abundant in the lake and also susceptible to traditional sampling gears.

The species accumulation curves demonstrate the higher number of species detected using eDNA compared to gill or fyke nets, and that relatively fewer eDNA samples are needed to reflect species richness in the lake (Figure 6). Also, gill nets detected more species than fyke nets at first, but with an increasing number of samples, fyke nets detected more species than gill nets overall (11 vs eight species, respectively). 12S and 16S had similar and mostly overlapping accumulation curves.

**Figure 6:**
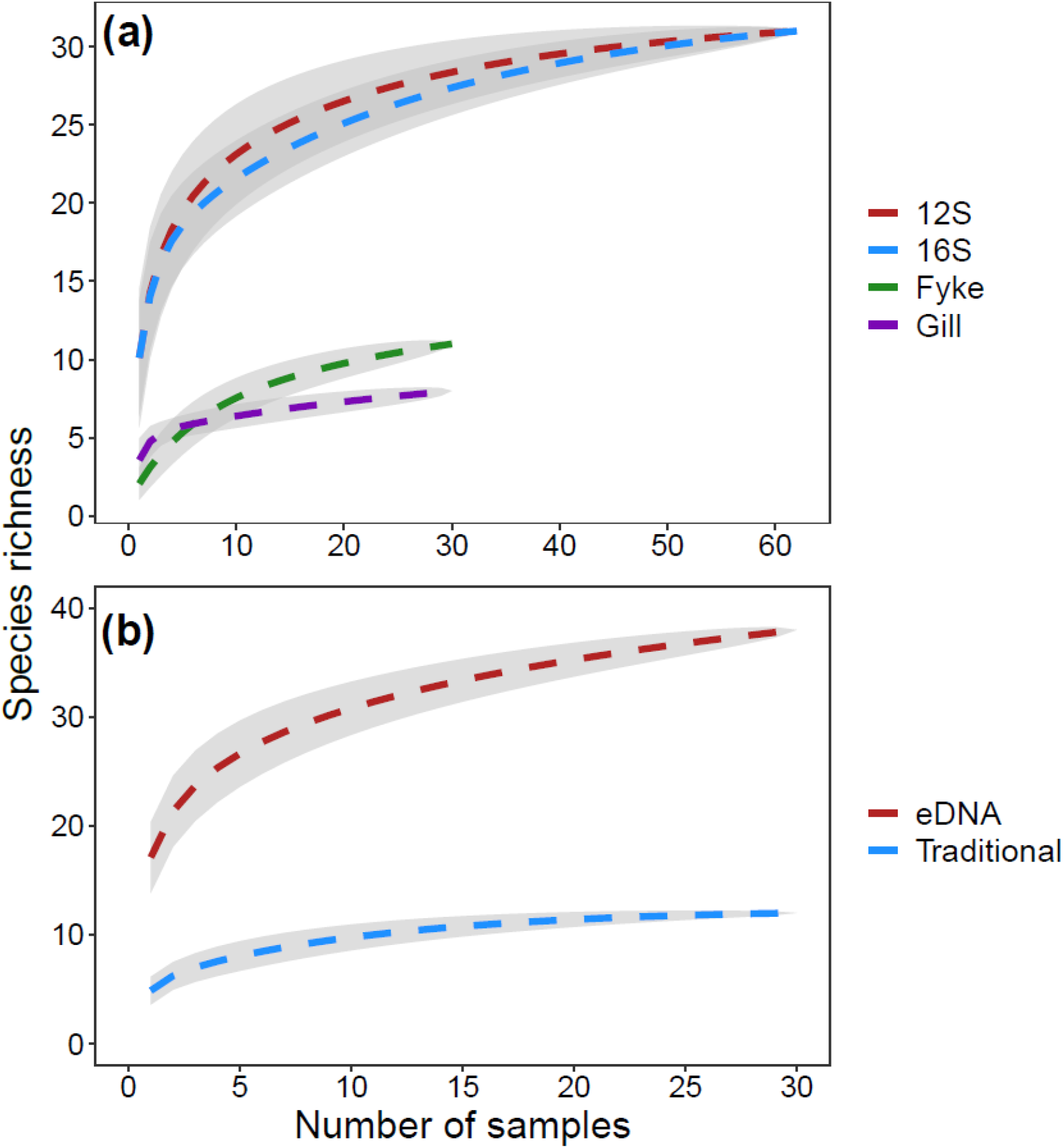
Species accumulation curves for Boardman Lake for a) each detection method based on species presence at individual waypoints (i.e. a net site where a fyke or gillnet was set and an eDNA sample was taken), and b) eDNA (12S and 16S combined) compared to traditional gear (gill and fyke nets combined) based on species presence in net groups. Grey shading represents the standard deviation of the expected number of species per sampling method.

Overall, the number of reads in both 12S and 16S of our five most common species did not consistently correlate with either biomass or number of fish caught in each net, suggesting that read count data are unreliable to estimate abundance based on net catches, at least in this system (Figures S10 & S11). Of the 60 correlations we assessed, only one had a *p* value < 0.05, which compared the number of yellow perch caught in fyke nets to the number of yellow perch reads in the corresponding 12S eDNA water samples (Figure S10n).

### Multivariate analysis results

Of the eight NMDS comparisons we performed to describe community compositions among variables and methods, there were five comparisons in which the ANOSIM analysis was significant: 12S vs 16S, Gill vs Fyke, eDNA vs traditional, eDNA by lake side, and eDNA above dam vs below dam (Table 2, Figures 7 & S12). For Figure 7, only the species with the top five loadings were included for ease of viewing, while Figure S12 contains all NMDS comparisons including all significant species. NMDS demonstrated clear differences between species caught with gill and fyke nets, with 95% confidence ellipses nearly completely separated (Figures 7a & S12a). The taxa with the highest loadings that were primarily driving these differences were *C. commersonii*, *E. lucius*, *M. salmoides*, *Etheostoma* species, and *A. rupestris*. Differences between 12S and 16S, although significant, were more subtle, and were not clearly driven by one or a few species (Figures 7b & S12b). Community composition estimates were highly differentiated between eDNA and traditional gear according to NMDS, with no overlap between 95% confidence ellipses and differences being driven by many species but primarily *Culaea inconstans*, *Cyprinus carpio*, Salmonidae, and *Salvelinus fontinalis* (Figures 7c & S12c). The NMDS of eDNA results grouped by lake side indicated significant spatial differences in community composition, with six net groups from the west side of the lake grouping closely together and away from other net groups, that appeared to be associated with the presence of *Micropterus dolomieu* (Figure 7d & S12d). These net groups were 11, 17, 18, 22, 23, and 24 and were located across the entire west side of the lake (i.e. they were not all clustered in a single area of the lake). Taxa that differentiated samples from the east and west side of the lake included *M. salmoides, Umbra limi, Salmo trutta,* Cyprinidae, and *N. crysoleucas*. The two points below the dam clustered together and were slightly differentiated from other points above the dam in the NMDS based on eDNA results (Figure S12e). The species driving these differences included *S. trutta*, *C. artedi*, *Esox masquinongy*, and *P. caprodes*. The three other NMDS comparisons we performed (eDNA by sampling zone, traditional gear by lake side, and traditional gear by zone) were insignificant (Figure S12f-h).

**Table 2:** Results from analysis of similarities (ANOSIM) comparing estimates of community composition among different sampling methods and areas. P-values <0.05 are in bold. See NMDS plots in Figs. 7 and S12 for visualizations of community compositions.

| Comparison | Anosim R | P-value |
| --- | --- | --- |
| 12S vs 16S | 0.05 | **0.002** |
| gill vs fyke | 0.67 | **0.001** |
| eDNA vs traditional | 0.94 | **0.001** |
| eDNA by zone | 0.07 | 0.053 |
| traditional by zone | 0.02 | 0.238 |
| eDNA by lake side | 0.11 | **0.010** |
| traditional by lake side | 0.03 | 0.755 |
| eDNA above vs below dam | 0.61 | **0.007** |

**Figure 7:**
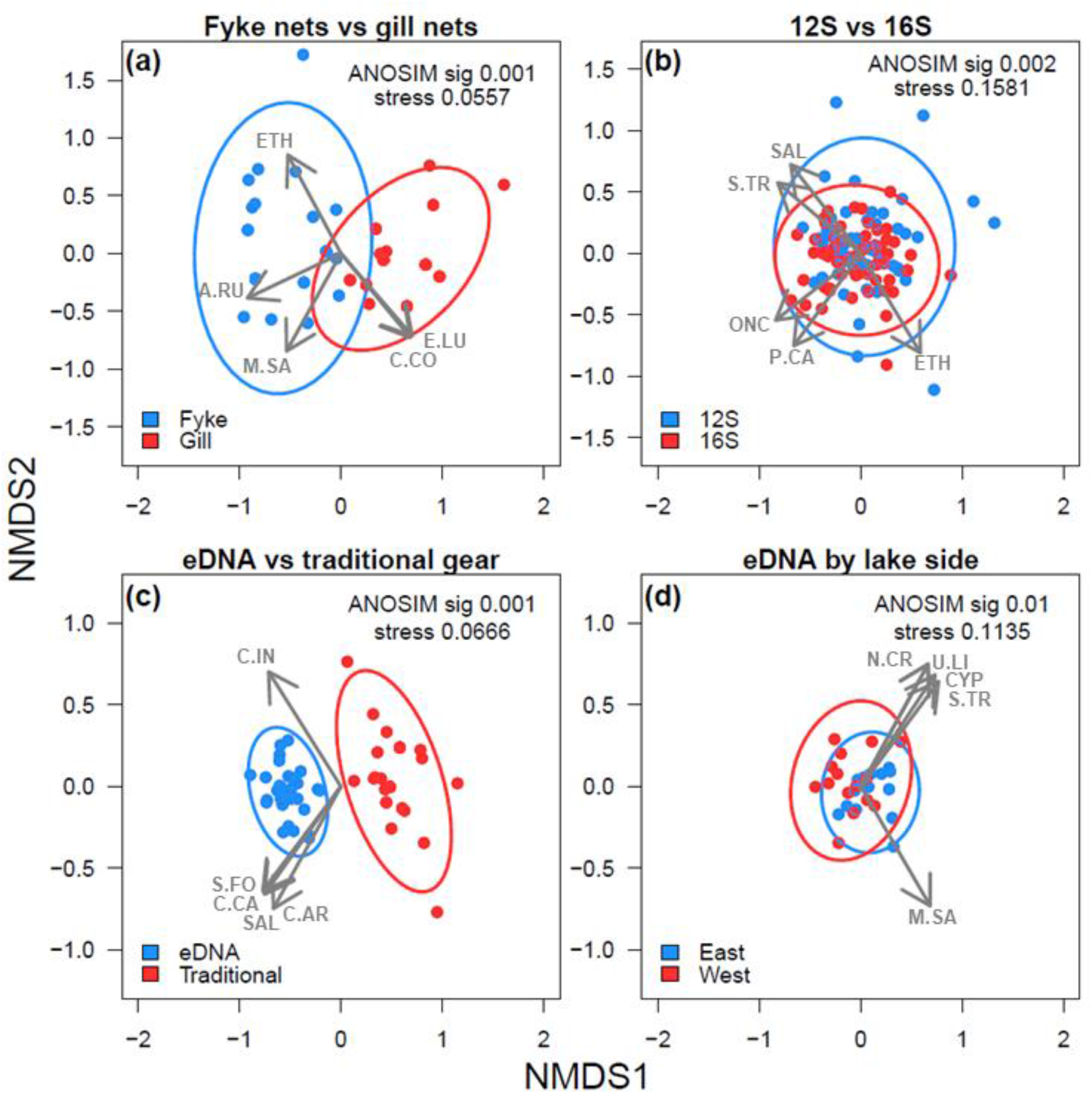
Non-metric multidimensional (NMDS) plots including 95% confidence interval ellipses for four of five significant comparisons (ANOSIM *p*<0.05). Each point represents a net group. For each plot, only the five species with the top loadings were included to facilitate visualizations. 3-letter codes represent the following taxa: A.RU = *Ambloplites rupestris*, C.AR = *Coregonus artedi*, C.CA = *Cyprinus carpio*, C.CO = *Catostomus commersonii*, C.IN = *Culaea inconstans*, CYP = Cyprinidae, E.LU = *Esox lucius*, ETH = *Etheostoma* sp., M.SA = *Micropterus salmoides*, N.CR = *Notemigonus crysoleucas*, ONC = *Oncorhynchus* sp., P.CA = *Percina caprodes*, SAL = Salmonidae, S.FO = *Salvelinus fontinalis*, S.TR = *Salmo trutta*, U.LI = *Umbra limi*. Plot (a) compares fyke nets and gill nets (traditional gear), (b) compares the 12S and 16S genes used in eDNA metabarcoding, (c) compares eDNA metabarcoding with traditional gear overall, and (d) compares the east and west side of Boardman lake from the eDNA data. The only other significant comparison not included here is the comparison between samples taken above and below the dam (S Fig 12e). See supplementary figure 12a-h for all NMDS plots including all significant species loadings.

RDA analysis demonstrated that no variables significantly influenced community composition estimated by traditional gear sampling (Table 3, Figure S13). For the eDNA sampling, however, all four variables (sampling date, zone, lake side, and water temperature) significantly influenced the community composition (Table 3, Figure 8). The most obvious differences in community composition existed between the six points on the west side of the lake (net groups 11, 17, 18, 22, 23, 24) identified in the NMDS analysis and all other points (Figure 8). The RDA analysis indicated that these points were differentiated by zone, lake side, and water temperature and identified similar species driving differences as were discussed above. Two of the environmental variables analyzed here showed high loadings on the same axes and are therefore correlated (lake side and water temperature) but removing one of these variables at a time still produced similar results.

**Table 3:**
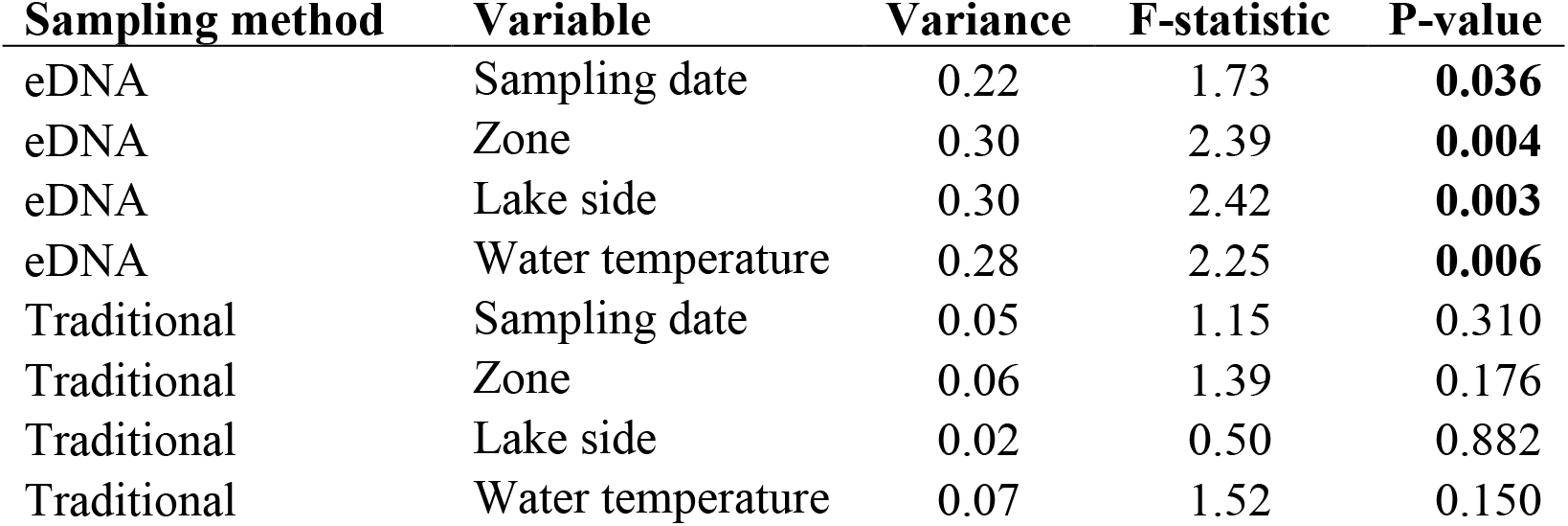
Results from redundancy analysis (RDA) to assess which variables significantly influenced estimates of community composition for eDNA and traditional sampling methods using presence/absence data. Variables assessed were sampling date, zone, lake side, and water temperature. ANOVA P-values <0.05 are in bold.

| Sampling method | Variable | Variance | F-statistic | P-value |
| --- | --- | --- | --- | --- |
| eDNA | Sampling date | 0.22 | 1.73 | **0.036** |
| eDNA | Zone | 0.30 | 2.39 | **0.004** |
| eDNA | Lake side | 0.30 | 2.42 | **0.003** |
| eDNA | Water temperature | 0.28 | 2.25 | **0.006** |
| Traditional | Sampling date | 0.05 | 1.15 | 0.310 |
| Traditional | Zone | 0.06 | 1.39 | 0.176 |
| Traditional | Lake side | 0.02 | 0.50 | 0.882 |
| Traditional | Water temperature | 0.07 | 1.52 | 0.150 |

**Figure 8:**
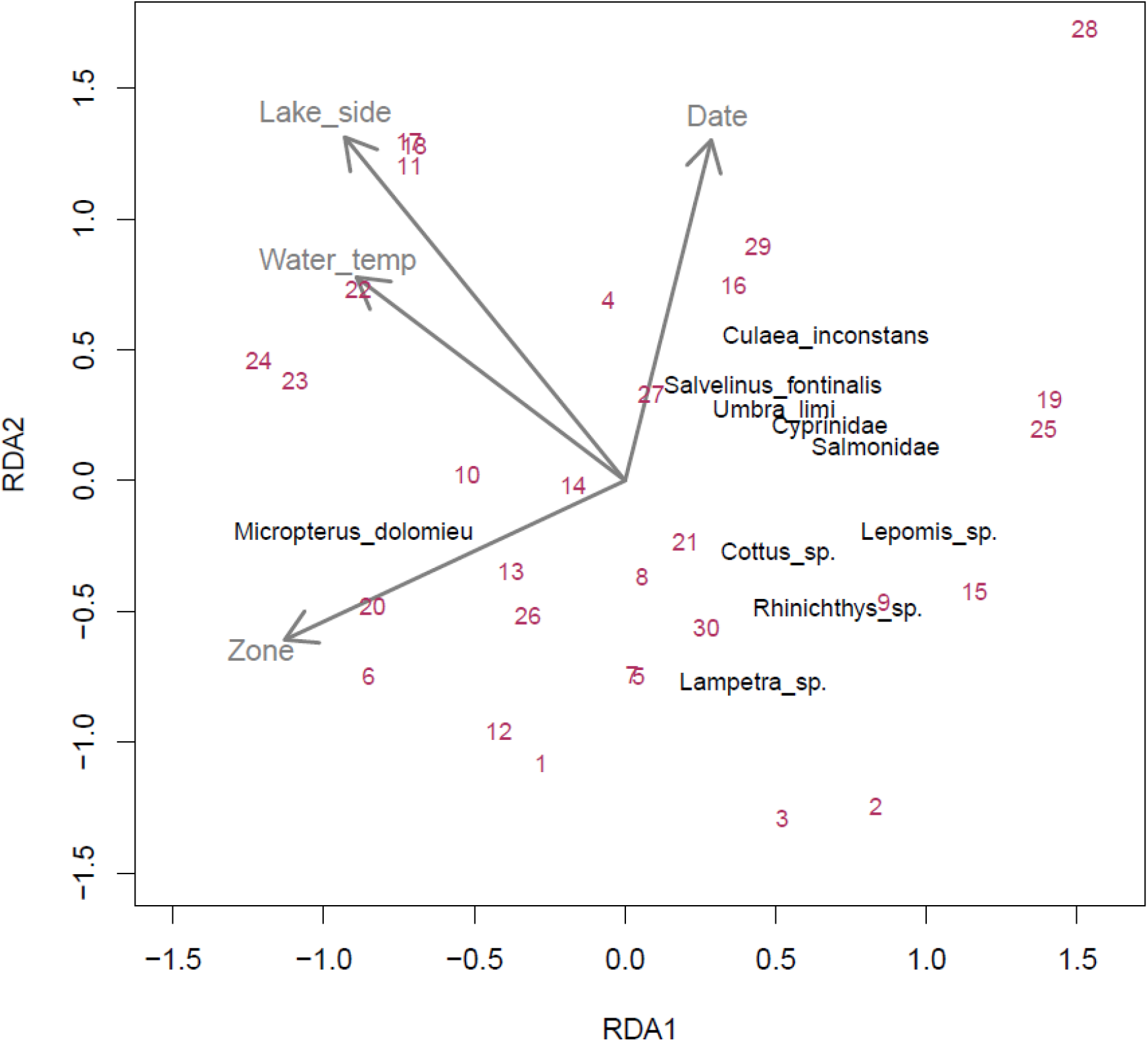
Redundancy analysis (RDA) plot demonstrating how four different variables represented with grey arrows (sampling date, sampling zone, lake side, and water temperature) influenced fish community composition in Boardman Lake measured with eDNA. Maroon numbers are net groups, and only the 10 taxa with the highest scores (black text) were included in the plot to facilitate visualization. All four variables were significant (*p*<0.05).

## Discussion

### Section 1: Comparing eDNA metabarcoding to traditional surveys

In Boardman Lake, more species were detected with 12S and 16S eDNA metabarcoding than traditional survey gears in all net groups. Our eDNA methods detected all the fish species captured in traditional gears, as well as 24 additional taxa. Several species appeared often in the eDNA water samples but were observed in few to none of the nets. For example, *Rhinichthys* sp. (daces), *Lampetra* sp. (lampreys), *Percopsis omiscomaycus* (trout perch), and *Cottus* sp. (sculpins) were present in water samples from over half of the net groups in Boardman Lake but were never observed in nets. *N. melanostomus* (round goby) and *Etheostoma* sp. (darters) were captured in a low number of fyke net sets (5 and 7, out of 30, respectively), but *N. melanostomus* was present in water samples from every net group, and *Etheostoma* sp. were present in 29 out of 30 net groups. Our metabarcoding data suggest that these species are quite common in the lake but are not susceptible to sampling by either of our traditional gears. A common characteristic of these fishes is they are generally sedentary, small-bodied, and/or occupy benthic habitats (Becker, 1983; Scott & Crossman, 1985). These qualities make them difficult to capture with traditional gears, and the species selectivity of these sampling methods is well-established (Zale et al., 2013). Our study thus demonstrates that eDNA metabarcoding can overcome problems with selectivity and the difficulty of capturing rare, small, and/or benthic fish species with traditional gears.

We also observed a clear difference in the species selectivity between our two traditional gear types. For example, *A. rupestris* was the most common species in fyke nets (68.2% of all fish), followed by *P. flavescens* (9.7% of all fish). In gill nets, however, *A. rupestris* only accounted for 14.6% of all fish caught. Our species accumulation curve demonstrated that at a low quantity of samples, gill nets detect more species than fyke nets, but with increasing samples, fyke nets detected more species overall. This could be because gill nets tended to catch more fish on average but they were larger, more pelagic fishes. Our fyke nets had a smaller mesh size and were set against the shoreline. Therefore, they may be more likely to capture smaller, littoral, and/or benthic species. For example, we observed *Etheostoma* sp. and *N. melanostomus* in fyke nets but never in gill nets. These results further demonstrate that multiple gear types are necessary for accurate fish community assessments using traditional gear and highlight the advantage of lower species selectivity with eDNA.

Overall, the number of sequence reads and fish biomass/catch in Boardman Lake was not correlated. Several studies have demonstrated that read counts can correlate to fish abundance (e.g. Hänfling et al., 2016; Lacoursière-Roussel et al., 2016; Thomsen et al., 2016), however many of these studies obtained accurate population estimates using standardized sampling methods and/or long-term data. These data were unavailable for Boardman Lake, and we hypothesized that the number of fish encountered in a net should be relatively correlated to their local abundance, which may be incorrect. It is also possible that the lack of correlation could be due to error introduced during the eDNA sample collection and laboratory processing, or even the introduction of genetic material from upstream. It is well understood that subsampling, primer bias, and PCR inhibition can lead to a lack of correlation between eDNA sequence reads and true abundance (Deiner et al., 2017; Goldberg et al., 2016; Porter & Hajibabaei, 2018), although we attempted to pre-emptively address these issues through replication, use of vetted and published assays, and inhibitor removal. Additionally, ecological factors such as high numbers of gametes in the water during spawning season could impact correlations. Both *P. flavescens* and *C. commersonii* were common in the system, and if they had recently spawned, amplification of eDNA from their gametes could have inflated read numbers. In general, our data suggest that the proportion of detections across all eDNA water samples may be a better predictor of relative abundance than read counts, as higher occurrences in samples are likely a result of species being common in the system. For example, *P. flavescens* was caught with traditional gear in 29 out of 30 net groups and was present in water samples in 30 out of 30 net groups, suggesting it is likely one of the most common species in Boardman Lake.

### Section 2: Detection of spatial heterogeneity of fish distribution with eDNA metabarcoding

We were able to detect differences in fish community composition between the west and east sides of Boardman Lake with multivariate analyses of our eDNA metabarcoding data, but not with the traditional sampling data. This result suggests that our eDNA methods were sensitive enough to detect some spatial heterogeneity in fish distributions across relatively short distances (500 meters between the east and west shores). There were six net groups from the west side of the lake that appeared to be driving the differences we observed. Based on our NMDS comparison of eDNA data by lake side, *M. dolomieu* (smallmouth bass) appears to be the main species causing these six points on the west side of the lake to differ from the others. It is well-known that fish have inherently patchy distributions due to spatial differences in habitats as well as variations in both abiotic and biotic factors (Romare et al., 2003; Smokorowski & Pratt, 2007; Weaver et al., 1997). Therefore, our results suggest that there is likely some physical or biological characteristic of the habitat on the west side of the lake that *M. dolomieu* prefer. This species is known to congregate in areas with dense underwater woody debris (Becker, 1983; Scott & Crossman, 1985), but this was not a variable that we assessed as part of this study. Lake bathymetry was also not a variable that we explicitly measured, but in the field, we observed that the lake bottom on the west side dropped off more abruptly. This physical attribute may have potentially affected the presence or distribution of fish in those locations.

Because of the inherent patchiness of fish distributions within a system, it is reasonable that with enough samples, eDNA metabarcoding could detect some spatial differences. This further emphasizes the importance of collecting numerous eDNA samples across multiple locations to fully characterize the fish community within a lake. However, the analysis we conducted was post-hoc and our study was not designed to test hypotheses about which environmental variables might be correlated with fish distributions. To test such hypotheses, future metabarcoding studies could incorporate detailed habitat surveys and pair them with intensive water sampling. For example, researchers could include water quality measurements like DO, temperature, and turbidity; physical variables like substrate composition, bathymetry, cover, and woody debris; and biological variables such as aquatic vegetation and zooplankton abundance/composition.

### Section 3: Obtaining robust eDNA metabarcoding results

Using both 12S and 16S for our eDNA metabarcoding analysis provided a more comprehensive characterization of the fish community than if we had used a single gene (e.g. Evans et al., 2017; Sard et al., 2019; Shaw et al., 2016; Stat et al., 2017). Both genes consistently detected the most common species in the system, but rarer species were not always detected with both genes in a given sample. For example, in some net groups *M. dolomieu* was detected just with 12S, in others it was detected only with 16S, and sometimes it was detected with both genes. Other rare species showed similar patterns, but no species that were resolved with both genes consistently amplified at one gene and not the other, indicating that variation in detections is likely a result of subsampling of extracted DNA or stochasticity in PCR rather than inherent differences in species-specific detection probabilities (Deiner et al., 2017; Kebschull & Zador, 2015). Taxonomic resolution varied between the two genes, with each classifying various taxonomic groups differently. For example, 12S contained sufficient variation to classify species within the *Etheostoma* genus (darters) while 16S could not. It is important to note that because of the difficulty of parsing some taxa to species, we sometimes assigned back to genus or family, and this could underestimate the total number of species present in the lake. For example, there are likely several species present within our Cyprinidae assignment for 12S, but we were not confident in those species assignments due to low sequence variation for that particular gene within that taxonomic group. Therefore, if a study requires high taxonomic resolution for a particular taxonomic group like Salmonidae or Cyprinidae, it would be beneficial to utilize additional genes that contain sufficient variation among taxa.

Our study demonstrates the importance of taking many water samples across time and space, especially to detect rare species. For example, *Pomoxis nigromaculatus* and *L. cornutus* were detected in water samples from only two out of 32 net groups, and *P. notatus* and *P. caprodes* were only detected in one. Taking samples in triplicate was also essential. We sometimes observed that even common species were only detected in one replicate out of three, meaning if we had only collected a single water sample, that species likely would have gone undetected. It is well known that PCR bias and random events from sample collection to bioinformatics can lead to false negatives, even when extreme care is taken (Goldberg et al., 2016; Porter & Hajibabaei, 2018; Thomsen & Willerslev, 2015). This emphasizes the importance of using multiple water sample replicates per sample site, which has also been suggested by other studies (e.g. Evans et al., 2017; Shaw et al., 2016; Willoughby et al., 2016), but has not consistently been implemented (Dickie, 2018). We show that having many samples with replicates allows much higher taxonomic coverage, increased likelihood of detecting rare species, and overall high confidence in results.

Our metabarcoding data provided some useful information beyond our main study objectives. For example, we were able to compare species detected below the Union Street Dam to those in Boardman Lake. Community compositions were relatively similar between locations, but several species appeared to be more common below the dam, including *C. artedi* (cisco or lake whitefish), *P. marinus* (sea lamprey), and *Alosa* sp. (likely alewife). It is logical that these species common in the Great Lakes would be present in the lower river section connected to Lake Michigan. We also explored the potential of detecting DNA transported downstream from the upper Boardman River into Boardman Lake. It is well-documented that rivers transport eDNA downstream (e.g. Civade et al., 2016; Deiner et al., 2016; Jane et al., 2015; Pont et al., 2018), however we did not detect any spatial patterns that would suggest a higher concentration of DNA from lotic species (e.g. brook trout *S. fontinalis*) where the river enters the lake. Based on the consistent distribution of eDNA around the lake, it is likely that most of the DNA we detected in water samples were from fish residing in the lake. However, the ecology and decomposition rates of eDNA in this system are unknown and we did not have upstream water samples for comparison, therefore we cannot make concrete conclusions about the origin of fish DNA in the lake. Finally, while this research was not focused on non-fish species, we were intrigued to discover that several other animals amplified in our eDNA water samples. These included a variety of birds (duck, goose, swan, cormorant, chicken), mammals (squirrel, mole, raccoon, beaver, red fox, and black bear), and snapping turtle. These non-target detections provide useful information on community composition that could be mined in future studies.

### Section 4: Quality control with eDNA metabarcoding

Our study emphasizes the importance of meticulous consideration of all taxonomic assignments, knowledge of species phylogenies, and understanding of species distributions and potential for hybridization. We developed a fully comprehensive and curated reference database and passed our ASVs through conservative filtering parameters. Some ASVs matched to species known not to be present in the region, for example *Sander lucioperca* (zander) which exists in Europe and Asia but is related to native species *S. vitreus* and *S. canadensis*. These foreign taxa known to not be in the region were removed to avoid any impossible matches and create a more refined reference database. After filtering, rather than simply accepting the top match, each retained ASV assignment was assessed individually before being accepted as valid. Some assignments were straightforward; for example, one ASV had three matches, all of which were variants of *C. commersonii*. However, one ASV had a 100% match to *Ameiurus melas*, a 99.3% match to *Ameiurus nebulosus*, and a 98.6% match to *Ameiurus natalis*. It would be easy for a researcher to observe the first 100% match and promptly assign this ASV to *A. melas*, but because there was also high similarity to two other species, we were not confident that this ASV was definitively *A. melas*. Therefore, we assigned that ASV to “*Ameiurus* sp.” We also made these more conservative taxonomic assignments in instances where there is known low sequence variation within a group (e.g. Salmonidae with 12S and *Cottus* with both genes), or when there was a potential for species hybridization (e.g. in the genus Lepomis; Avise & Saunders, 1984). As mentioned above, these broader assignments may be undercounting true species richness within the system, but we felt it was important be completely confident in each taxonomic assignment.

In addition to ensuring confidence in taxonomic assignments, we chose to be quite conservative when determining if a species was truly present in a sample. Because we commonly detected hundreds or thousands of sequence reads for each species in our water samples, we set our minimum read requirement as 10. This is more conservative than other metabarcoding studies, which have set their minimum required reads as 2 or 3 (Balasingham et al., 2018; Olds et al., 2016). There is clearly the potential to lose some true detections with our higher cutoff, but once again we felt it was important to be conservative and confident in our detections. The fact that we observed 40 taxa even with this conservative data filtering suggests that our metabarcoding assay was sensitive and our methods were robust.

Most of our field negative controls had low contamination, proving that our sterilization techniques were effective. In instances where contamination did exist, it was often from one of the most common species in the Boardman system according to net sampling, suggesting that the contamination likely occurred during sample collection or filtering rather than during eDNA extractions or pre- and post-PCR. The low contamination in our laboratory positive and negative controls also verify that contamination from the laboratory, as well as cross contamination among plates and wells, was minimal. Surprisingly few studies report sources of contamination or when it occurred during the process, but Harper et al. (2019) found that most contamination occurred during sampling and PCR, and Furlan et al. (2020) found that some contamination occurred through all stages of sampling and processing, but mostly during PCR. Because we subtracted the maximum number of reads in controls from all read counts in associated samples, we were confident that the remaining reads were true detections. Our results emphasize the importance of using field and lab controls to track any potential contamination, and accounting for any contamination during downstream bioinformatics.

### Section 5: Guidance for future studies and conclusions

In conclusion, we demonstrated that surface water eDNA detected roughly three times as many unique fish taxa than traditional methods in a small temperate lake. Our eDNA metabarcoding assay characterized the fish community much more comprehensively than traditional net sampling, providing evidence that metabarcoding can overcome some of the problems associated with traditional sampling and provide a more sensitive approach to describe community composition. eDNA sample collection is fast and simple, and laboratory and analytical processes are becoming more streamlined. We suggest that eDNA be considered for estimating fish species richness, as the personnel and infrastructure required to collect eDNA samples is minimal compared to traditional techniques. This decrease in cost should allow researchers and managers to sample more areas, sample more frequently, and conduct more intensive sampling, which could provide valuable information to inform management and conservation of aquatic systems. However, eDNA metabarcoding is not expected to replace standard fisheries techniques for many applications such as studies on population dynamics, as eDNA does not distinguish between live or dead organisms, nor provide information on age, sex, or sizes of fish. Therefore, eDNA metabarcoding is likely best used to complement traditional fisheries methods, rather than replace them. Our results reveal the utility of collecting many samples across time and space, and the usefulness of collecting multiple replicates for each sampling event. Additionally, we demonstrated that fine-scale heterogeneity can be detected using eDNA. Fish community structure may not be homogenous across water bodies and collecting spatially representative samples is critical. Finally, this study demonstrates the importance of understanding species phylogenies and taking a conservative approach when classifying taxa and assigning species to achieve robust results.

## Acknowledgments

This manuscript is contribution #3 of FishPass (http://www.glfc.org/fishpass.php). FishPass is the capstone to the 20-year restoration of the Boardman (Ottaway) River, Traverse City, Michigan. The mission of FishPass is to provide up- and down-stream passage of desirable fishes while simultaneously blocking or removing undesirable fishes, thereby addressing the connectivity conundrum. We are grateful to the primary project partners: Grand Traverse Band of Ottawa and Chippewa Indians, Michigan Department of Natural Resources; U.S. Army Corps of Engineers; U.S. Fish and Wildlife Service, and the U.S. Geological Survey. We also extend sincerest thanks to the primary partner, the City of Traverse City. Without the city’s support and the vision of the city commission, FishPass would not have been possible. Funding for this contribution came from the Great Lakes Restoration Initiative and the Great Lakes Fishery Commission. We thank Reid Swanson, Heather Hettinger, and Kelly Boughner for assistance with sampling. Jacek Maselko and Mike Spear provided valuable editorial commentary on this manuscript.

## Data Archiving Statement

Upon acceptance, raw sequence reads (*fastq* format) used in this research along with corresponding metadata will be archived in the NCBI sequence read archive using the publicly accessible Genomic Observatories Metadatabase (GEOME, http://www.geome-db.org/)

## Author Contributions

WL and RG designed the study. RG performed field sampling. NS provided laboratory and sampling protocols, which were refined by KG. RG and KG performed laboratory analysis. Data analyses were conducted by RG and WL with input from YS and NS. RG and WL wrote the manuscript and all other authors provided feedback and manuscript edits.

## Supplementary Materials

**Table S1:** Sampling metadata for all eDNA water samples collected in Boardman Lake and below Union Street Dam in the Boardman River.

**Table S2:** Number of unique taxa in each net group for each sampling method, which included eDNA (12S and 16S genes) and gill and fyke net sampling, with mean, standard deviation, and standard error. A net group consisted of one gill net and one fyke net with their corresponding water samples (3 replicates per net set).

**Table S3:** Counts and percentages of catches for all species caught in fyke nets and gill nets in Boardman Lake.

**Figure S1:** Boxplot of sequence read counts for 12S and 16S per water sample from Boardman Lake and below-dam Boardman River. The average number of reads in 12S was significantly higher than that of 16S (*p*<0.001).

**Figure S2:** Heatmap illustrating the number of replicates in which a taxon was present in each net group for 12S. A net group in Boardman Lake consisted of one gill net and one fyke net, each with 3 corresponding water samples, therefore the maximum number of detections for net groups 1-30 (Boardman Lake) is 6. Net groups 31 and 32 (below Union Street Dam in the Boardman River) consisted of 9 water samples and had no corresponding net sets, therefore anything between 6 and 9 detections is labeled (>6) for these two “net groups”.

**Figure S3:** Heatmap demonstrating the number of replicates in which a taxon was present in each net group for 16S. A net group in Boardman Lake consisted of one gill net and one fyke net, each with 3 corresponding water samples, therefore the maximum number of detections for net groups 1-30 (Boardman Lake) is 6. Net groups 31 and 32 (below Union Street Dam in the Boardman River) consisted of 9 water samples and had no corresponding net sets, therefore anything between 6 and 9 detections is labeled (>6) for these two “net groups”.

**Figure S4:** Correlation between the number of unique taxa detected and the number of reads for (a) 12S and (b) 16S for each eDNA water sample replicate from Boardman Lake and Boardman River. Spearman’s correlation coefficient r_s_ was = 0.47 and 0.41 for 12S and 16S, respectively, and p<0.001 for both genes.

**Figure S5:** Correlation between the number of instances a taxon was detected using 12S and 16S in eDNA samples. Points represent all detected taxa and data include both Boardman Lake and below-dam Boardman River water samples. Pearson’s correlation coefficient r was 0.91 and *p*<0.001.

**Figure S6:** Number of fish DNA reads in negative field controls associated with water samples from Boardman Lake and below Union St Dam in the Boardman River for 12S and 16S genes. Three large outliers were removed prior to analysis. The number of reads in negative controls for each gene did not significantly differ (*p*=0.362).

**Figure S7:** Correlation of species contamination in negative field controls from Boardman Lake and below Union St Dam in the Boardman River detected with 12S and 16S genes. Each point represents a fish species detected in negative controls. Three large outliers were removed prior to analysis. Pearson’s correlation coefficient r was 0.75 and *p*<0.001.

**Figure S8:** Boxplot of average number of fish caught per net for fyke and gill nets in Boardman Lake. Gill nets caught significantly more fish on average than fyke nets (*p*=0.004).

**Figure S9:** Heatmap indicating taxa detected in each net group using traditional net sampling in Boardman Lake. Red indicates instances in which a species was only observed in a gill net, blue indicates species that were only observed in a fyke net, purple indicates when a species was detected in both net types, and grey represents no observation of that species in either net.

**Figure S10:** Pearson correlations between the number of fish of a particular species in a net and the number of DNA reads for that species in corresponding eDNA water samples from Boardman Lake. Correlations were performed only for the five most common species (*Ambloplites rupestris*, *Catostomus commersonii*, *Esox lucius*, *Sander vitreus*, and *Perca flavescens*) for each net type and gene. Each point represents an individual water sample. All correlations were insignificant except for *Perca flavescens* in fyke nets with 12S (*p*=0.01).

**Figure S11:** Pearson correlations between the biomass (g) of a particular species in a net and the number of DNA reads for that species in corresponding eDNA water samples from Boardman Lake. Correlations were performed only for the five most common species (*Ambloplites rupestris*, *Catostomus commersonii*, *Esox lucius*, *Sander vitreus*, and *Perca flavescens*) for each net type and gene. Each point represents an individual water sample. No correlations were significant at α=0.05.

**Figure S12:** Plots of all eight non-metric multidimensional (NMDS) comparisons performed. Each point represents a net group. All taxa that were significant (*p*<0.05) in each analysis were included. 95% confidence interval ellipses were drawn around groups for all comparisons except 12e due to the downstream group only having two data points.

**Figure S13:** Redundancy analysis (RDA) plot demonstrating how four different variables represented with grey arrows (sampling date, sampling zone, lake side, and water temperature) influenced fish community composition in Boardman Lake estimated with traditional net sampling. Maroon numbers are net groups, and only the 10 taxa with the highest scores were included in the plot. No variables were significant in this analysis (all with *p*>0.05).

**Supplementary File 1**. Table with number of sequence reads for each taxa in each water sample replicate from Boardman Lake and Boardman River for the 16S gene, including field negative controls, lab negative controls, and lab positive controls. Metadata for each sample is included. Note that contamination in controls has not been subtracted from sample read counts listed here. See Supplementary file 3 for read counts in each sample after accounting for contamination in controls.

**Supplementary File 2**. Table with number of sequence reads for each taxa in each water sample replicate from Boardman Lake and Boardman River for the 12S gene, including field negative controls, lab negative controls, and lab positive controls. Metadata for each sample is included. Note that contamination in controls has not been subtracted from sample read counts listed here. See Supplementary file 3 for read counts in each sample after accounting for contamination in controls.

**Supplementary File 3**. Table with number of sequence reads for each taxa in each water sample replicate from Boardman Lake and Boardman River for the 16S gene after accounting for contamination in field and lab controls. Metadata for each sample is included.

**Supplementary File 4**. Table with number of sequence reads for each taxa in each water sample replicate from Boardman Lake and Boardman River for the 12S gene after accounting for contamination in field and lab controls. Metadata for each sample is included.

