## Supplementary figures and images for "eDNA metabarcoding outperforms traditional fisheries sampling and reveals fine-scale heterogeneity in a temperate freshwater lake"

### Figure S1

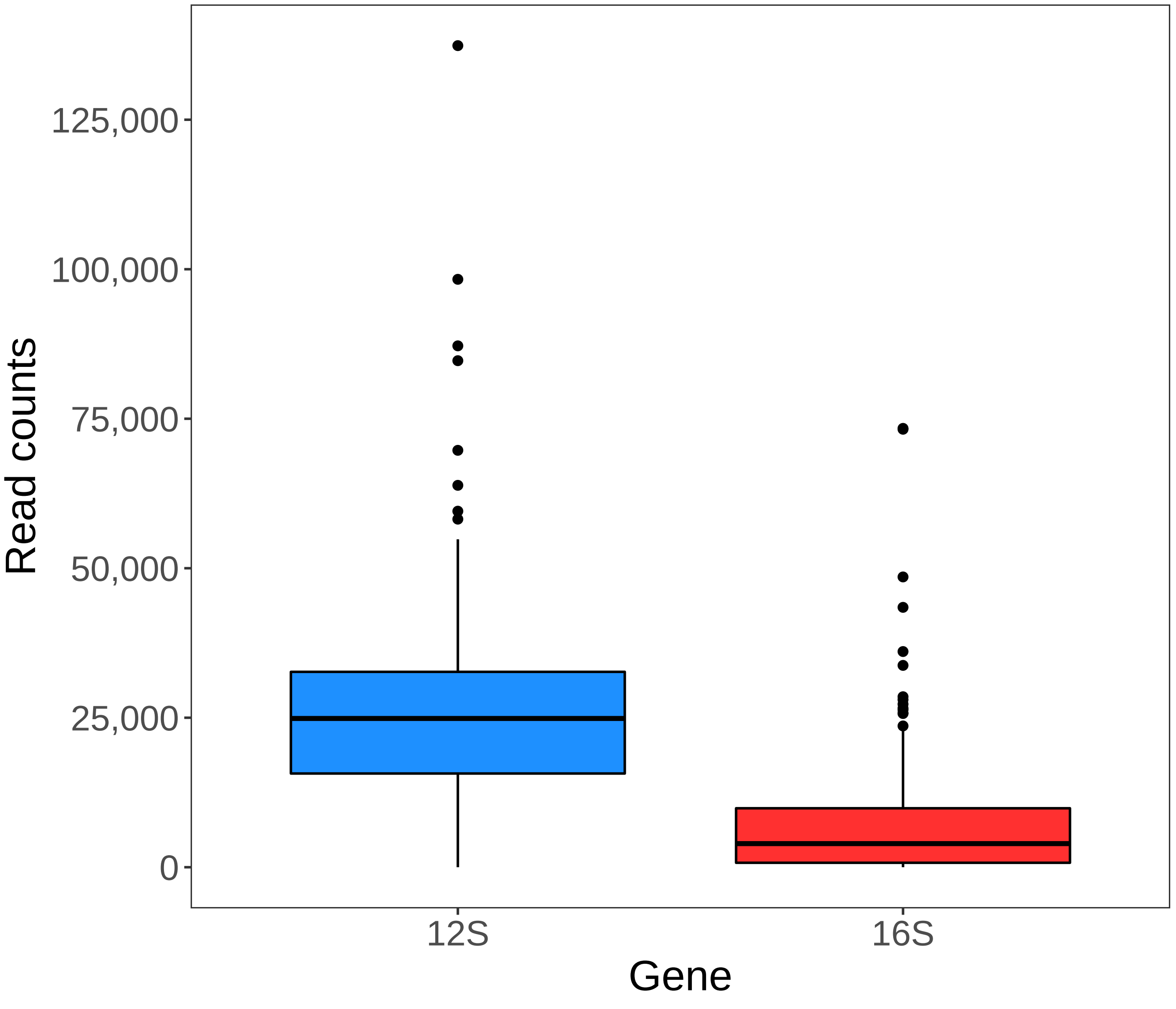

### Figure S2

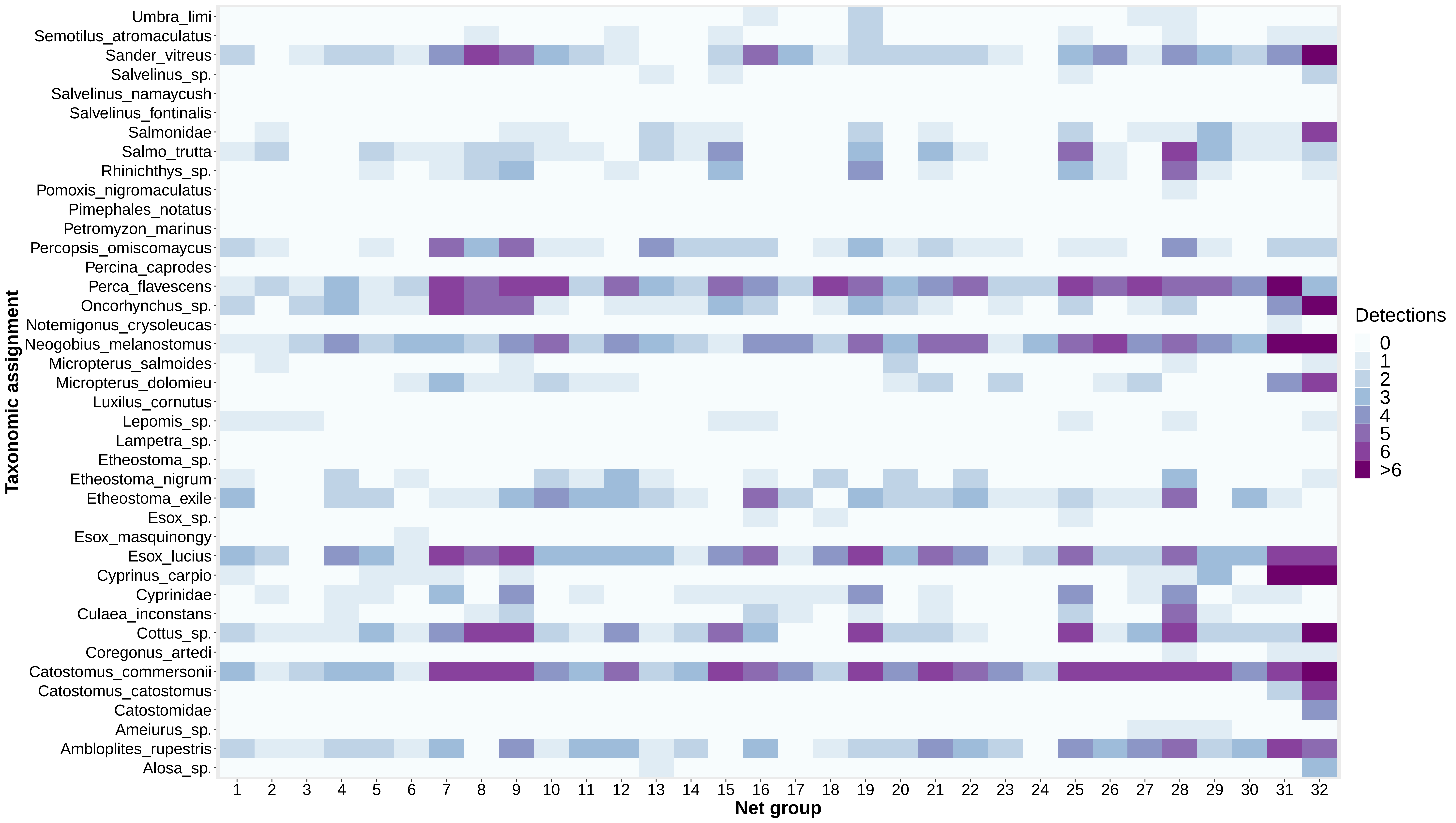

### Figure S3

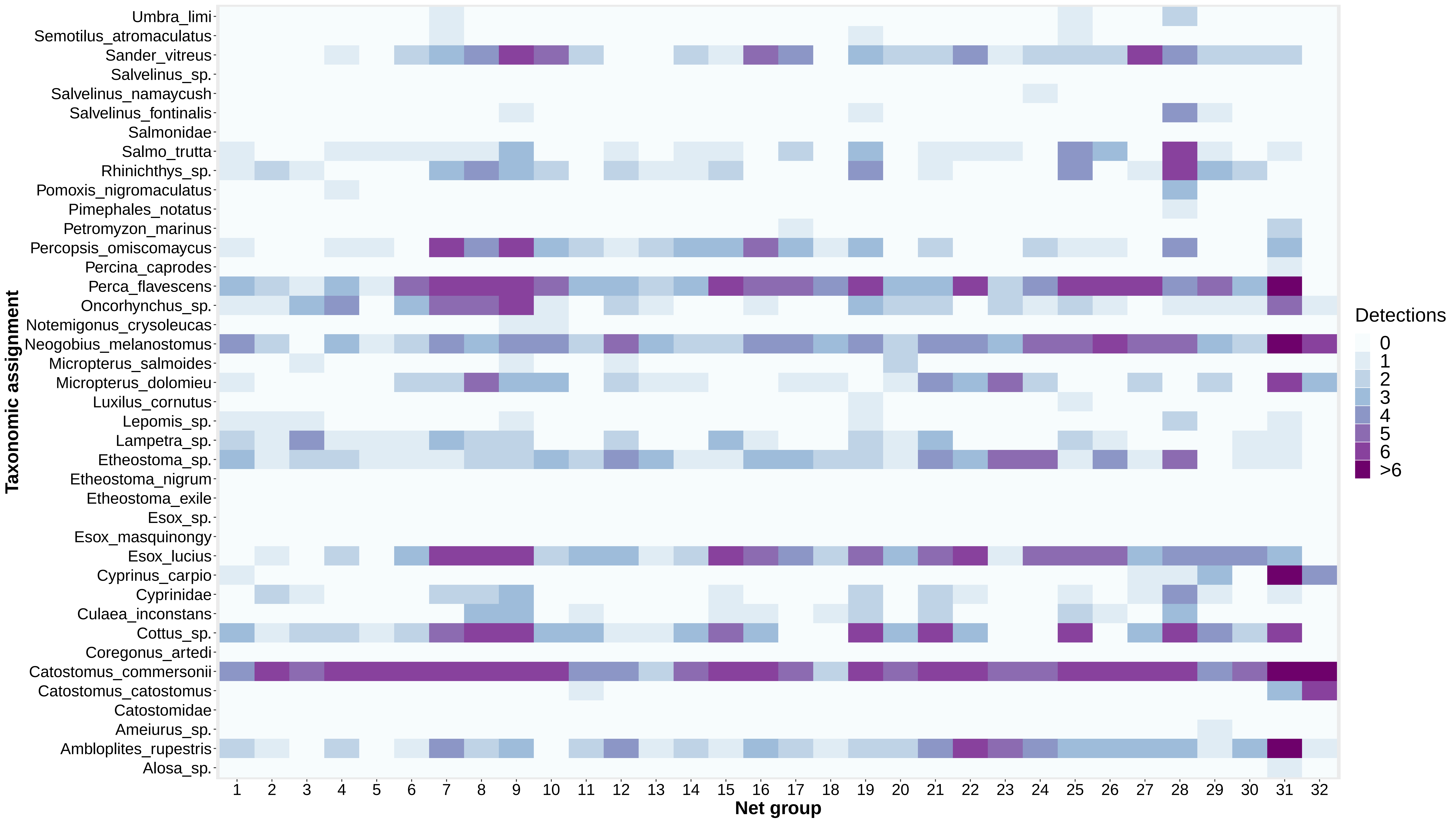

### Figure S4

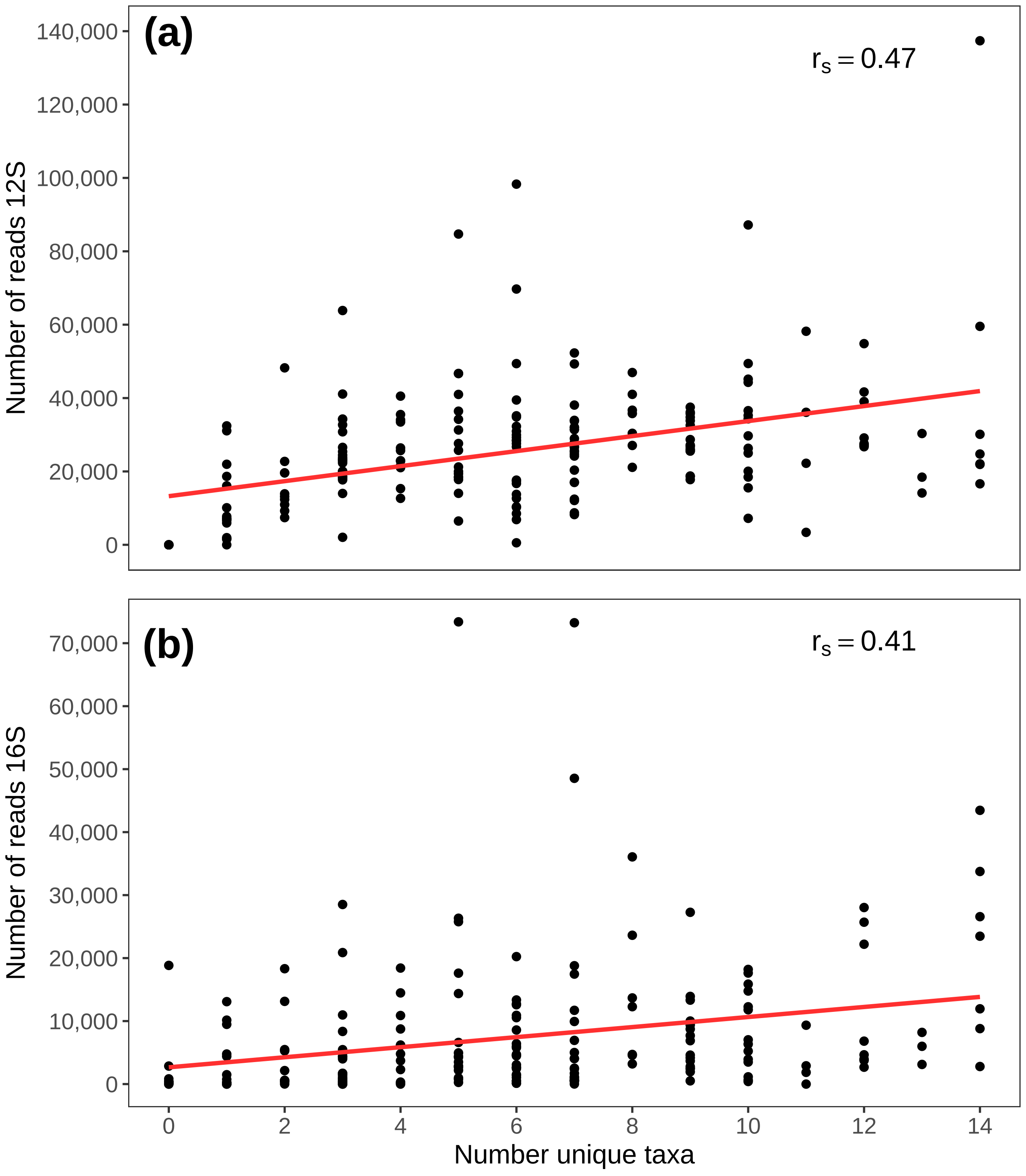

### Figure S5

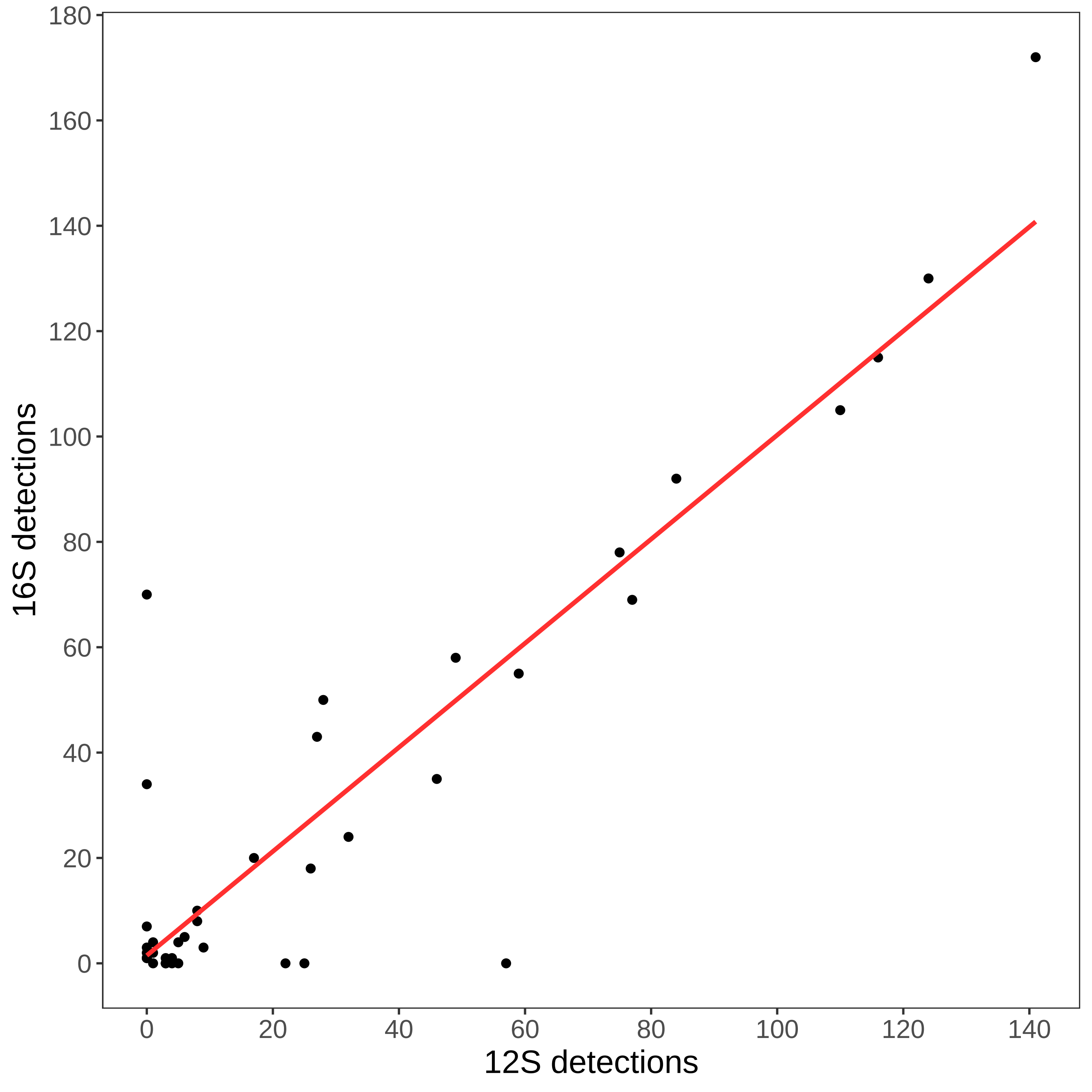

### Figure S6

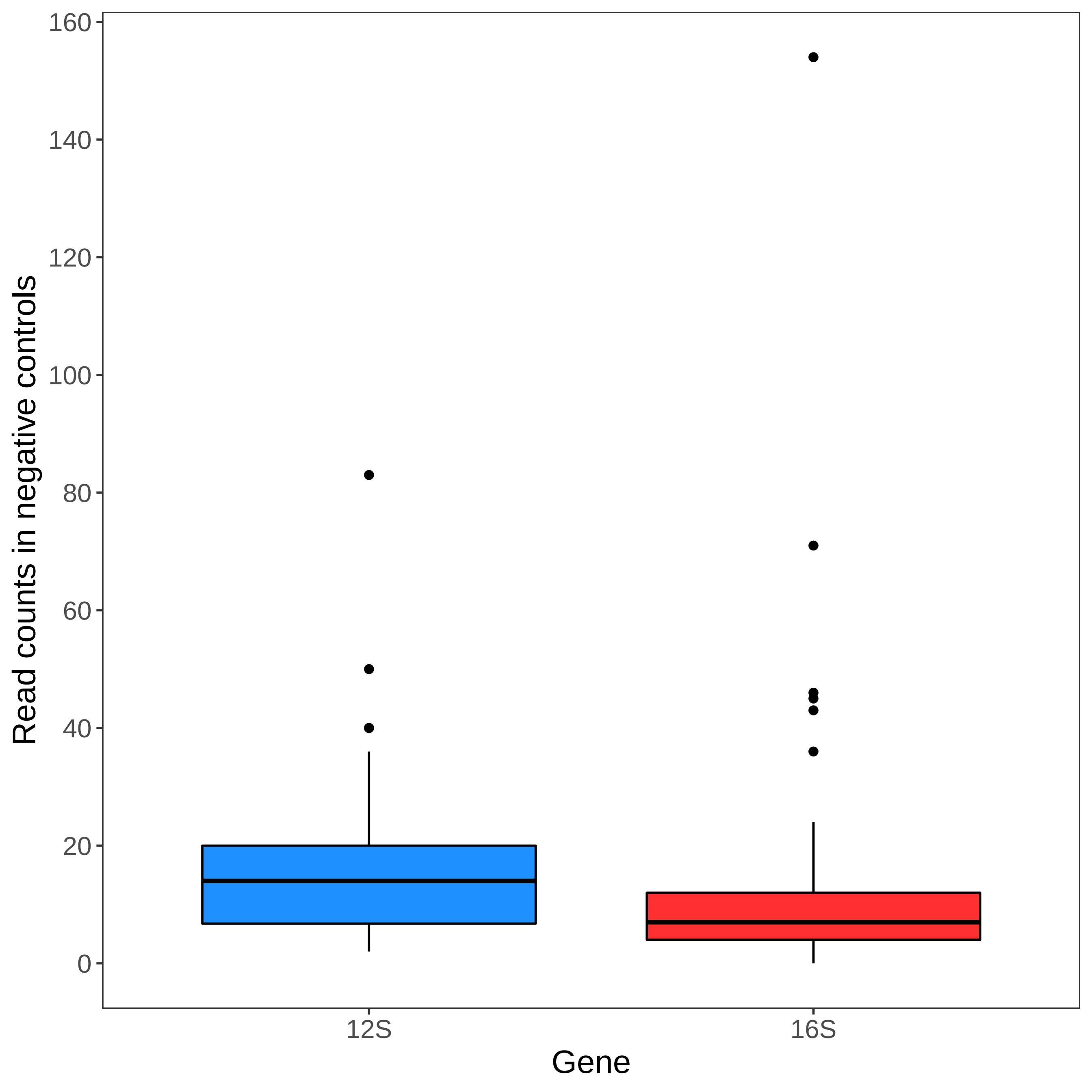

### Figure S7

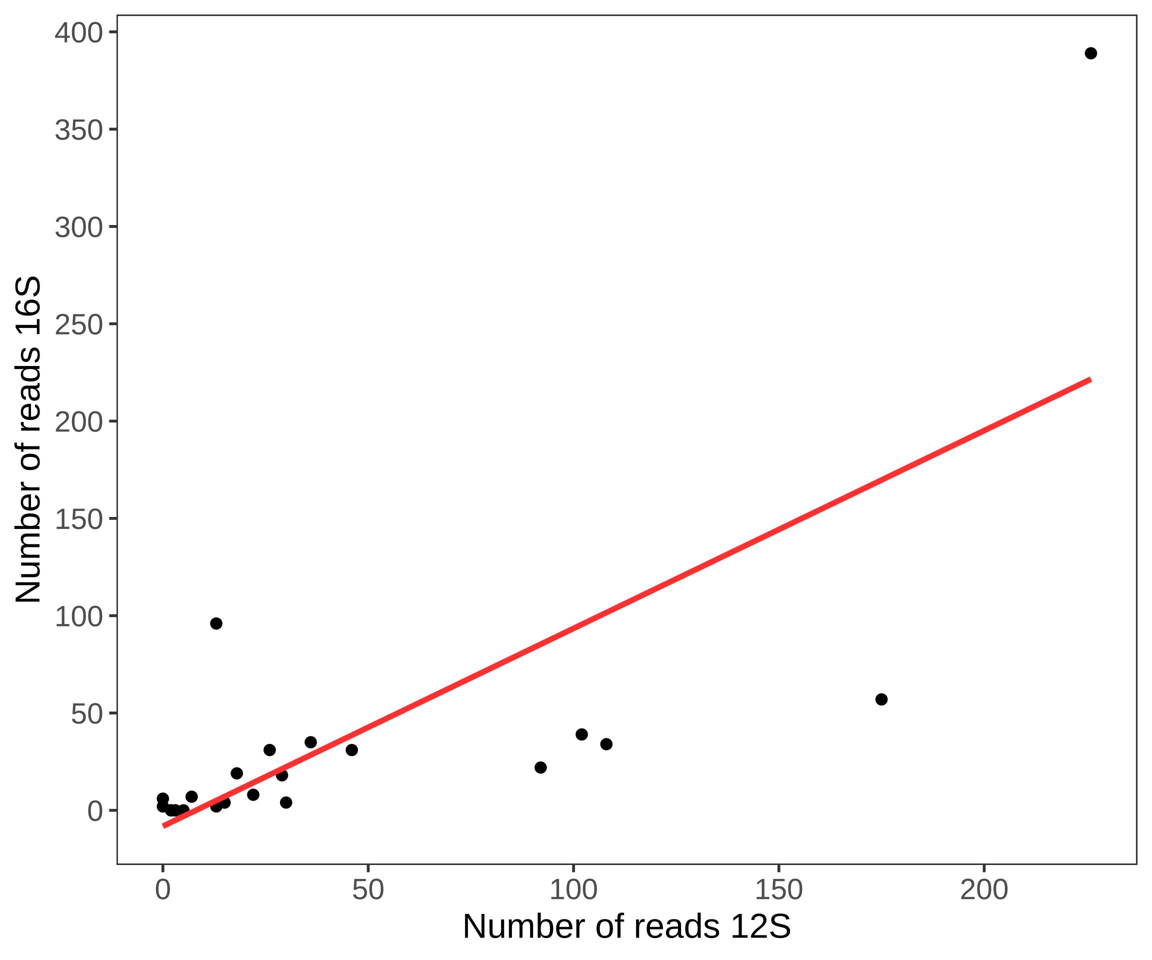

### Figure S8

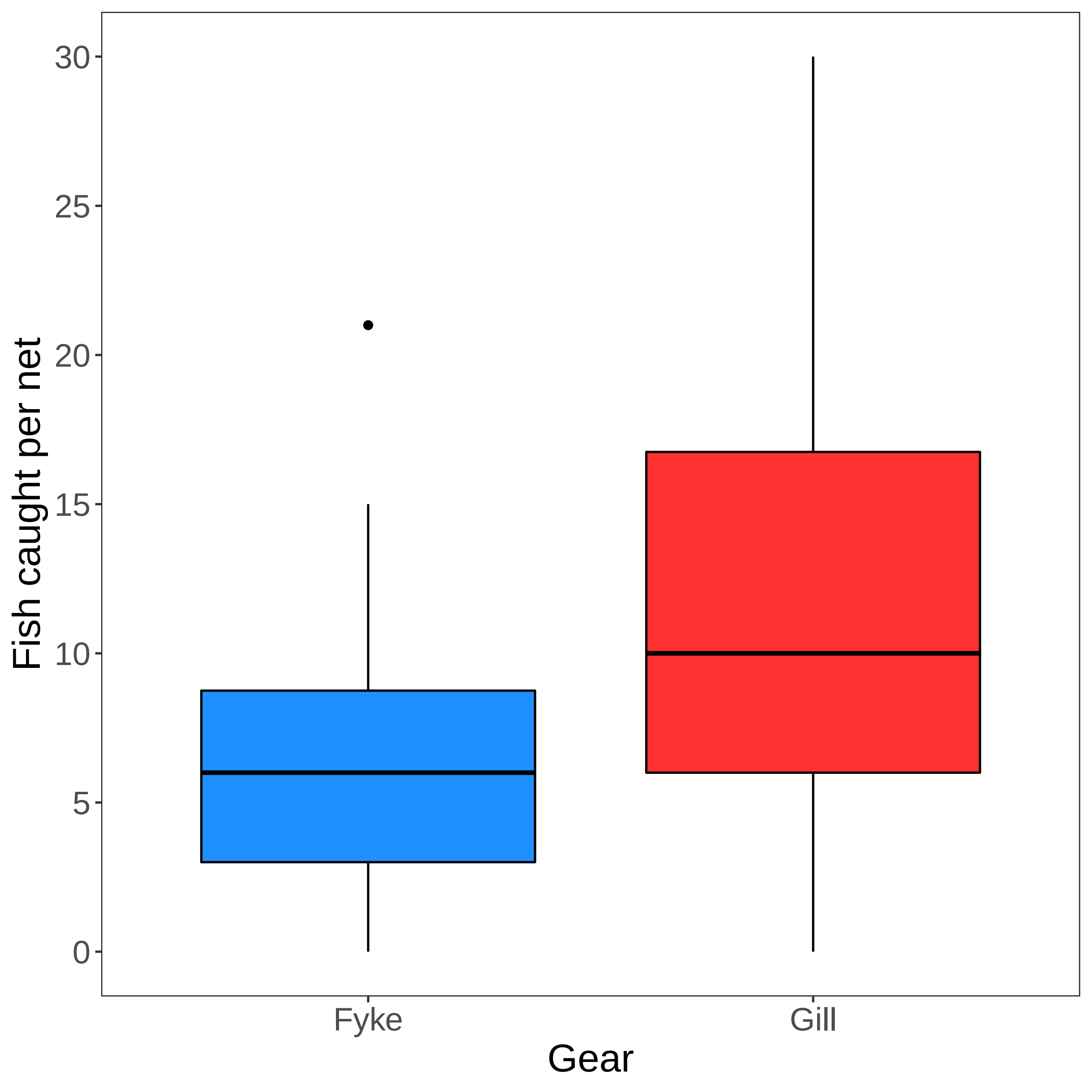

### Figure S9

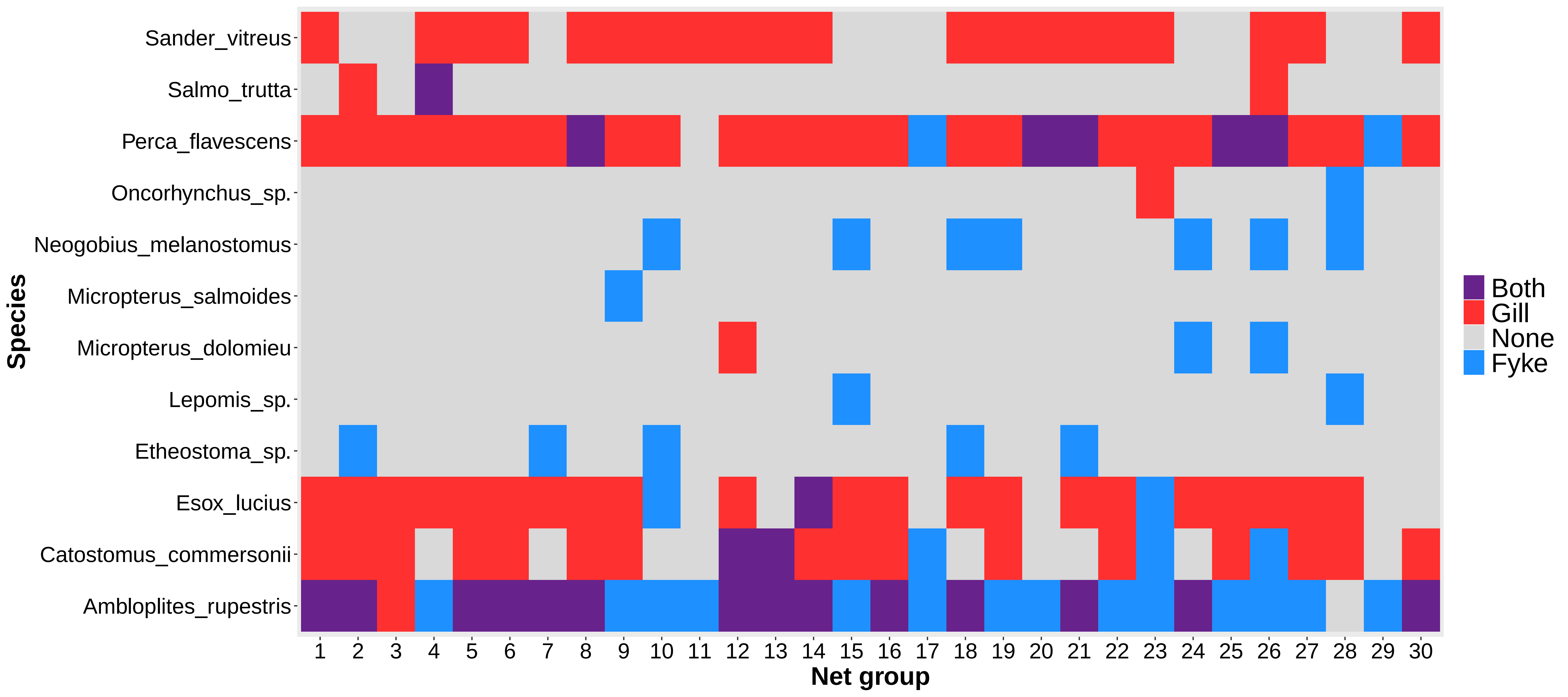

### Figure S10

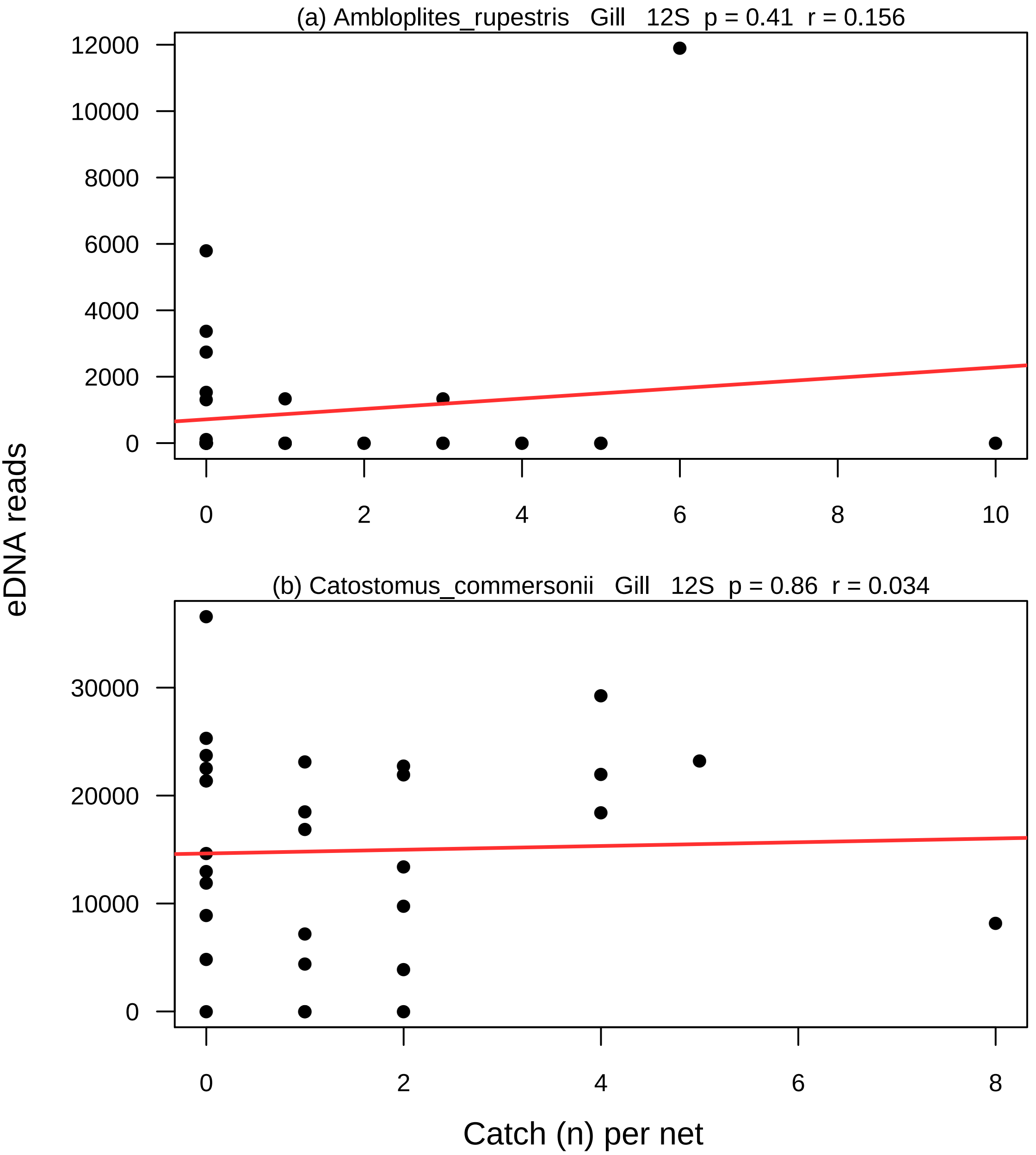

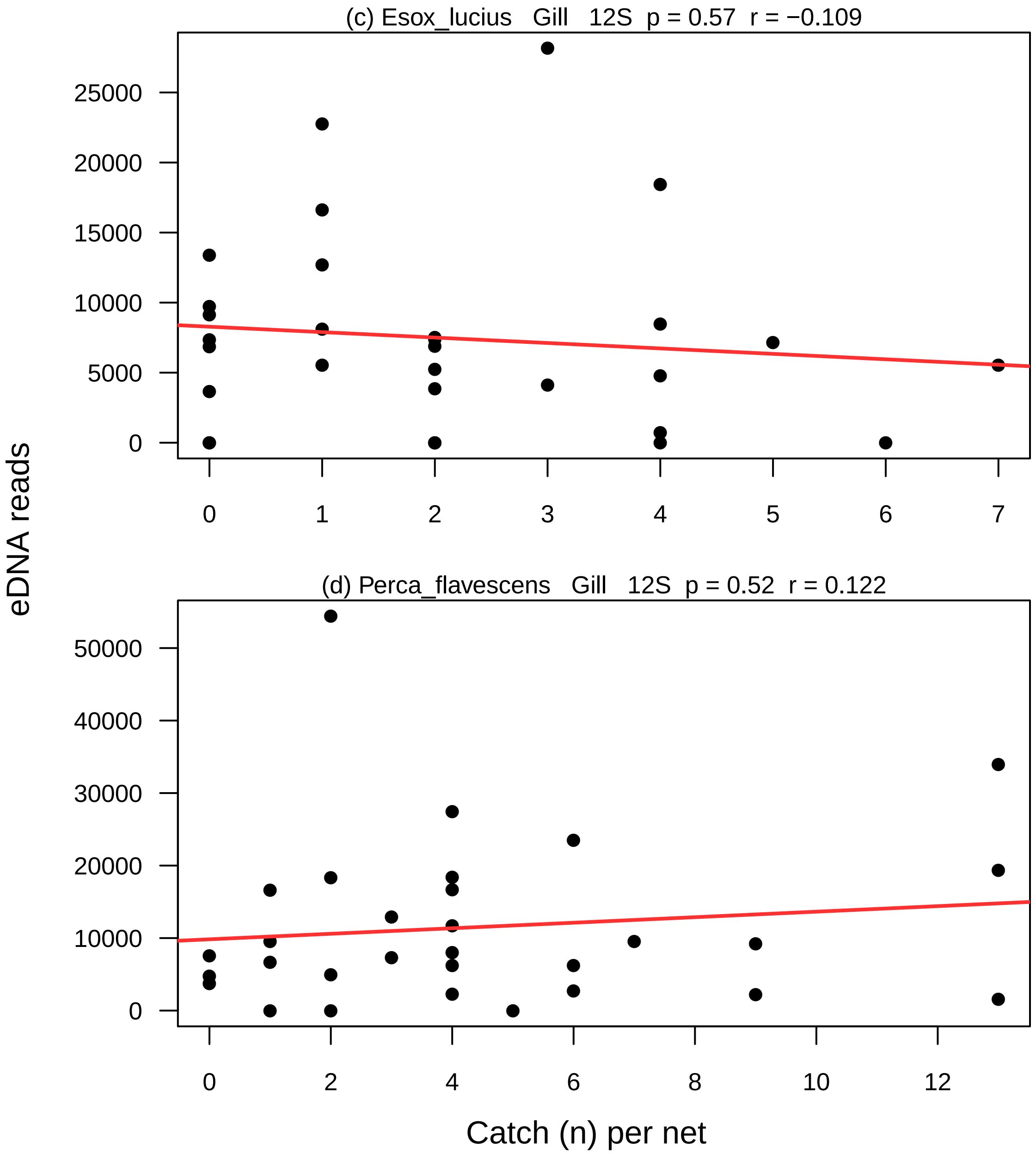

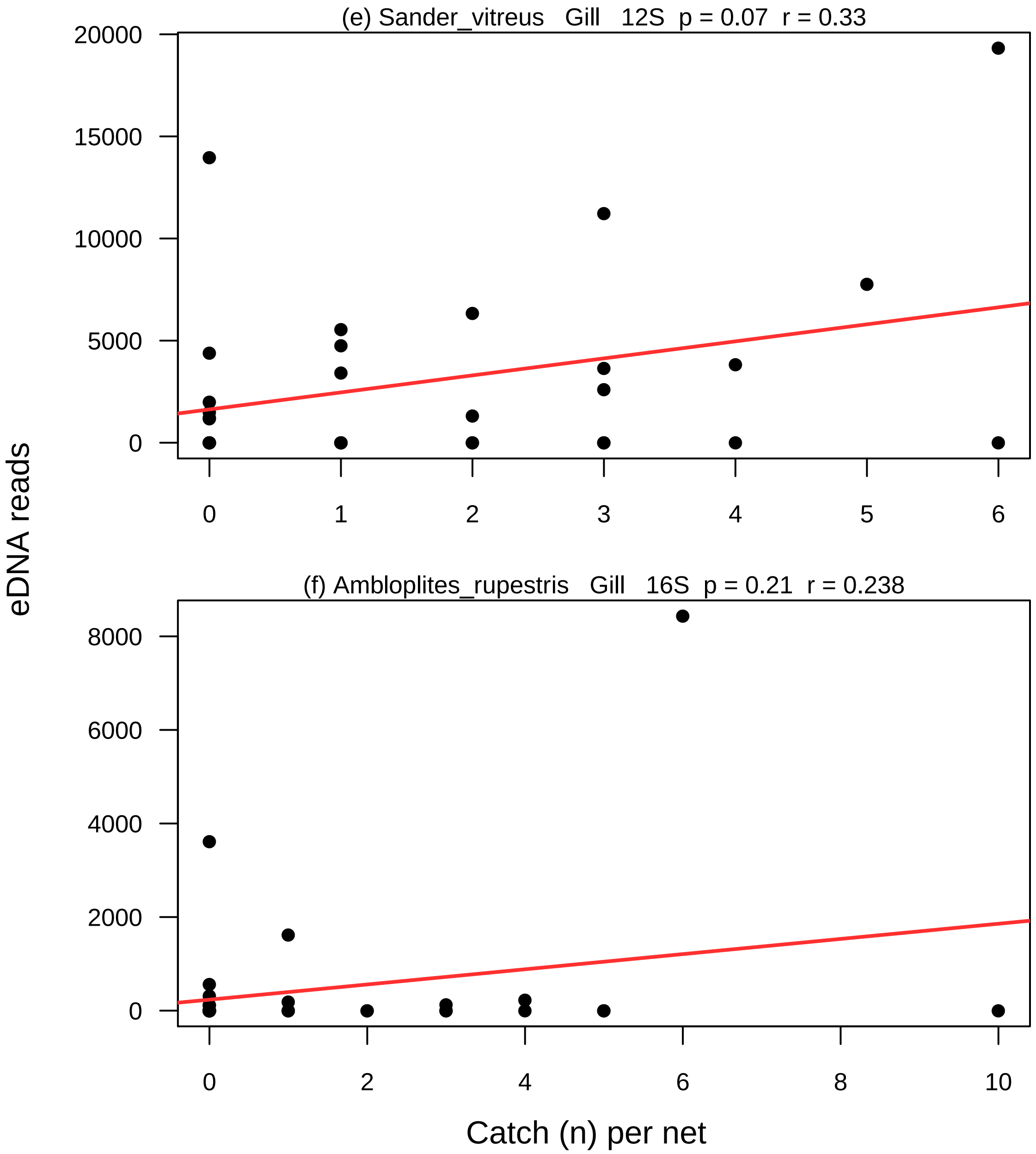

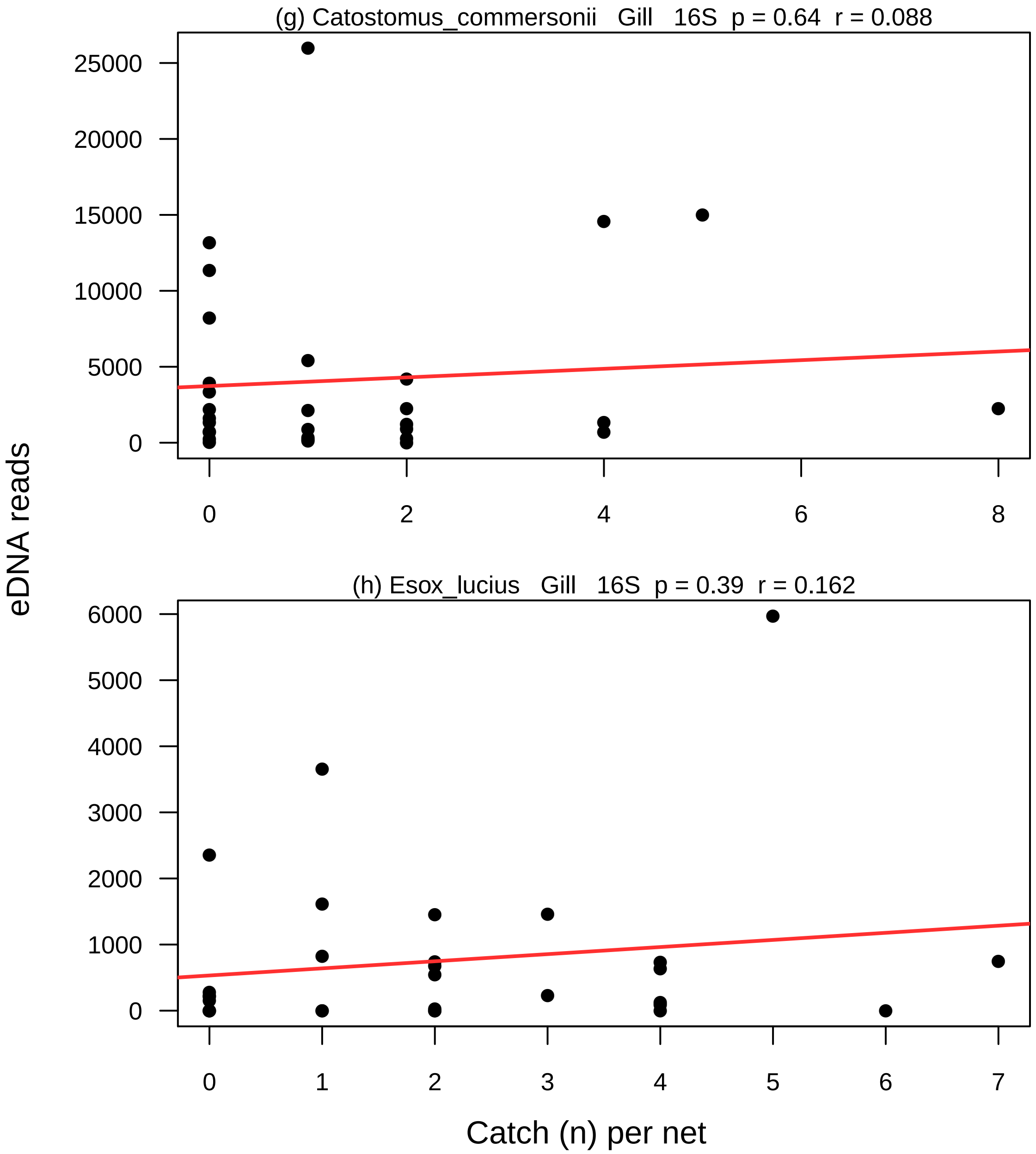

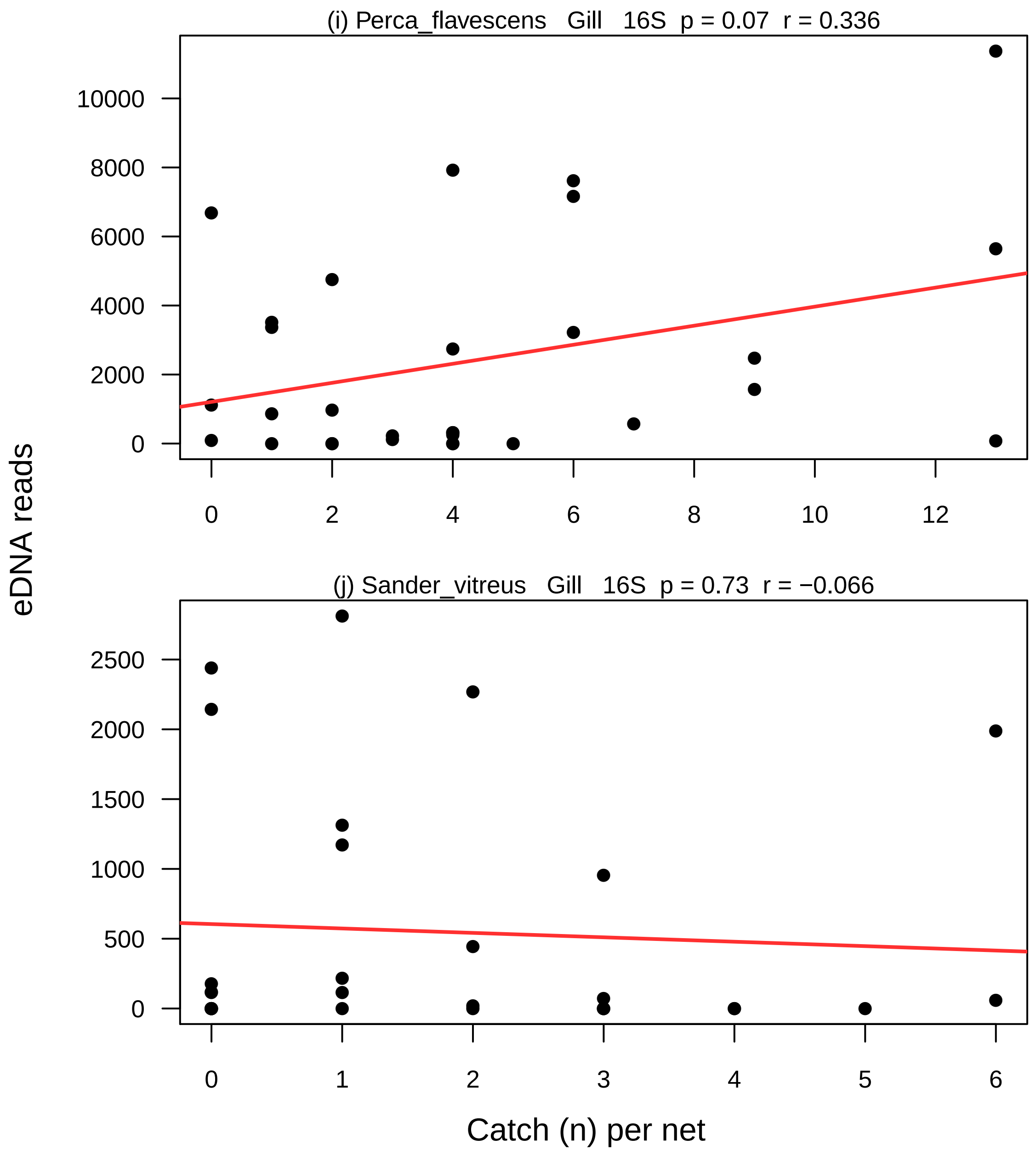

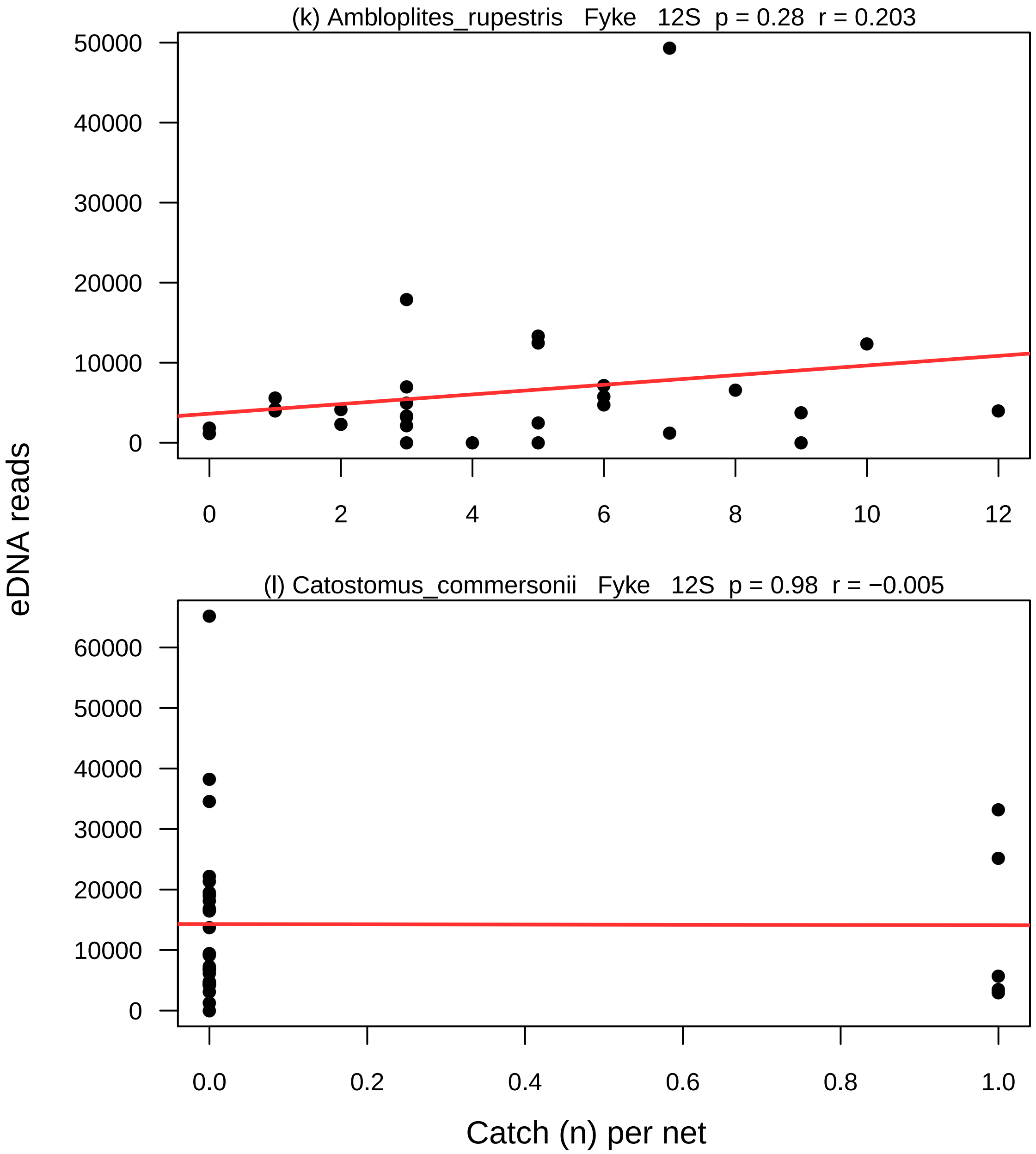

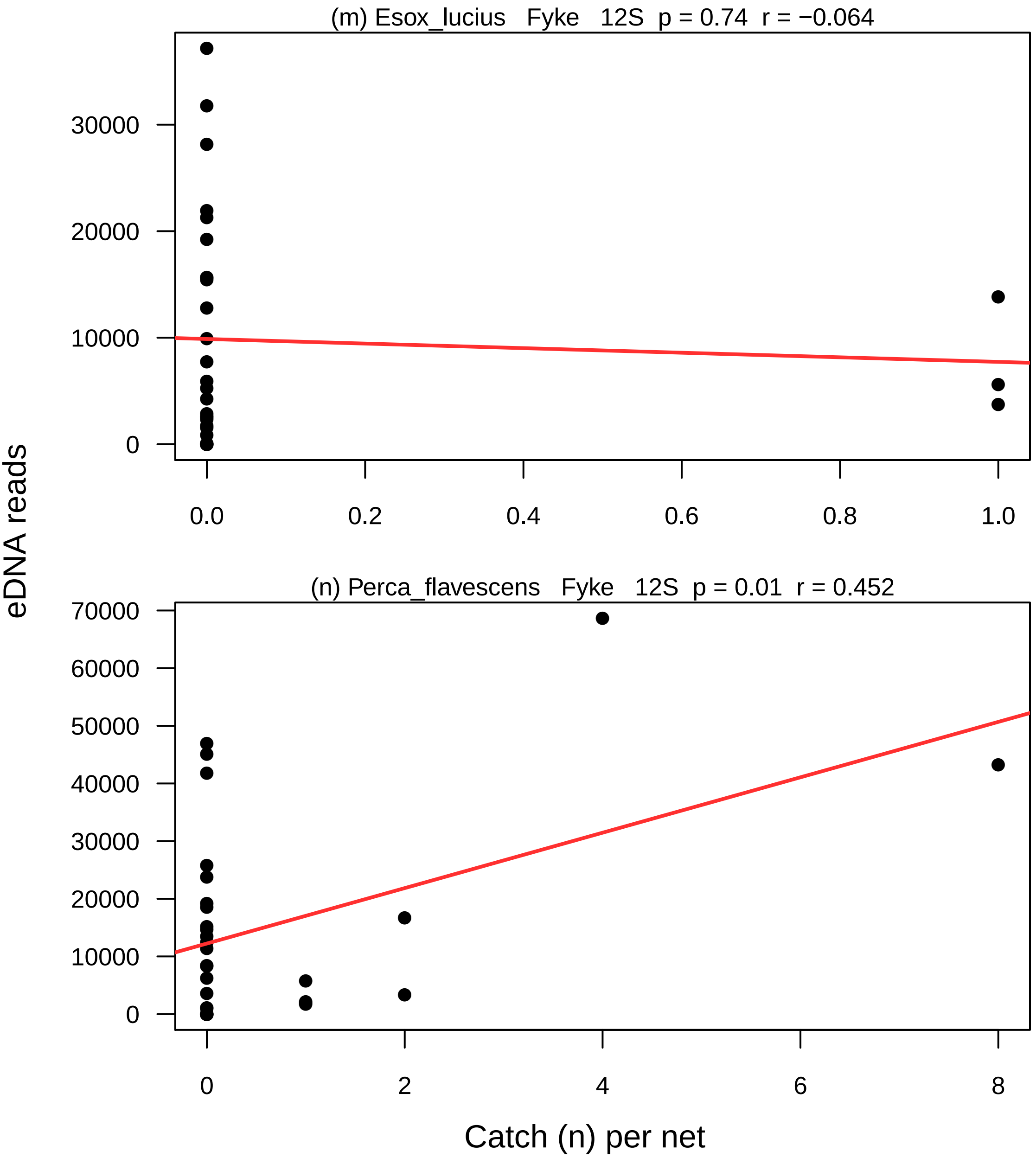

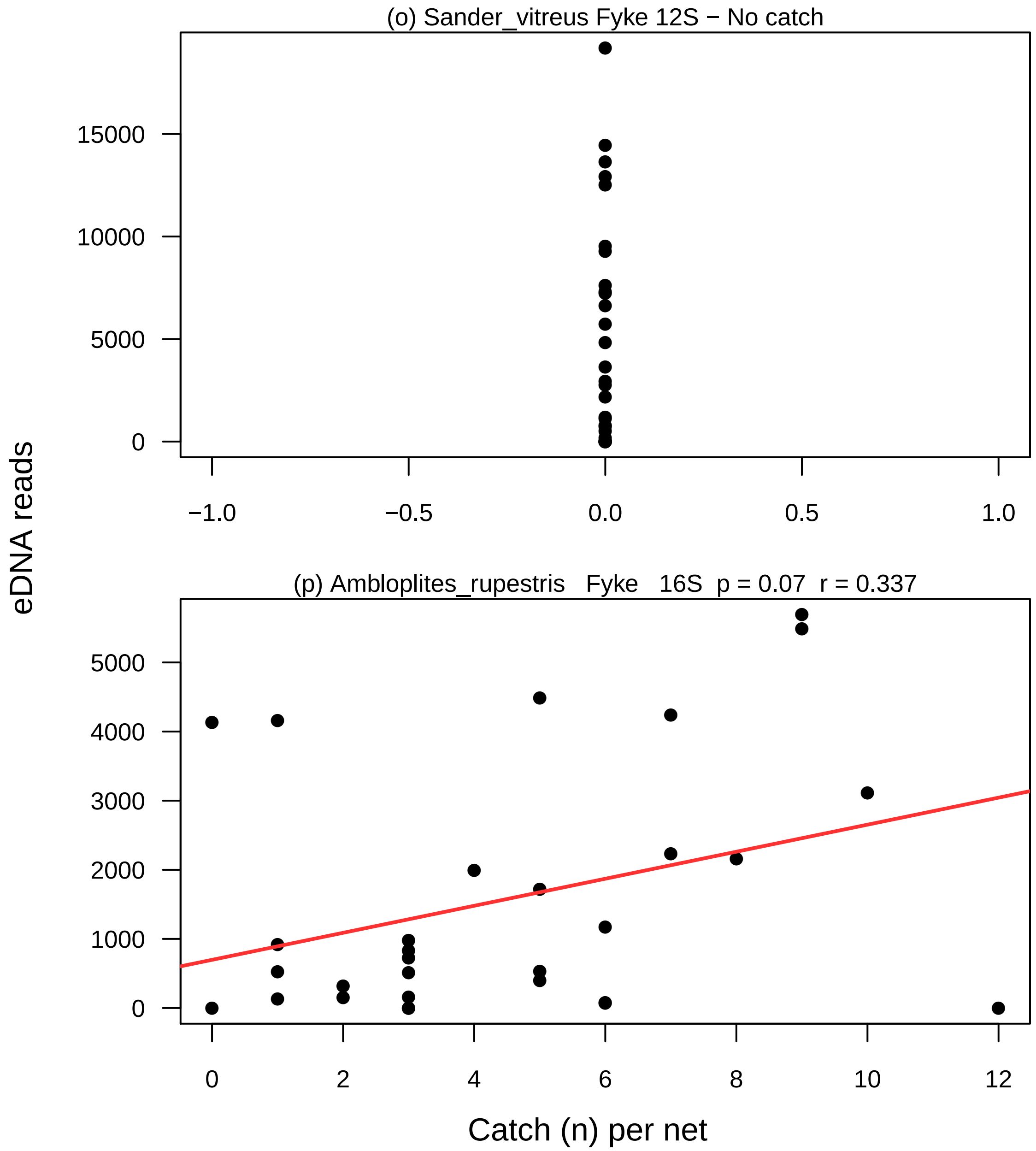

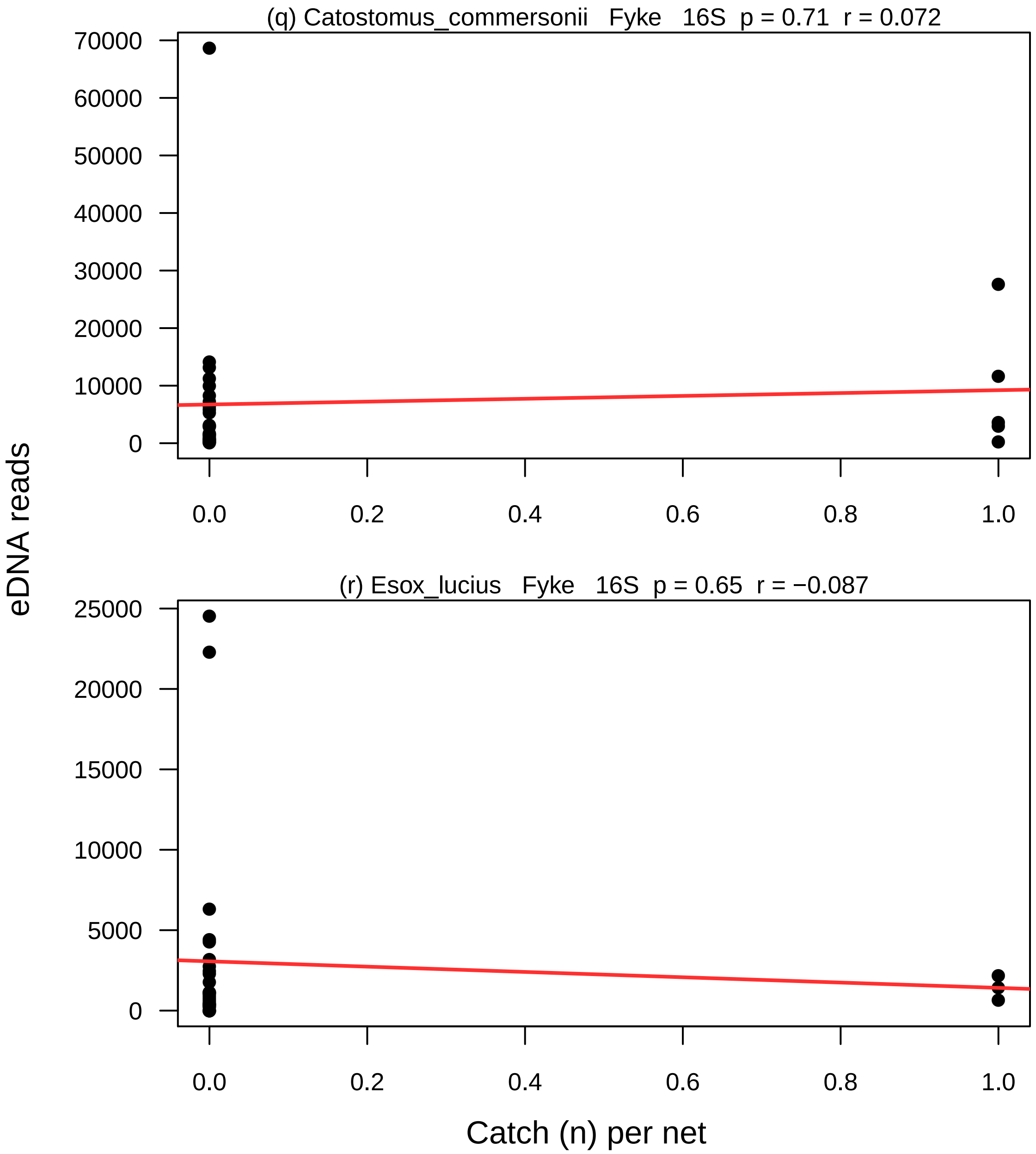

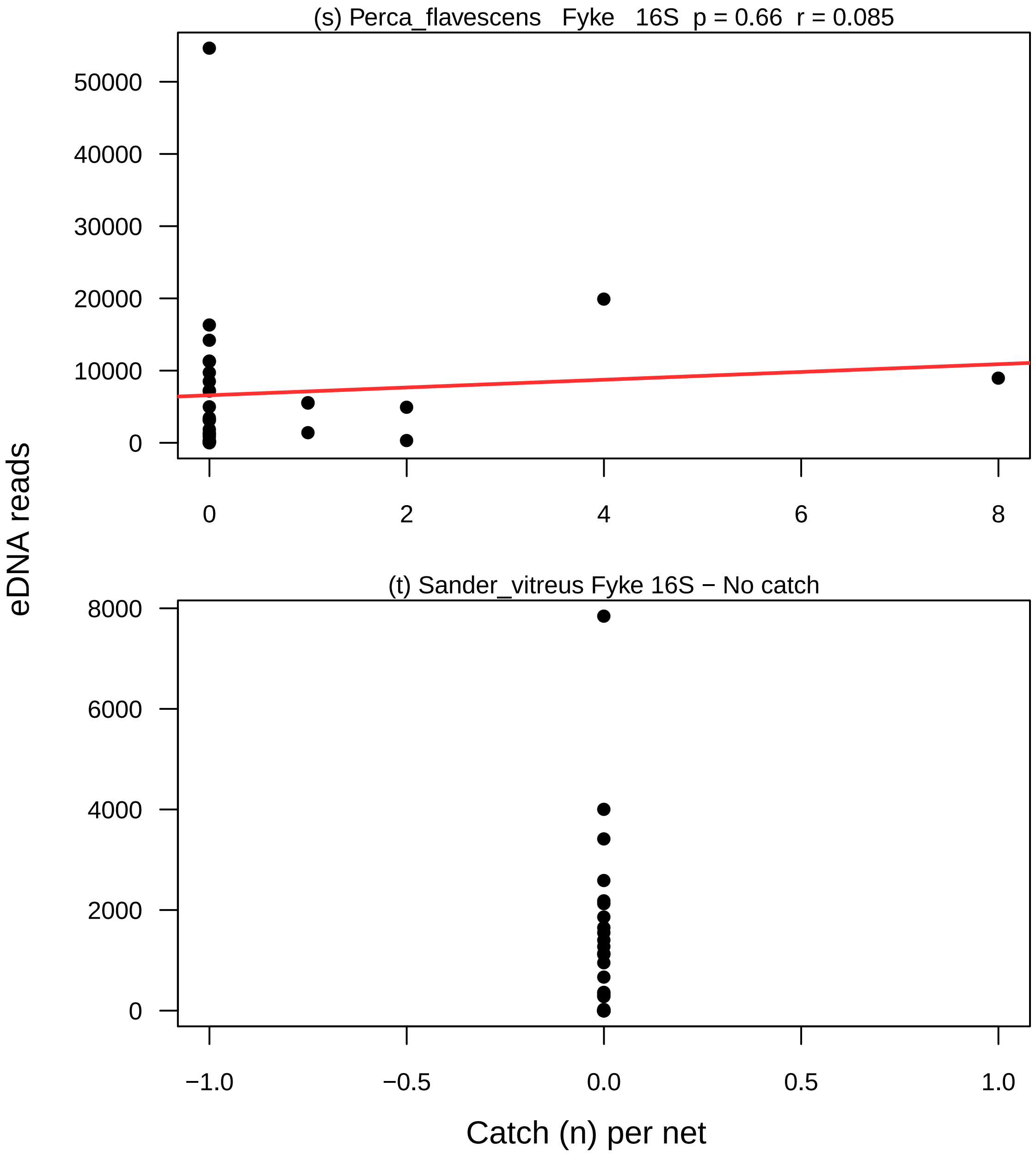

### Figure S11

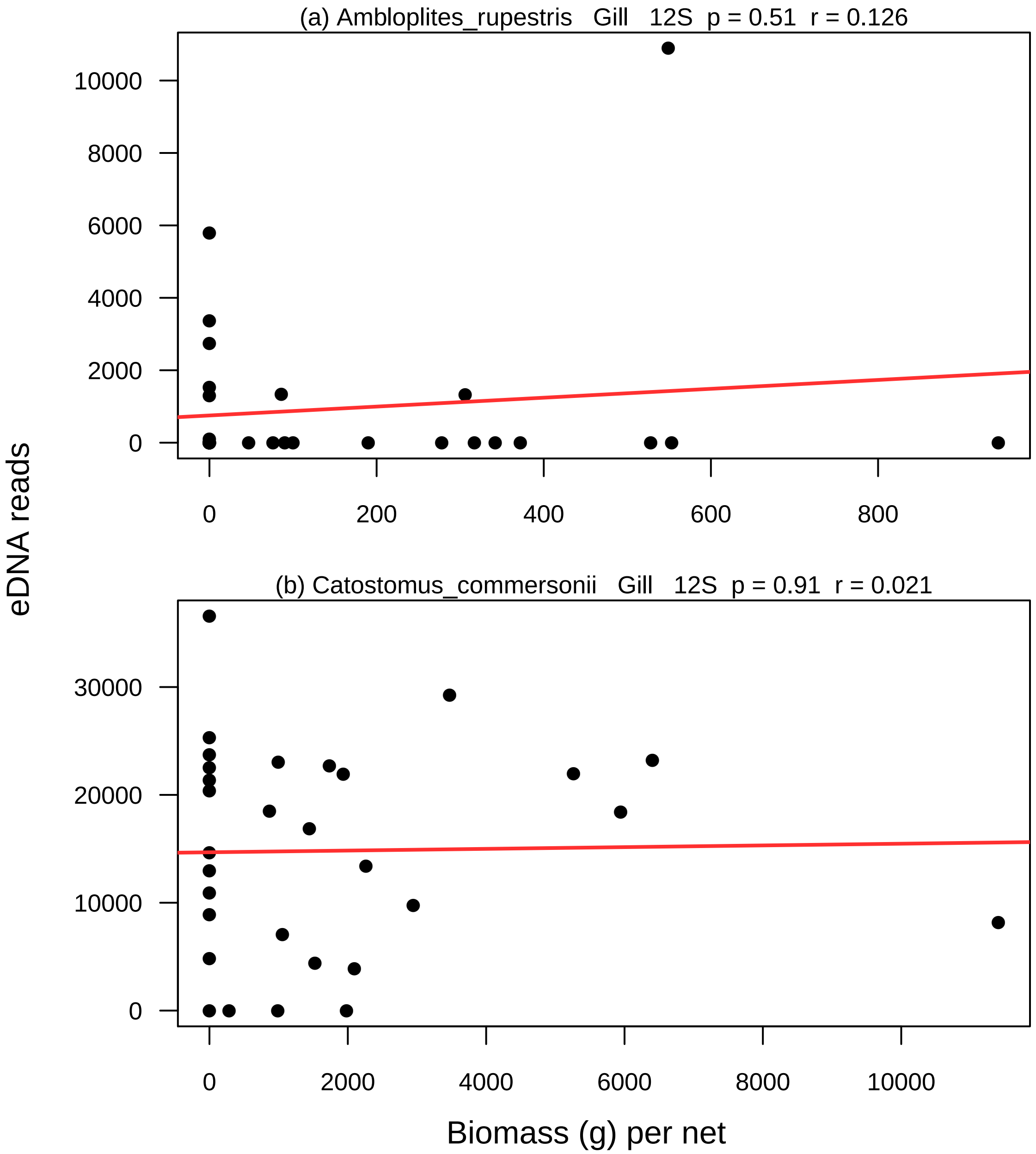

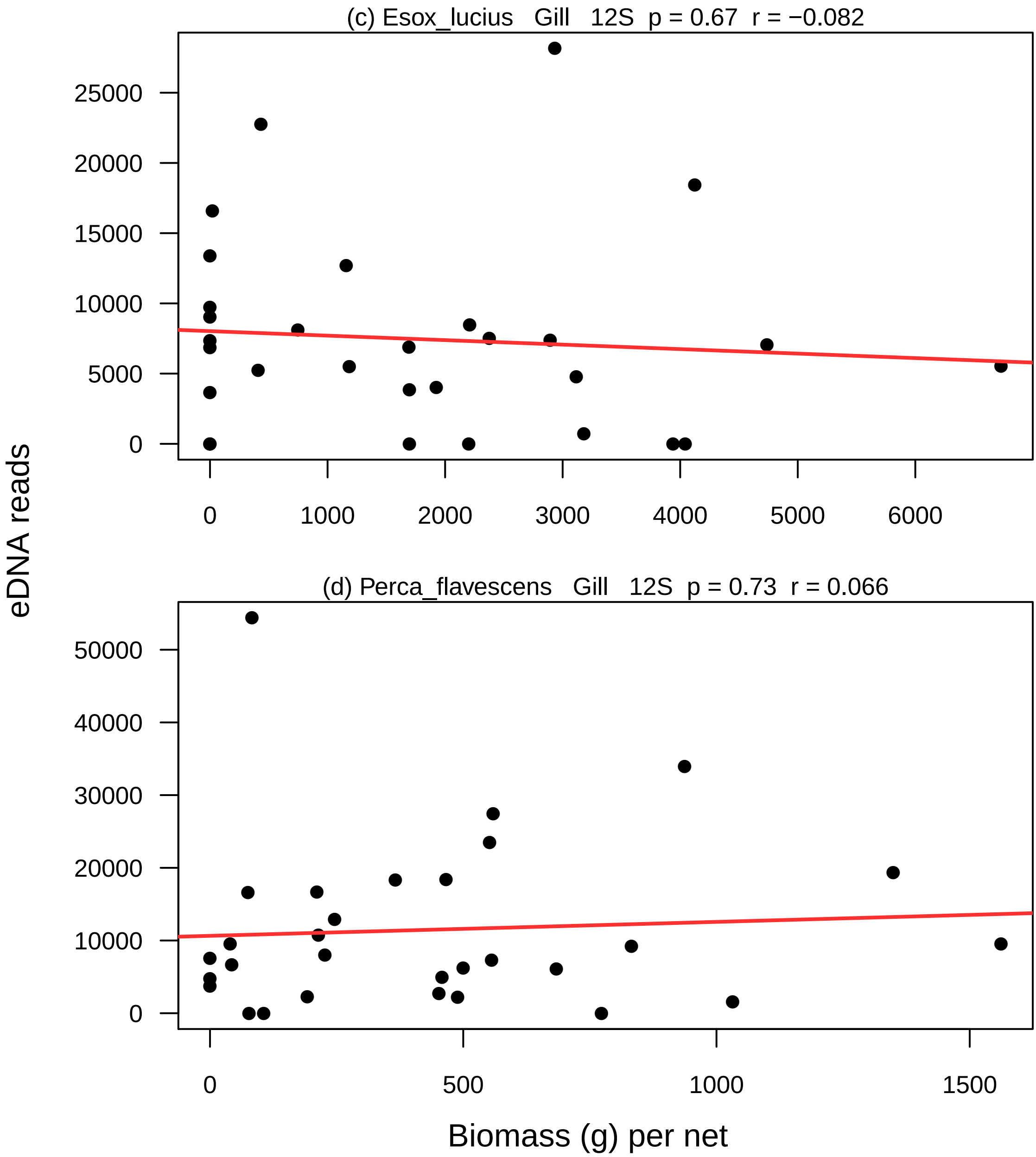

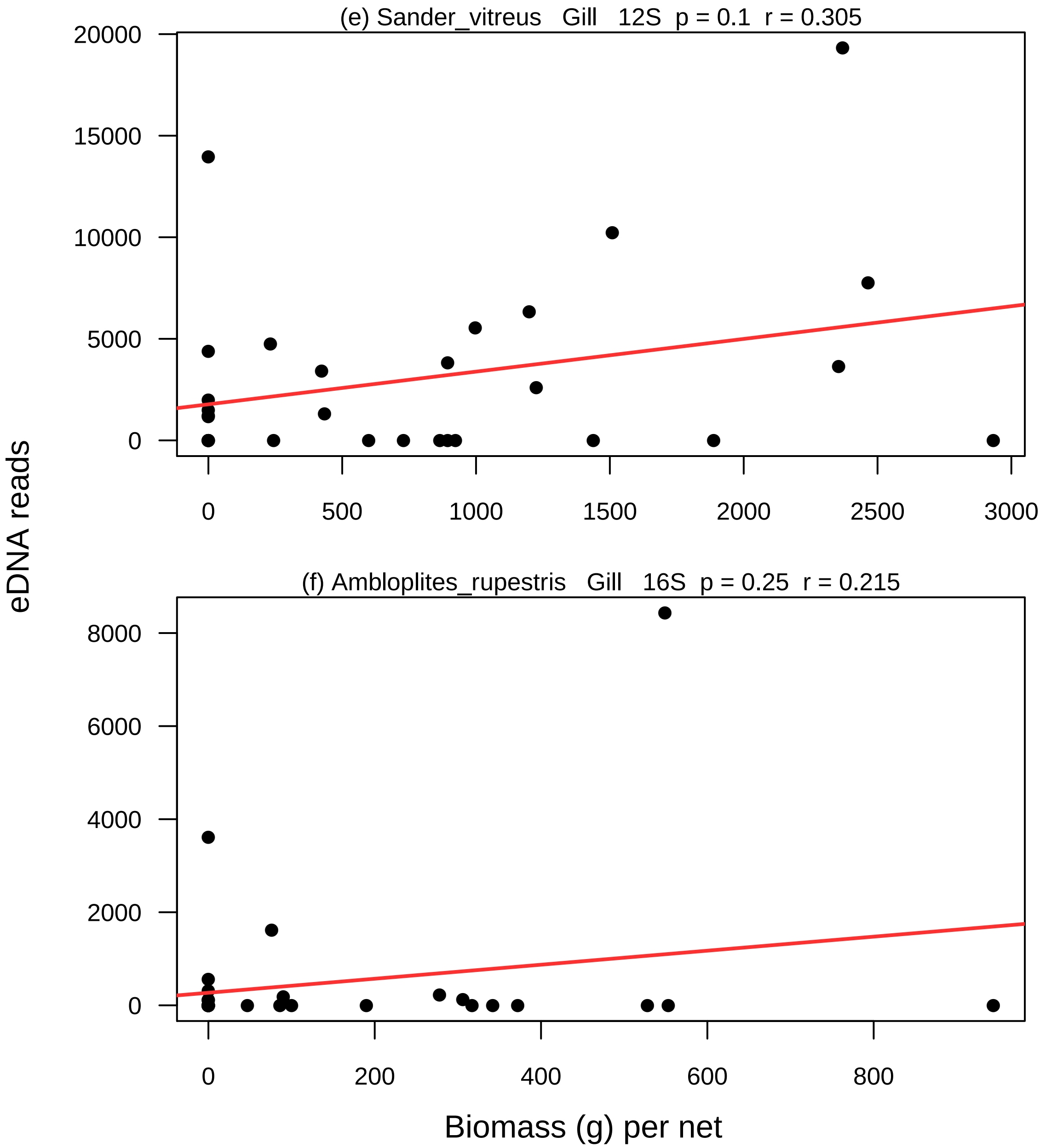

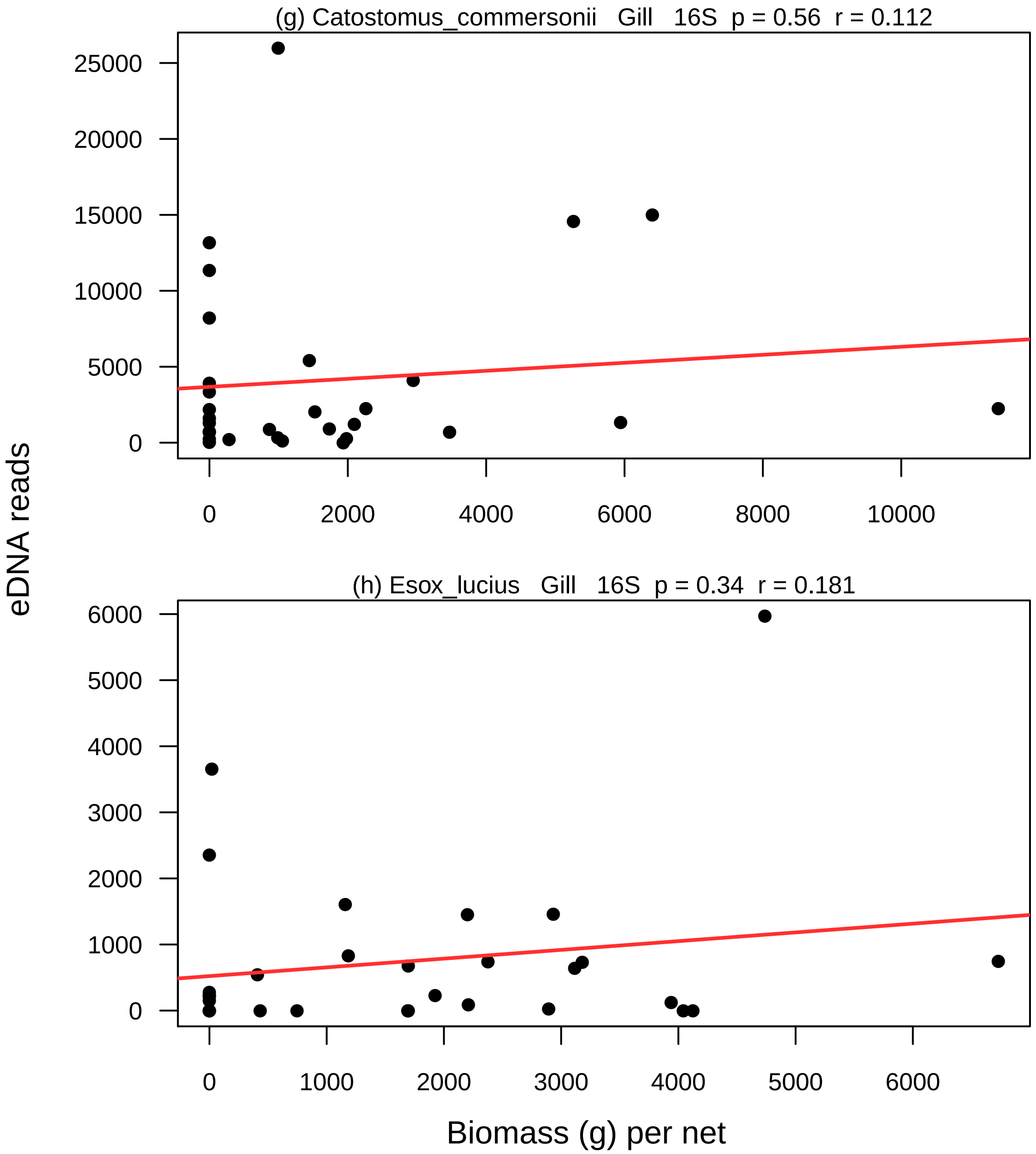

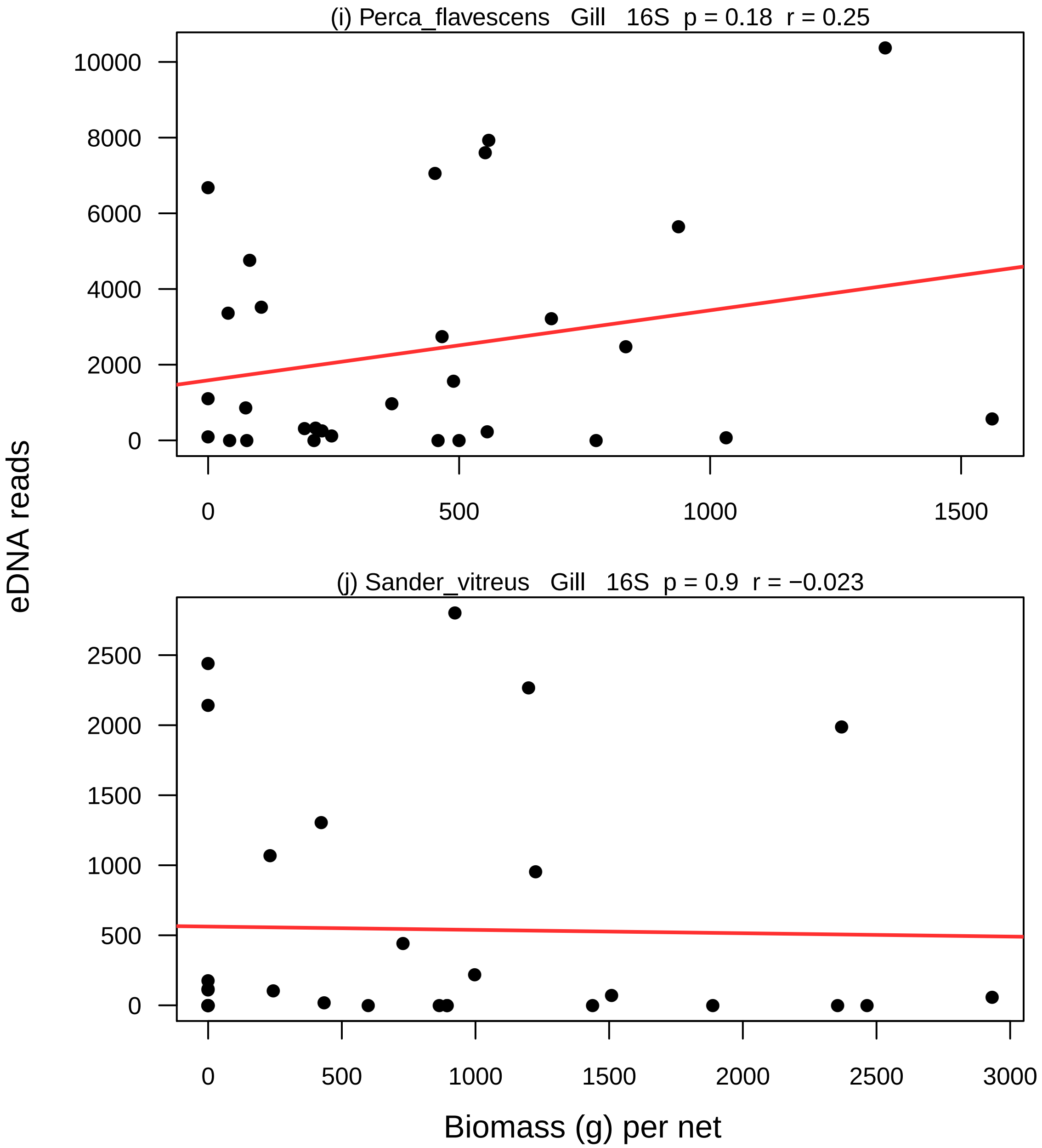

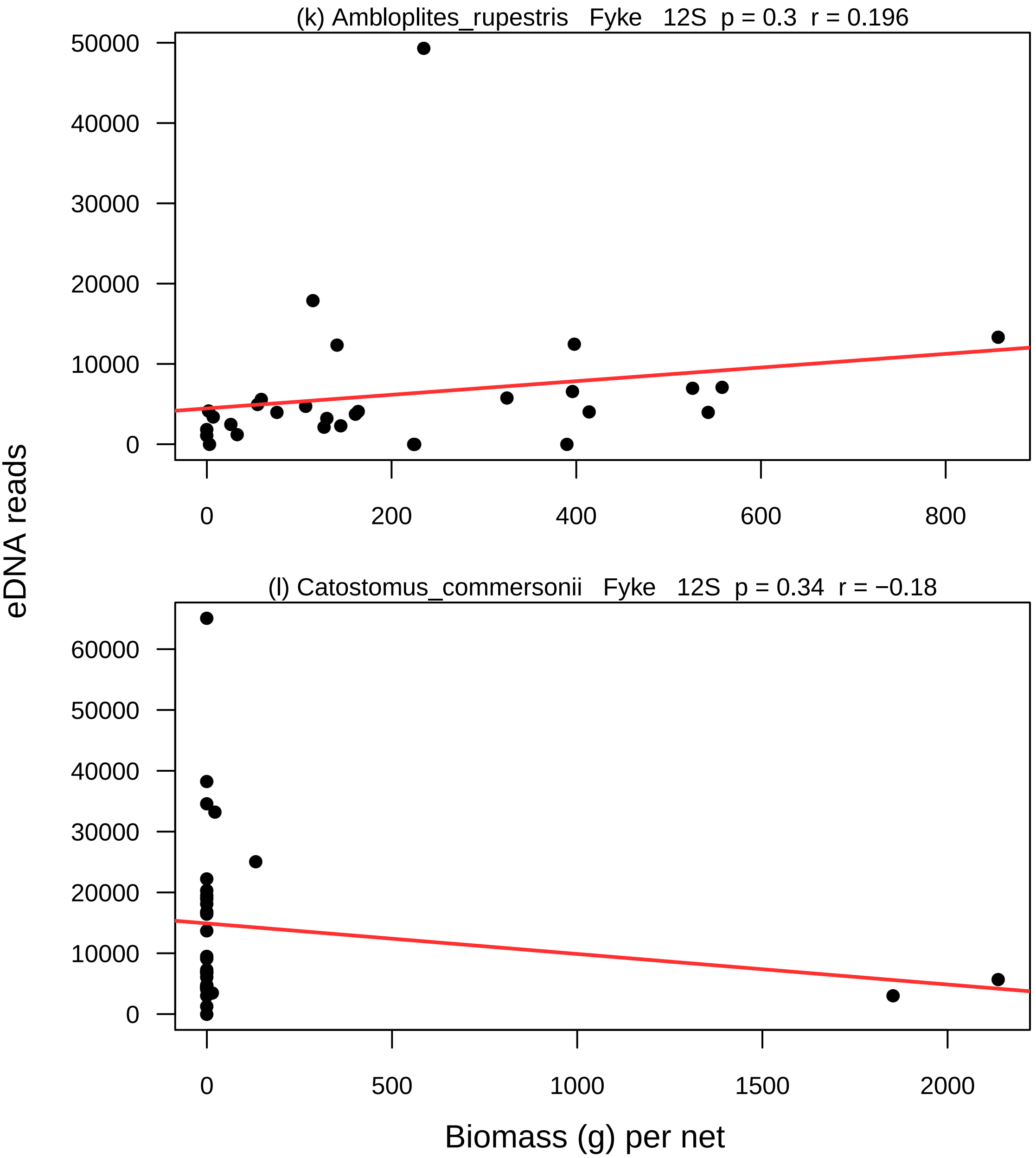

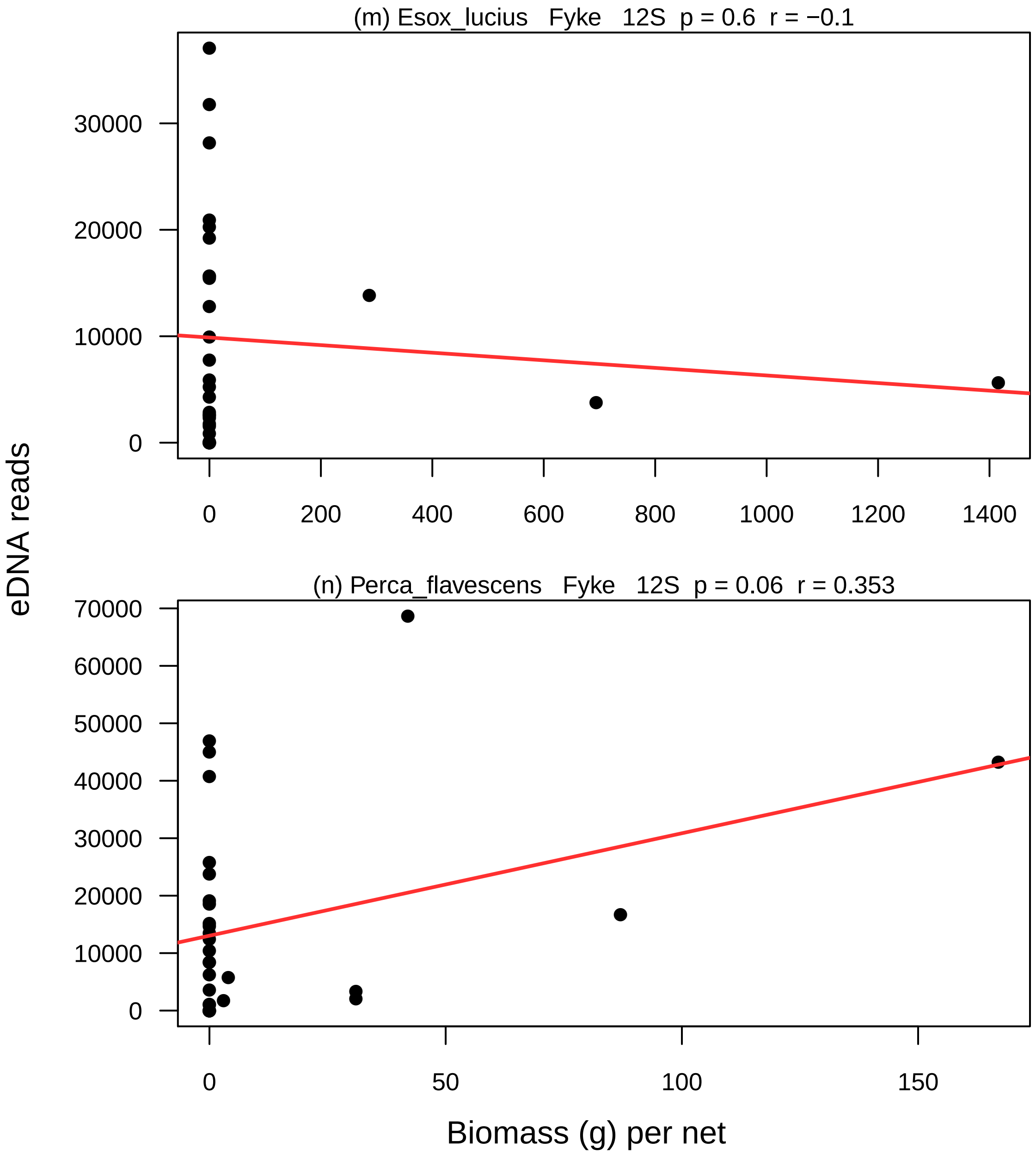

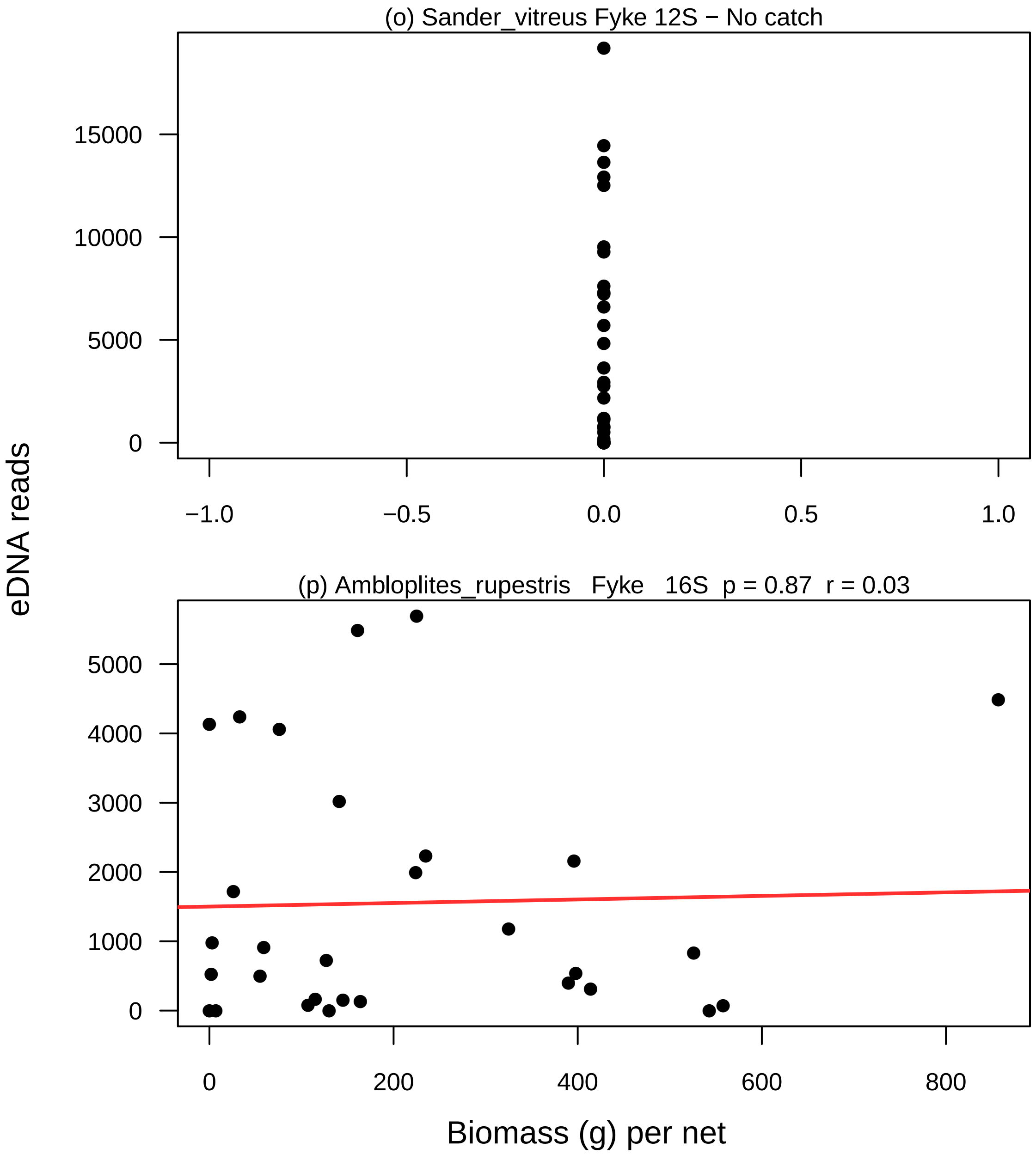

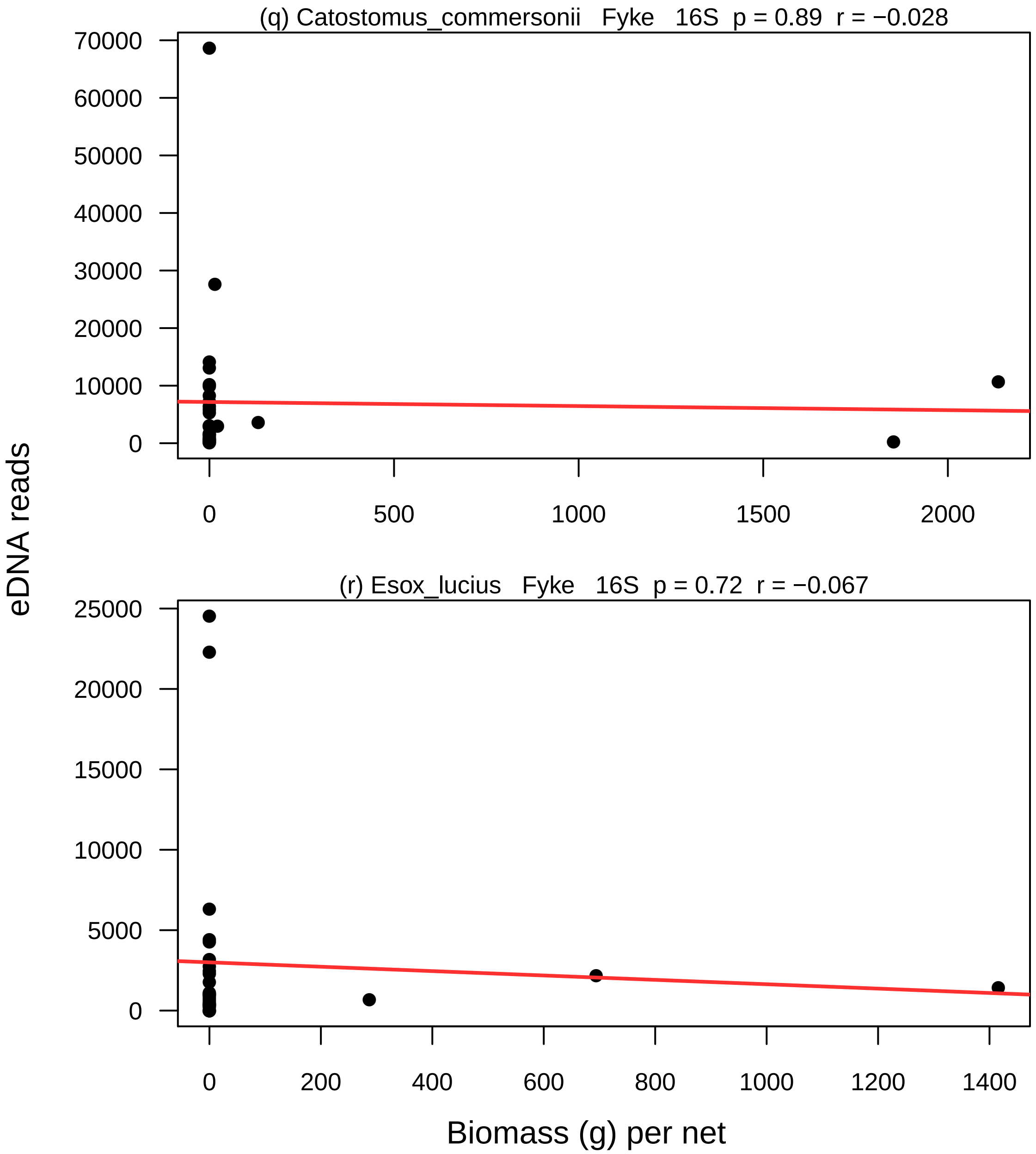

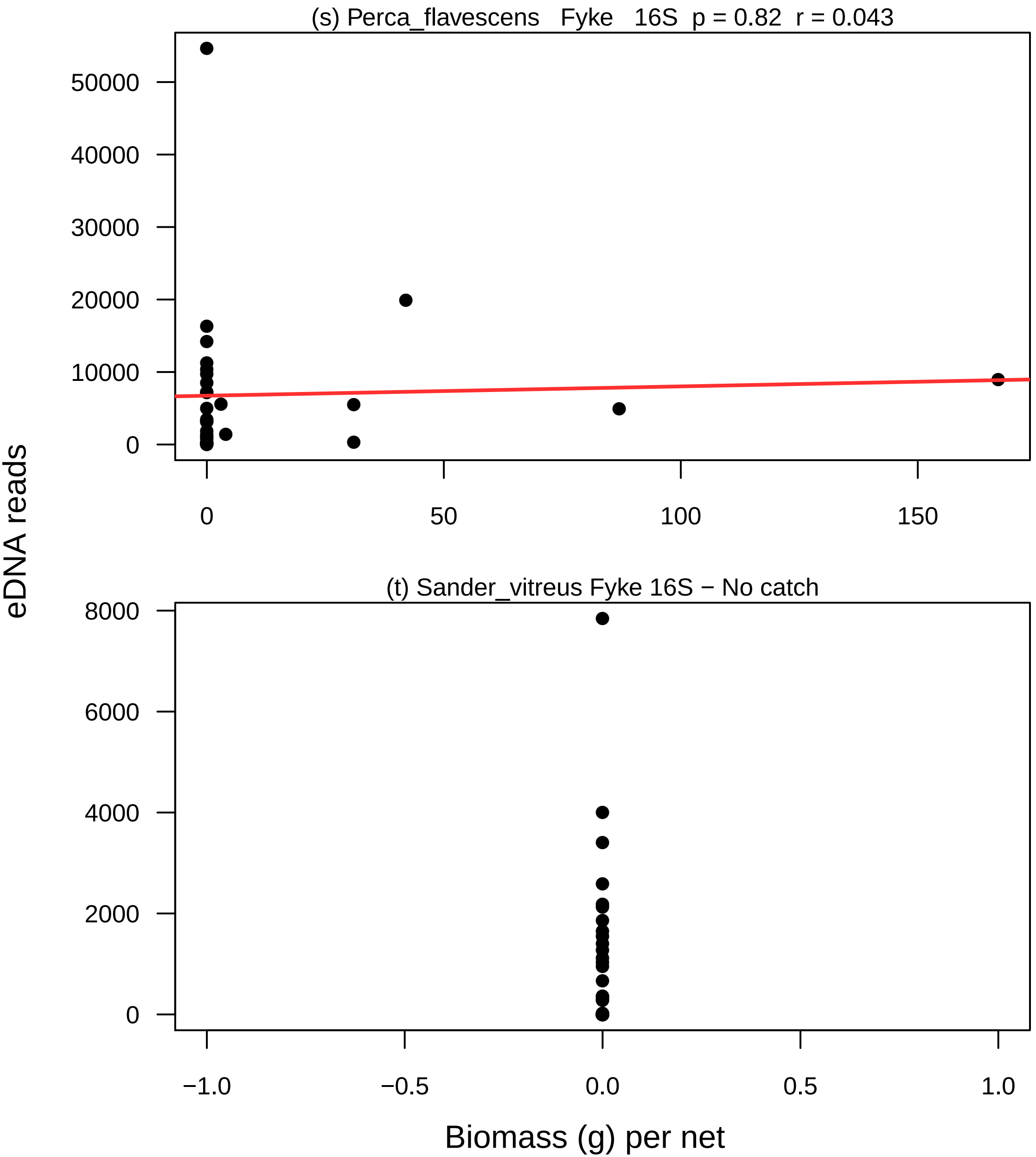

### Figure S12

**Fyke nets vs gill nets**

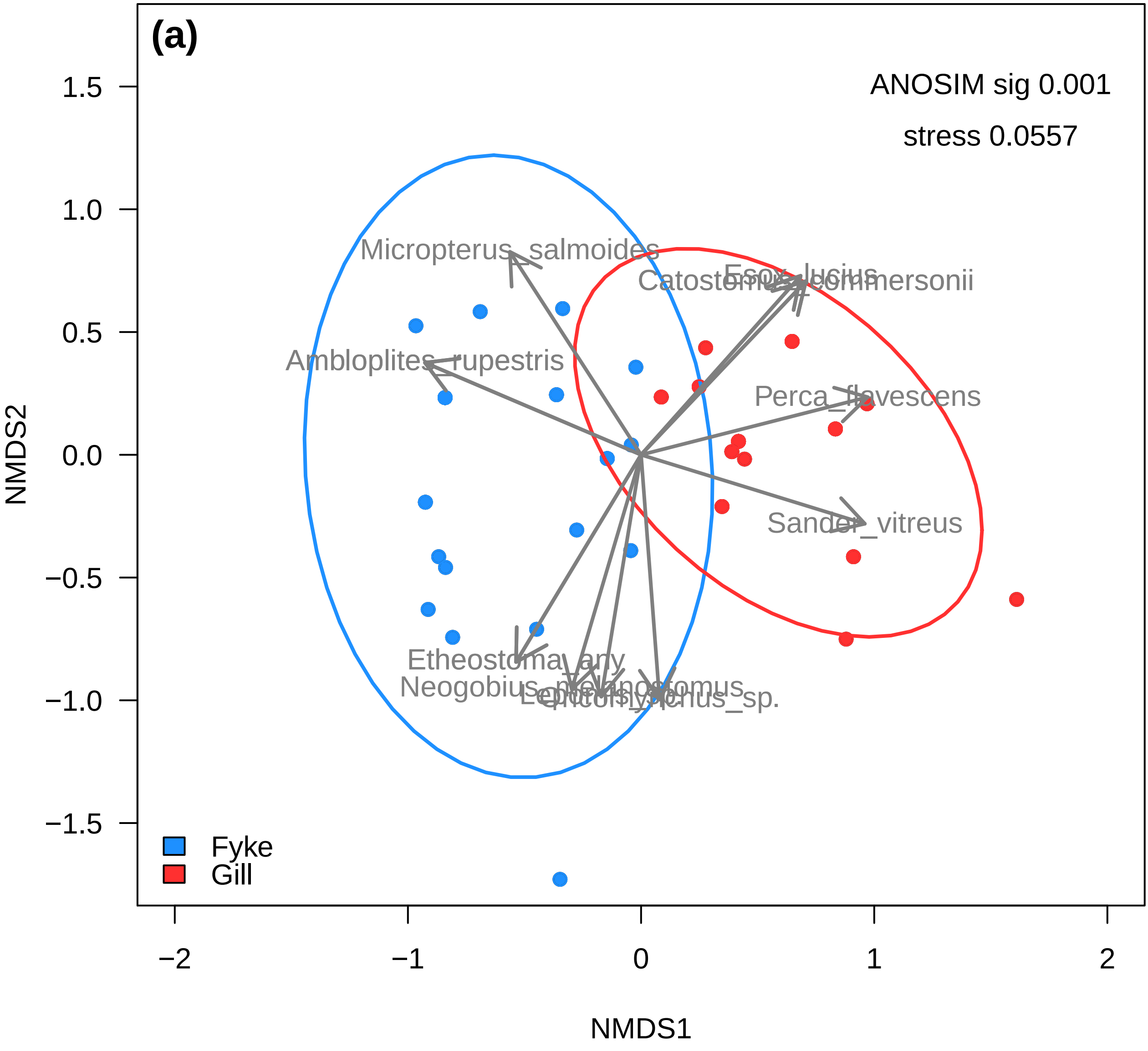

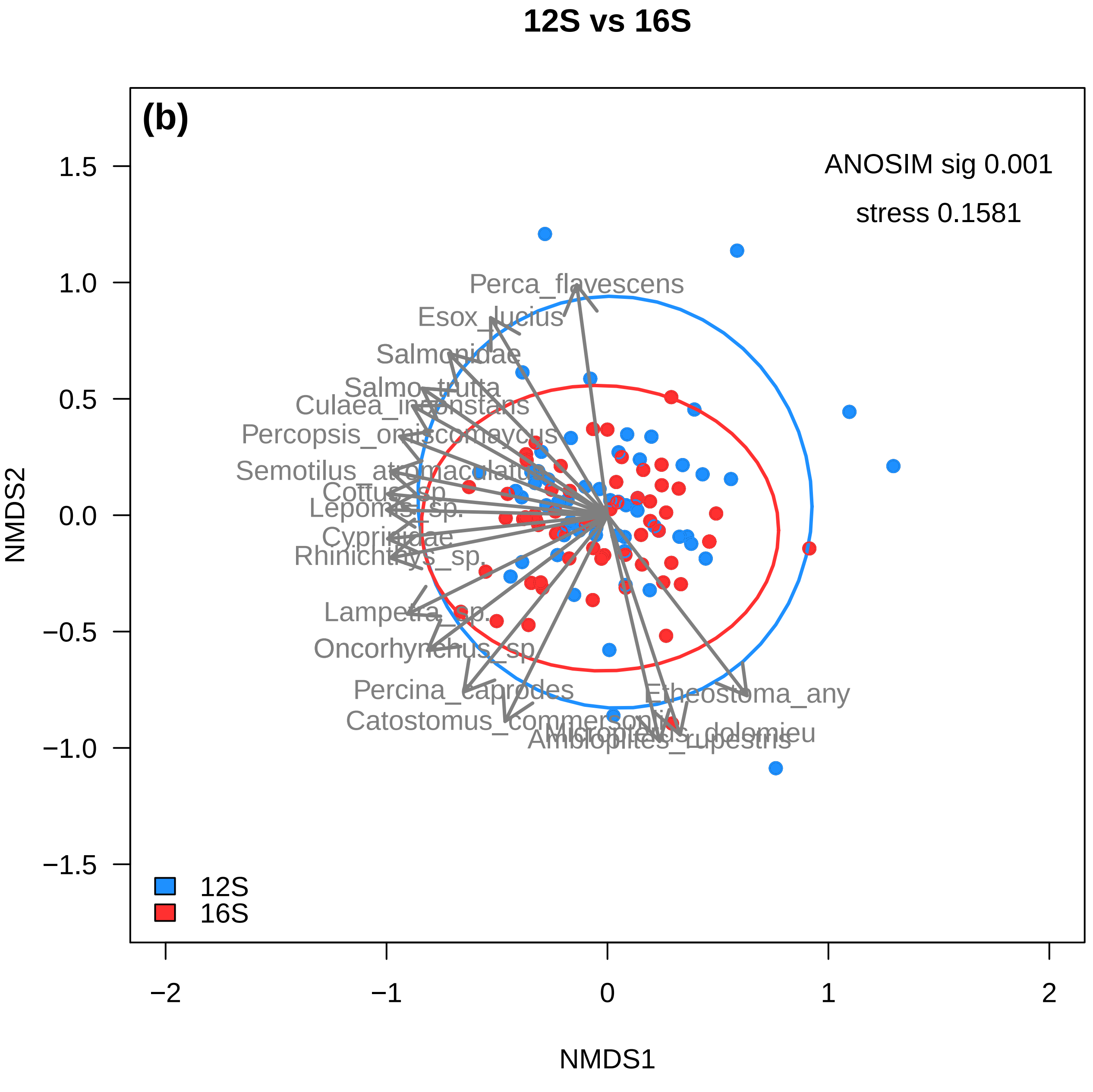

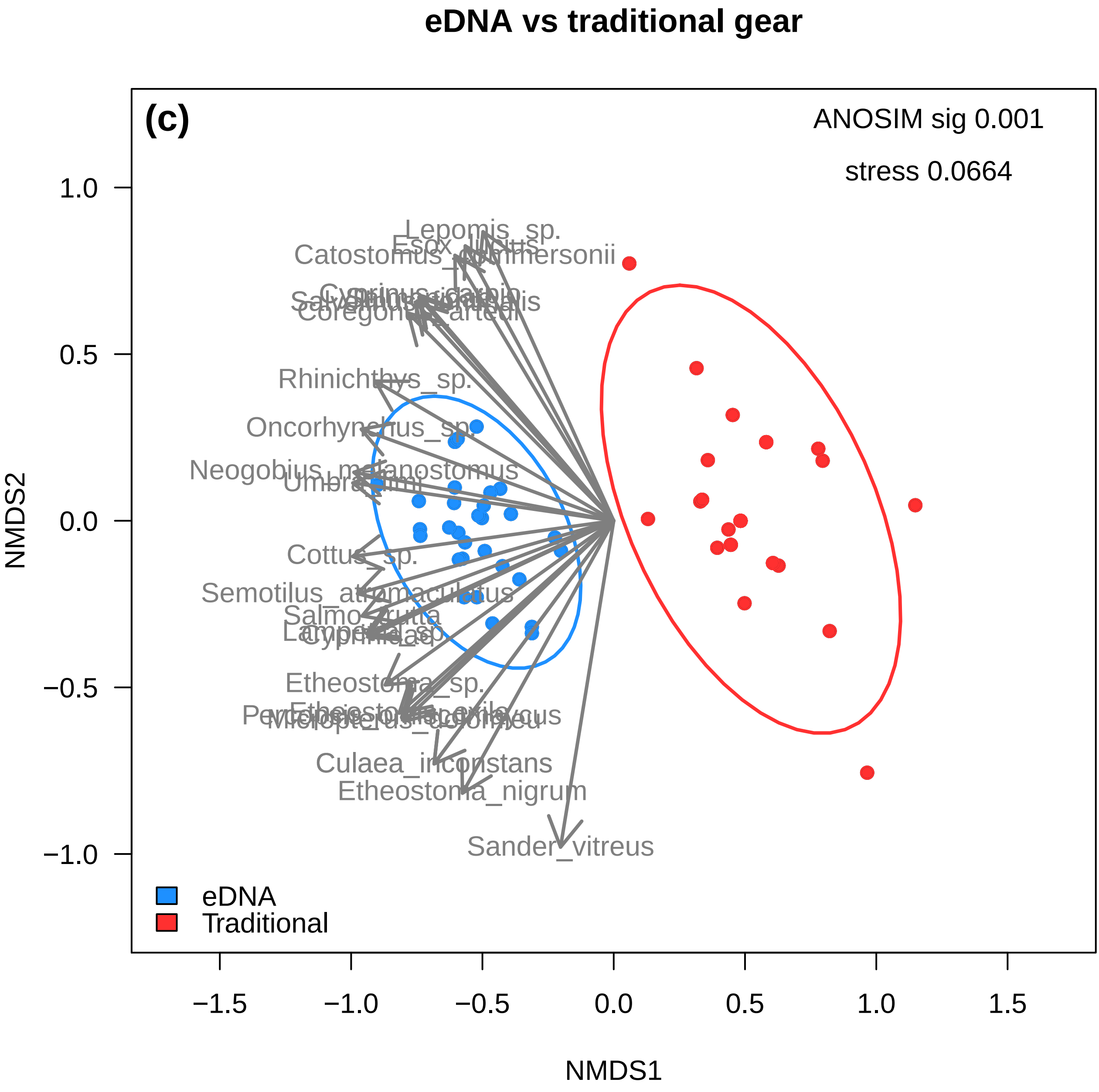

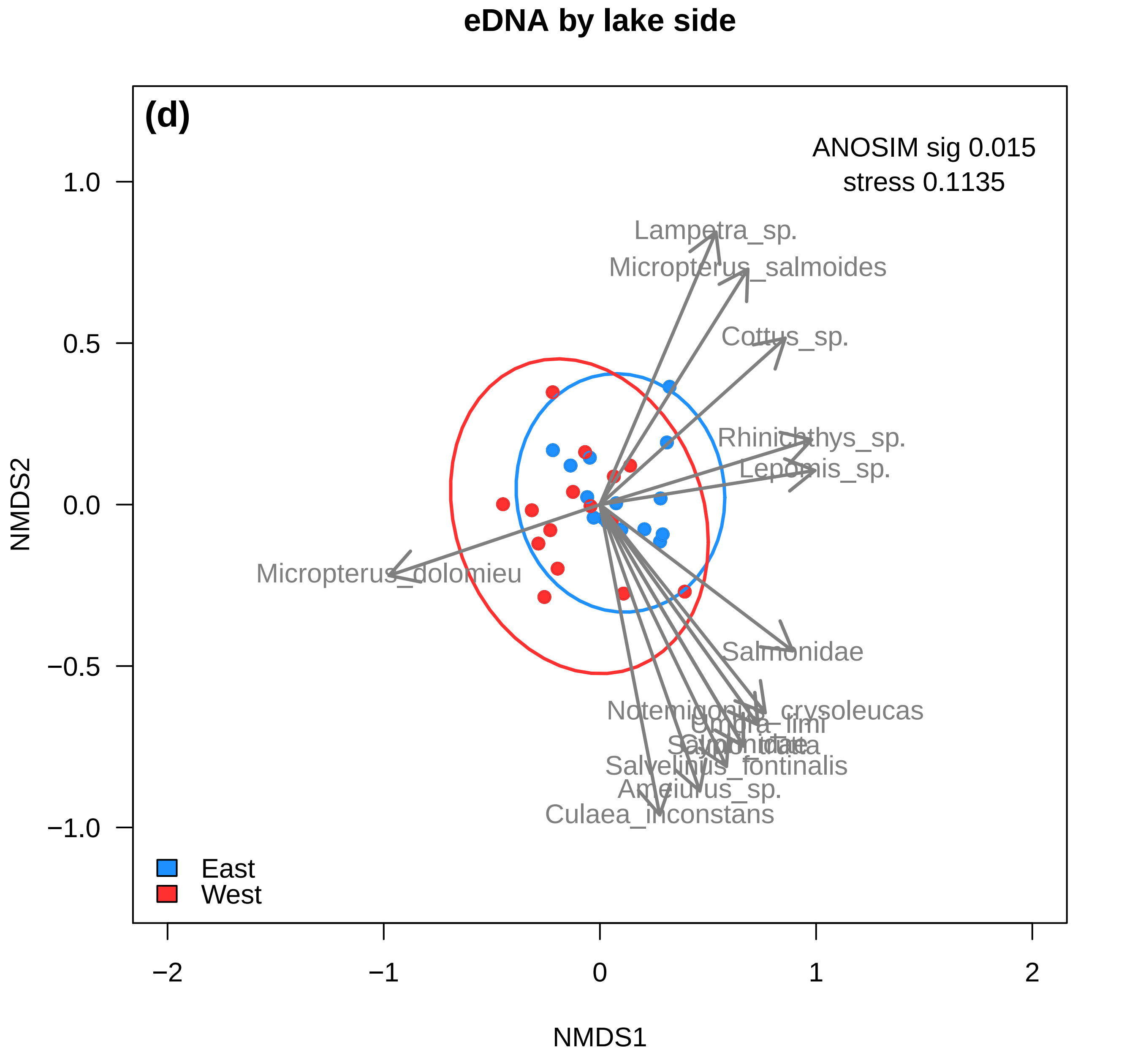

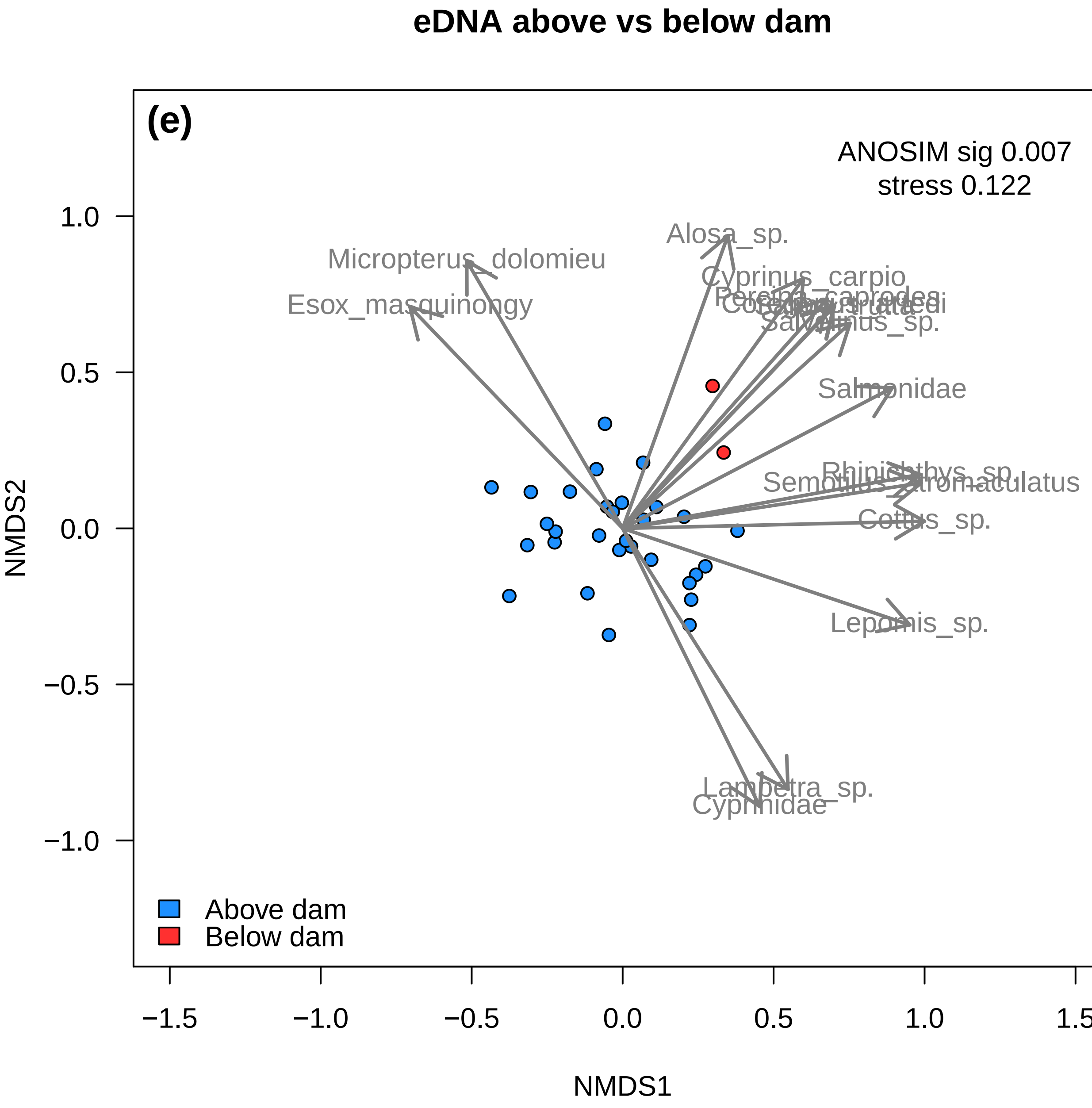

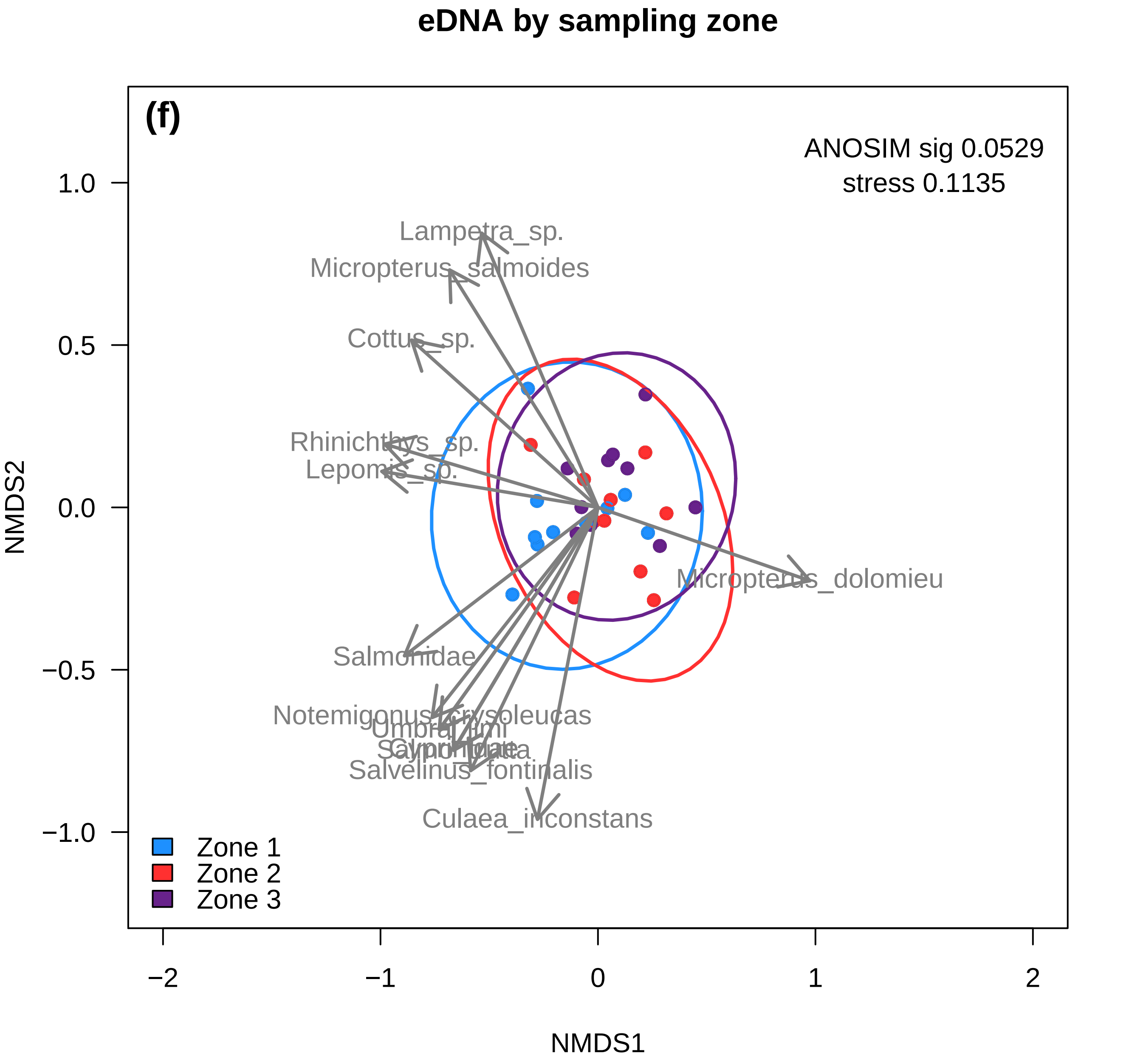

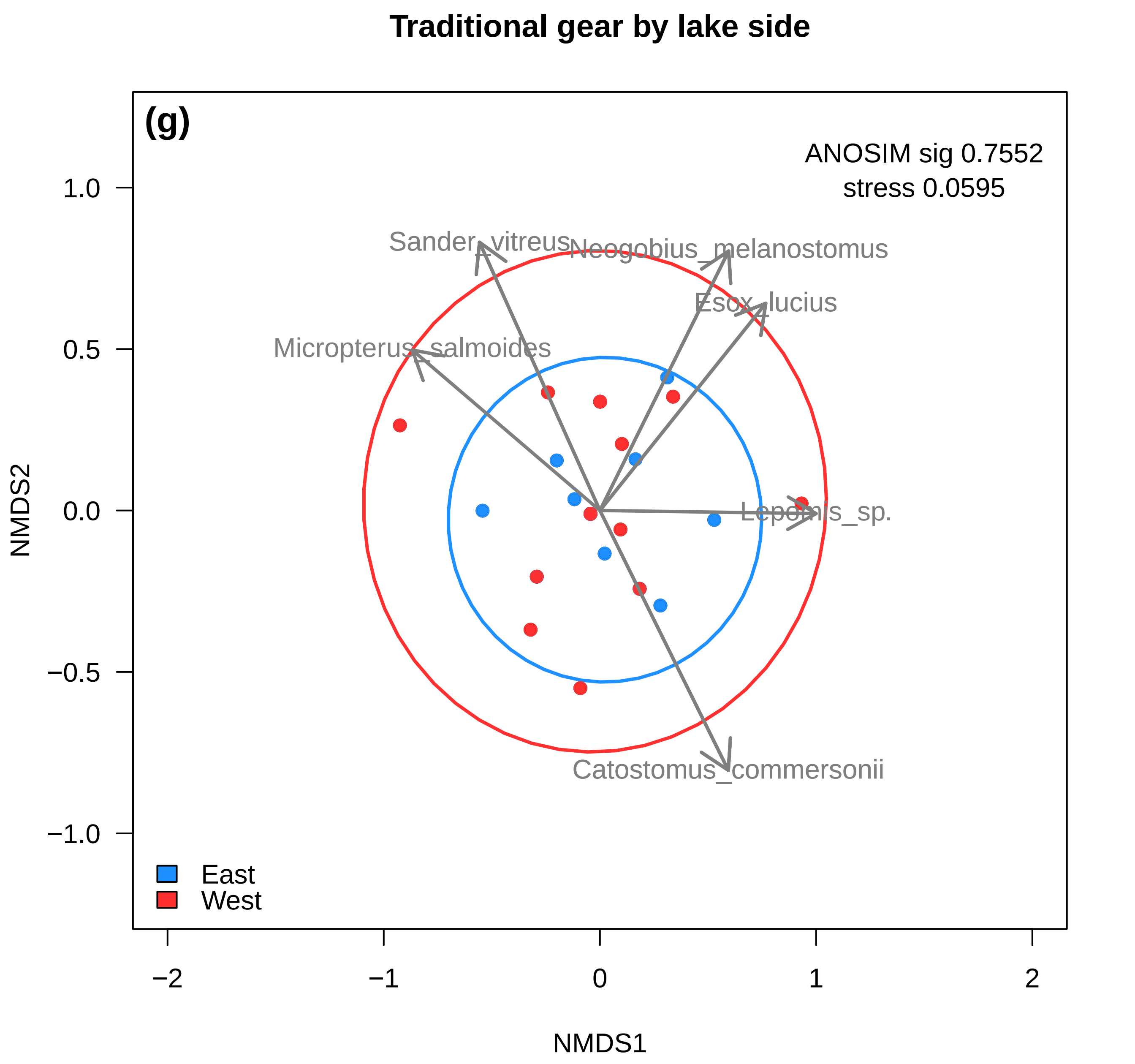

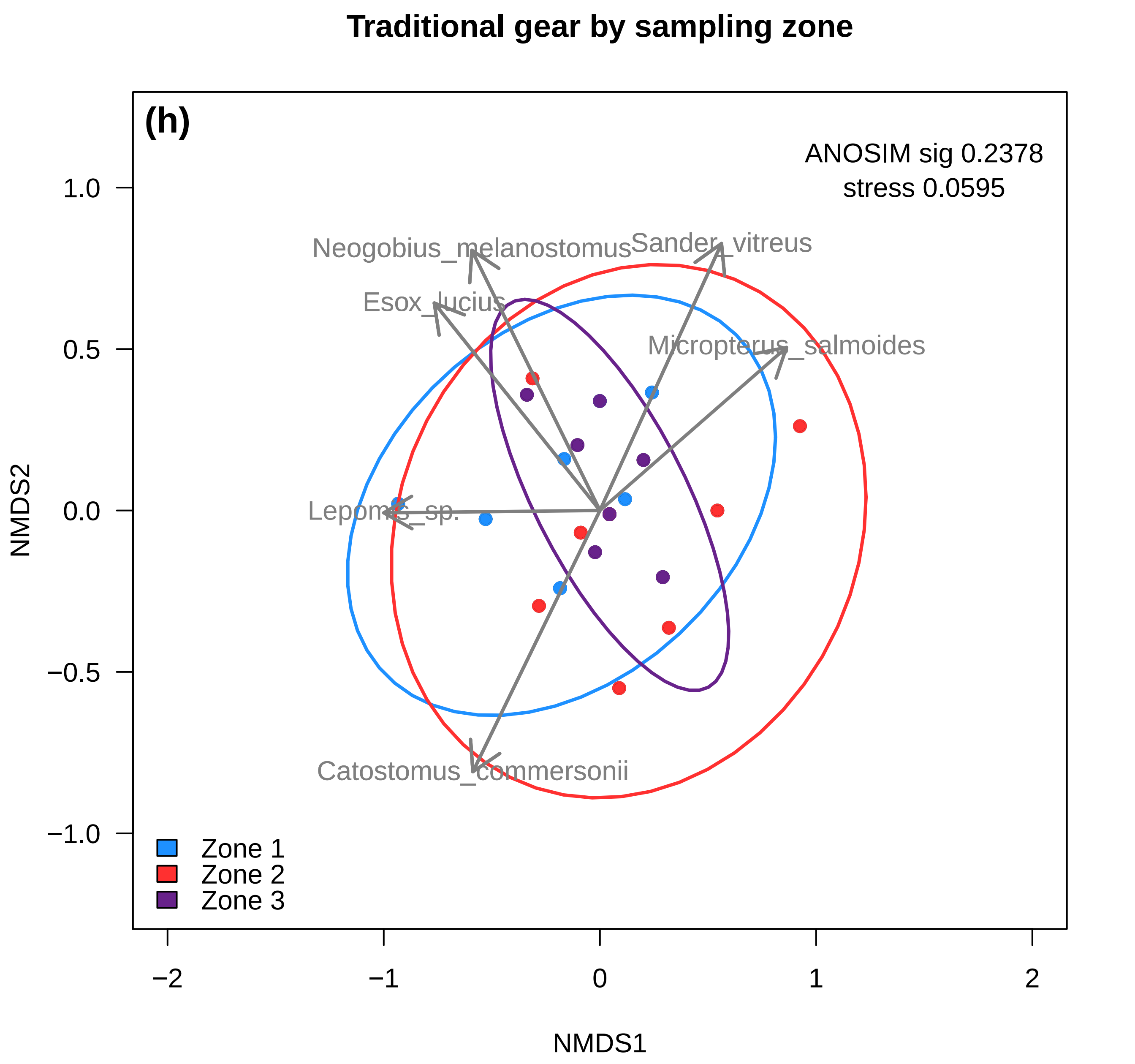

### Figure S13

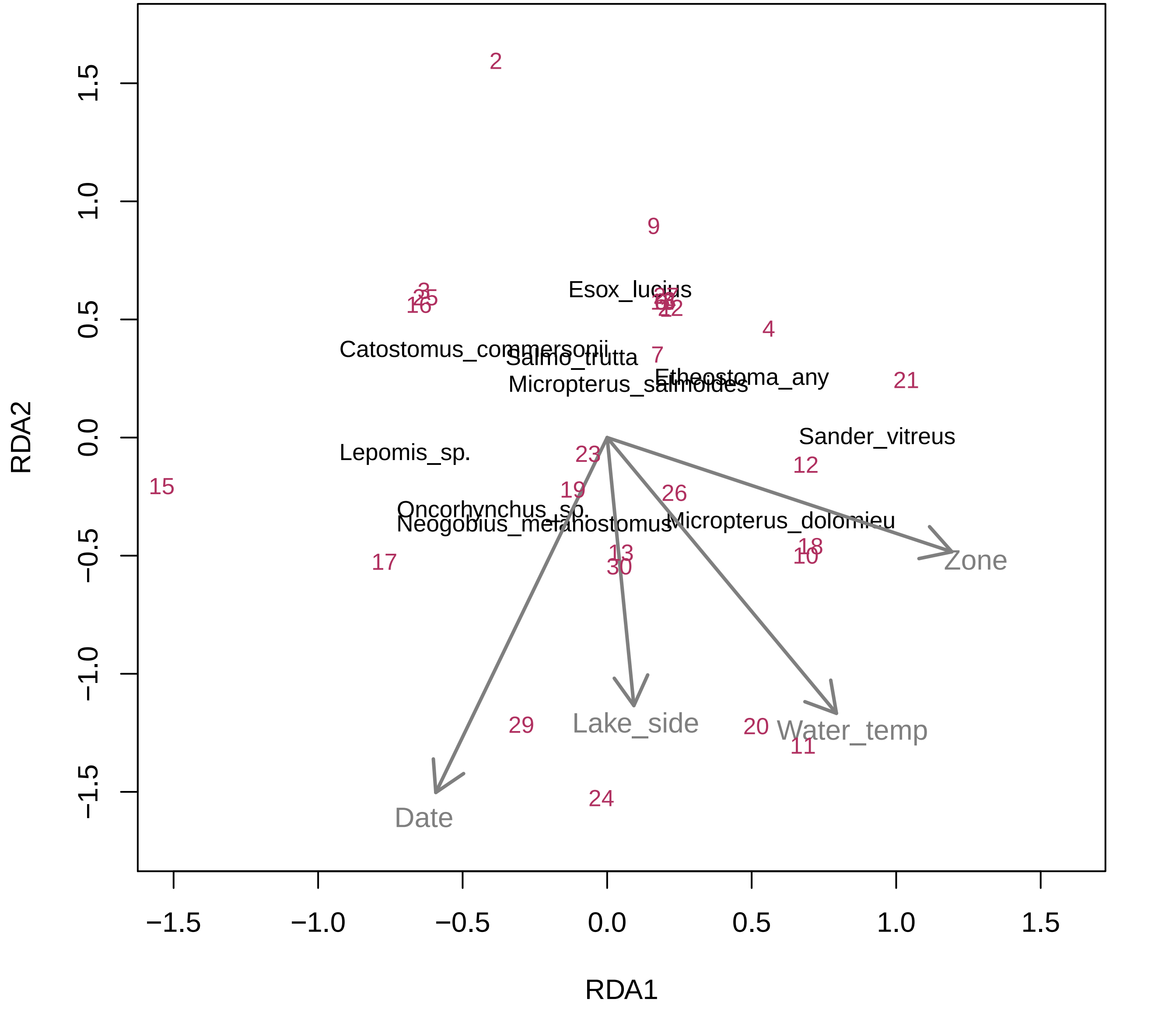
